# HNRNPC-mediated RNA processing promotes VHL-associated excessive vasculogenesis in von Hippel-Lindau syndrome revealed from patient-derived blood vessel models

**DOI:** 10.64898/2026.10.30.762950

**Authors:** Hidenori Ito, Iori Sato, Mami Matsuo-Takasaki, Tomoya Shimizu, Michiya Noguchi, Yukio Nakamura, Naoki Kubo, Daigo Takagi, Masaru Tamura, Megumu K Saito, Shunsaku Takayanagi, Nobuhito Saito, Eijiro Nakamura, Yohei Hayashi

## Abstract

Von Hippel-Lindau Syndrome (VHLS) is an inherited disorder characterized by excessive vascularization, which contributes to the development of hemangioblastomas and benign or malignant tumors, as well as renal cell carcinoma (RCC) and cysts in multiple organs. VHLS is caused by mutations in *VHL* tumor suppressor gene, which encodes a component of an E3 ubiquitin ligase complex. Although therapeutic agents targeting hypoxia-inducible factor 2 alpha (HIF2A), which is stabilized by VHL loss and promotes tumor angiogenesis, have been approved for clinical use, their therapeutic response remains limited in RCC patients associated with VHLS. Therefore, there is an unmet need for the development of alternative therapeutic approaches independent of HIF2A. Here, we establish a human iPSCs-based model of VHLS-associated excessive vasculogenesis using induced pluripotent stem cells (iPSCs) derived from VHLS patients and VHL-edited isogenic iPSC lines. Vascular cells (VCs) differentiated from these VHLS-specific iPSCs exhibited excessive blood vessel formation compared to healthy donor-derived VCs, leading to vascular lesion formation characterized by abundant blood vessels and abnormal cellular growth *in vivo*. Mechanistically, RNA-binding proteins (RBPs) were preferentially stabilized in the patient-VCs. Among these RBPs, the stabilization of heterogeneous nuclear ribonucleoprotein C1/C2 (HNRNPC), which is a substrate for pVHL-mediated protein degradation independent of HIF2A expression, was critical for the dysregulated vascularization caused by *VHL* inactivation in this model. These findings provide an iPSCs-derived vascular platform to elucidate VHLS mechanisms and identify HNRNPC as a potential therapeutic target for VHL-driven vascular lesion formations through an HIF2A-independent mechanism.

## Introduction

Von Hippel-Lindau Syndrome (VHLS) (OMIM #193300) is an autosomal dominant inherited disorder and occurs in approximately 1 in 36,000 live births^1–3^. VHLS is characterized by benign tumors, cysts, and hemangioblastomas composed of excessive blood vessels and neoplastic stromal cells in multiple organs, such as the brain, retina, and spinal cord^3,4^. Also, VHLS patients frequently develop malignant tumors including renal cell carcinoma (RCC) and phaeochromocytomas^5^. VHLS patients carry a heterozygous mutation of *VHL* tumor suppressor gene located on chromosome 3p25-26. Notably, somatic mutations of VHL are a major hallmark in RCC and affect millions of people worldwide^6,7^. The inactivation of VHL protein (pVHL), which is an E3 ubiquitin ligase, fails to regulate ubiquitination of target proteins and to degrade them through proteasome-mediated pathways. Among them, hypoxia-inducible factor 2 alpha (HIF2A) is an important factor in controlling gene expression of angiocrines (VEGF, PDGF, and TGFα)^8,9^. The protein stabilization of HIF2A promotes tumor angiogenesis, the process of new blood vessels forming from existing blood vessels, to maintain continuous proliferation in RCC cells^1^. However, there are limited therapeutic responses to the HIF2A-specific inhibitor belzutifan (MK-6482, previously called PT2977) developed in recent years for RCC patients associated with VHLS^10^, implying that HIF2A is just one of the drivers for the abnormal blood vessel formation in the disease, and additional targets are required to suppress the malignant progression. Thus, there is an urgent need to understand the underlying mechanism of VHLS and develop alternative therapeutic approaches independent of HIF2A.

To date, researchers have attempted to investigate the pathogenesis of VHLS-associated tumorigenesis using *VHL* gene-deficient disease models; however, a sufficient model to comprehensively recapitulate the diverse vascular and tumor-associated phenotypes of VHLS has not been established. Homozygous *VHL*-knockout (KO) mice result in embryonic lethality due to impaired placental vasculogenesis^11^, the process by which endothelial progenitor cells (EPCs) and hemangioblasts differentiate to form blood vessels. Heterozygous deletion or biallelic *VHL*-KO mice via the Cre-mediated conditional system have been associated with vascular abnormalities, including cavernous hepatic hemangiomas and focal vascular proliferative lesions, as well as fibrotic lesions in several organs^12–14^. These findings suggest species-specific differences in the vascular consequences of VHL loss. On the other hand, human immortalized cell lines derived from RCC samples, such as RCC4 and 786-O, provide the cellular and molecular alterations caused by the inactivation of the *VHL* gene in RCC^15–17^. Indeed, HIF2A-specific inhibitors were developed largely based on VHL-deficient RCC models^18–20^. However, these models do not address how VHL mutations affect human vascular cells in the context of VHLS or whether such effects depend on the genetic and cellular backgrounds of the affected cells. These limitations underscore the need for human disease models that directly recapitulate the excessive vasculogenesis associated with VHLS.

In blood vessel development, pluripotent stem cells differentiate into posterior primitive streak of the mesoderm, the hemangioblasts or hemogenic endothelium, and vascular endothelial cells (VECs) sequentially^21,22^. VECs give rise to multiple vascular subtypes including artery, vein, capillary, and lymphatic ECs, with unique cellular processes such as protein trafficking, protease/angiocrine secretion, migration, differentiation, and proliferation^23–26^. Several groups have generated blood vessel organoid (BVO) models^27–31^ containing VECs and other vascular cell types derived from pluripotent stem cells and/or xenograft mouse models^31,32^ transplanted with BVOs to promote blood vessel formation for various tissue development and recapitulate the pathogenesis of related diseases. In addition, tumor-associated VECs have been shown to acquire cellular and molecular alterations to enhance the sprouting activity of blood vessels, leading to excessive vasculogenesis/angiogenesis in human non-small cell lung cancer (NSCLC)^33^. However, how VHL mutations directly alter the behavior of human vascular cells (VCs) and drive dysregulated vascular development in VHLS remains poorly understood.

Human induced pluripotent stem cells (iPSCs) are a promising bioresource to generate human VCs, including endothelial cells, pericytes, and smooth muscle cells for disease modeling of vasculopathy^27,31,34^. In this study, we established a patient-derived human iPSC-model of VHLS-associated excessive vasculogenesis using iPSCs derived from VHLS patients and gene-edited isogenic controls carrying heterozygous inactivation of the *VHL* gene. VCs derived from these iPSCs exhibited excessive vasculogenesis and formed vascular lesions characterized by abundant human-derived blood vessels and abnormal cellular growth in vivo owing to pVHL-driven cellular and molecular alterations in the VCs. Furthermore, we identified the heterogeneous nuclear ribonucleoprotein C1/C2 (HNRNPC) as a substrate of pVHL for proteasome-mediated degradation and found that its protein stabilization caused by *VHL* inactivation was required for excessive vasculogenesis and the induction of the oncogenic molecular signatures partly distinct from HIF2A. Our findings identify HNRNPC as a potential therapeutic target for VHL-associated vascular abnormalities and provide a human patient-derived model for investigating HIF2A-independent mechanisms underlying the vascular manifestations of VHLS.

## Results

### Development of VHLS-specific iPSCs-derived vascular models using patient-derived and gene-edited iPSCs *in vitro*

We first aimed to construct VHLS-specific iPSCs-derived vascular models. To address this point, we established and characterized patient-derived iPSCs from three VHLS patients (VHLS-iPSCs). These VHLS-iPSCs were generated from peripheral blood mononuclear cells (PBMCs) of each VHLS patient with nonintegrating episomal DNA vectors carrying reprogramming factors (*OCT3/4*, *SOX2*, *KLF4*, *MYCL1*, *LIN28*, *mP53DD*, and *EBNA1*) under feeder-free culture conditions^35,36^. To test whether these VHLS-iPSCs removed transgene *EBNA1* in episomal vectors after the generation, we performed genomic quantitative PCR for 18 iPSC clones from three VHLS patients. Then, we selected a clone from each patient (HPS2574, HPS2583, and HPS2592) with little or no copy numbers of *EBNA1* remaining in the genomic DNA (Extended Data Fig. 1a). Selected clonal VHLS-iPSCs formed typical human embryonic stem cell (hESC)-like morphology and expressed self-renewal markers, NANOG, OCT3/4, SSEA-4, and TRA-1-60 (Extended Data Fig. 1b-d). Pluripotent potentials to differentiate into three germ layers in these VHLS-iPSC clones was confirmed through *in vitro* embryoid body formations and *in vivo* teratoma formations (Extended Data Fig. 1e and f). These clones held normal karyotypes without any noticeable CNV abnormality revealed in comparative genome hybridization (CGH) / copy number variation (CNV) microarray analysis (Extended Data Fig. 1g). In addition, these iPSC lines were confirmed to be mycoplasma-negative by indirect DNA fluorescent staining and nested-PCR (Extended Data Fig. 1h). Then, the identity of these lines was verified with short tandem repeat (STR) analysis.

To identify mutations in *VHL* gene, genomic DNAs from these clones were used for PCR sequencing analysis. HPS2574 carried c.485 G > A (p.C162Y); HPS2583 carried c.414 A > G (abnormal splicing resulting in deletion exon2); HPS2592 carried c.481 C > T (p.R161STOP). These mutations were heterozygous and identical to the sequence in the reported VHLS patients^37,38^ (Extended Data Fig. 2a-c). As the mRNA expressions of Exon 2-3 in *VHL* gene were down-regulated or splicing abnormal in these patient samples (Extended Data Fig. 2d-f), pVHL expression also decreased in these VHLS-iPSCs compared to healthy donors-derived iPSCs (Healthy-iPSCs) (Extended Data Fig. 2g and h), implying that these mutations in *VHL* gene contribute to the mRNA stability or its splicing. Together, these VHLS-iPSCs maintained self-renewal, pluripotency, genomic integrity, and heterozygous mutations in the *VHL* gene.

To develop human VHLS-associated vascular models *in vitro*, we differentiated these VHLS-iPSCs and Healthy-iPSCs, which were matched in race and sex to the VHLS-iPSCs, into VCs and analyzed the cellular phenotypes in monolayer culture and the generation of three-dimensional vascular network models (Fig. 1a). Modified from previous studies^31,39–42^, we initially established a robust protocol to generate VCs from these iPSCs via mesoderm by activating BMP and WNT signaling pathways first with 50 ng/mL BMP4 and 8 µM CHIR99021 with a transient DMSO treatment, followed by activating VEGF and FGF signaling pathways with 50 ng/mL VEGF-A and 50 ng/mL basic FGF together with inhibition of the TGFβ signaling pathway with 10 µM SB431542 (Fig. 1a). These differentiated cells from VHLS-iPSCs and Healthy-iPSCs expressed VECs surface markers (i.e., CD31 and CD144) on day11 revealed with immunocytochemistry and flow cytometric analysis (Fig. 1b-c, Extended Data Fig. 3a and b). Further experiments with bulk RNA sequence analysis and quantitative RT-PCR analysis showed that angiogenesis-related genes including *PECAM1*, *CDH5*, *ACTA2*, *HIF1*, *EPAS1(HIF2*A), and *DLL4* were significantly enriched in both VHLS-VCs and healthy-VCs compared to those of undifferentiated iPSCs (Extended Data Fig. 3c-e). In contrast, self-renewal markers such as *NANOG*, *POU5F1*, and *SOX2* were markedly suppressed in these VCs, showing that this protocol effectively works for VCs induction from these iPSCs. To compare cellular phenotypes between VHLS-VCs and Healthy-VCs, we investigated migration and growth abilities of VC*s,* which were maintained in Endothelial Growth Medium 2 (EGM2). Importantly, VHLS-VCs consistently showed higher migration and proliferation abilities compared to Healthy-VCs, despite comparable proliferative ability among undifferentiated iPSC states (Fig. 1d-f and Extended Data Fig. 3f).

**Fig. 1.**
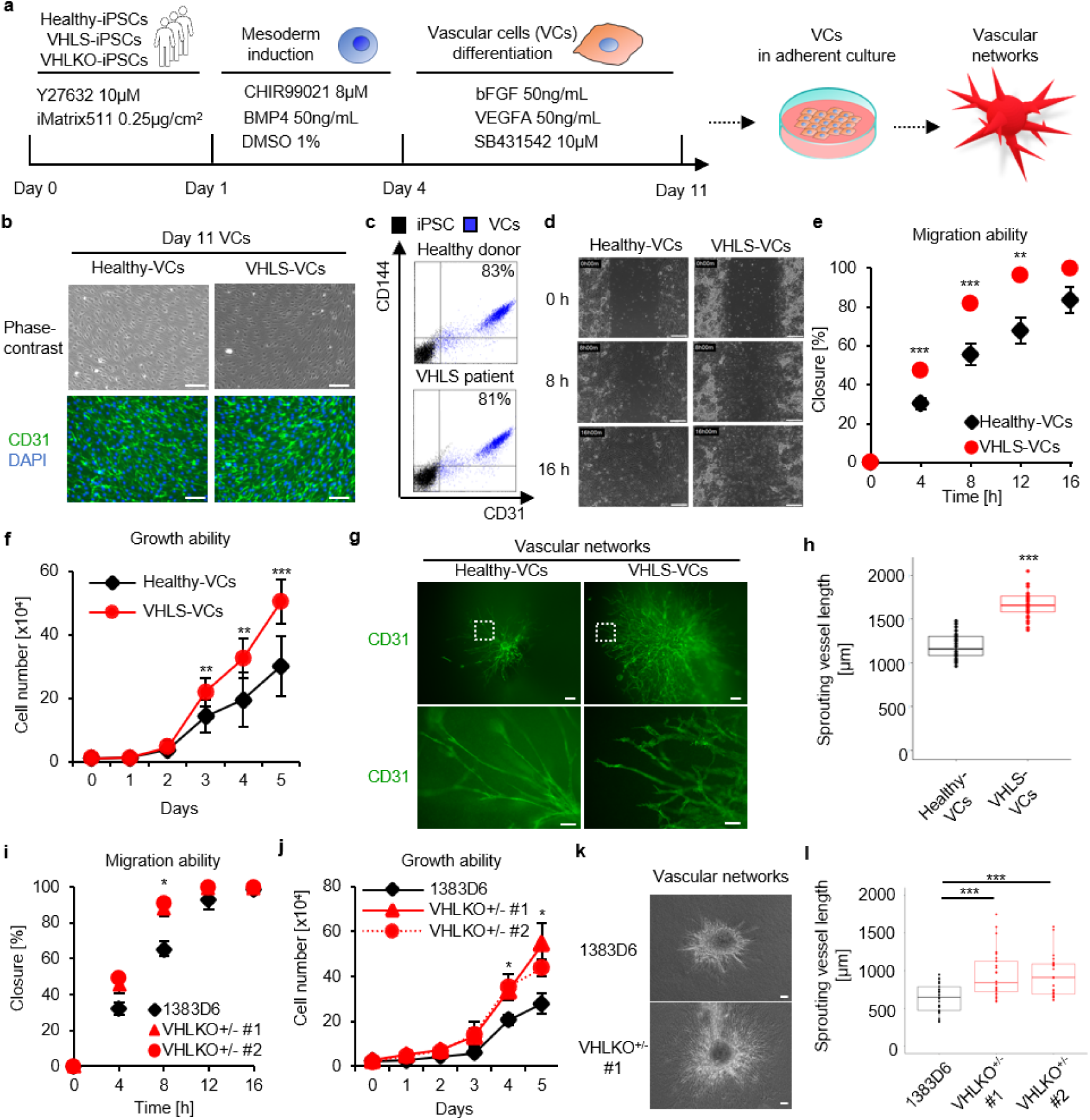
Development of VHLS-vascular models using VHLS-iPSCs in vitro a, Schematics of the VCs differentiation protocol for 11 days from Healthy-iPSCs and VHLS-ÌPSCs. The VCs were maintained and used to generate vascular networks, b, Representative phase-contrast and immunocytochemistry images of CD31^+^ VCs (green) differentiated from Healthy-iPSCs and VHLS-iPSCs on day 11. Nucleus was stained with DAPI (blue). Scale bars, 200 μm. c. Flow cytometric analysis for detecting vascular endothelial cell markers CD31 and CD144 in the VCs. d, Time-lapse images of VCs on wound healing assay at 0, 8, and 16 hours after scratch. Scale bars, 200 μm. e. Percent closure, determined at consistent locations, assessed for Healthy-VCs and VHLS-VCs at 0,4,8,12, and 16 hours after scratch (Healthy-VCs n=9, VHLS-VCs n=9). f, Cell numbers were counted for growth curves in the VCs (Healthy-VCs n=9, VHLS-VCs n=9). g, Immunocytochemistry images of CD31^+^ blood vessels (green) sprouting from the central core of VC aggregate in the Healthy-VCs and VHLS-VCs derived vascular networks. Scale bars, 200 μm. h, Sprouting vessel lengths from the central core of VC aggregate in both vascular networks (Healthy-VCs n=26, VHLS-VCs n=27). I, Percent closure measured for 1383D6 WT-VCs and VHLKO^+/-^-VCs on wound healing assay at 0,4, 8, 12, and 16 hours after scratch. Data are shown as mean = SEM (n=3). j, Cell numbers were counted for growth curves in 1383D6 WT-VCs and VHLKO^+/-^-VCs. Data are shown as mean ± SEM (n=3). k. Phase images of vascular networks from 1383D6 WT-VCs and VHLKO^+/-^-VCs. Scale bars, 200 μm. I, Sprouting vessel lengths from the central core of vascular networks (1383D6 WT-VCs n=29, VHLKO^+/-^ #1-VCs n=28, VHLKO^+/-^ #2-VCs n=27). P-values were determined by one-way ANOVA Tukey s test (*P < 0.05, **P < 0.01, ***P < 0.001).

To analyze the sprouting activity of blood vessels, we next established the vascular network model in hydrogel using these VCs by modifying previous protocols^27,28,31,43^. VCs were aggregated in a floating culture condition for 4 days, then transferred to a collagen drop on a culture dish, and kept in three-dimensional culture conditions for 6 days (Extended Data Fig. 4a and b). These vascular networks were 2-4 mm in size with almost no dead cells (Extended Data Fig. 4c and 4d), and sprouted CD31^+^ blood vessels from a central aggregate (Fig. 1g). Some of the sprouting blood vessels in vascular networks were covered with NG2-positive pericytes and α-SMA-positive smooth muscle cells (Extended Data Fig. 4e and f), and retained fluorescence-labeled dextran in the 3D structures (Extended Data Fig. 4g and h). Interestingly, VHLS-VCs-derived vascular networks significantly elongated sprouting vessel lengths more than Healthy-VCs-derived vascular networks (Fig. 1g and h). To check whether the sprouting activities of blood vessels in the vascular networks were dependent on pVHL expression, we established gene-edited isogenic controls carrying heterozygous inactivation of the *VHL* gene (VHLKO^+/-^ #1-iPSCs and VHLKO^+/-^ #2-iPSCs) from Healthy-iPSCs (1383D6 line) by using CRISPR-Cas9 system (Extended Data Fig. 5a and b). VHLKO^+/-^ #1-iPSCs and VHLKO^+/-^ #2-iPSCs were differentiated into VCs (Extended Data Fig. 5c) and used for the generation of vascular networks. VHLKO^+/-^-VCs showed higher migratory and proliferative abilities than their original Healthy-iPSCs, contributing to enhancing sprouting blood vessels in the vascular networks (Fig. 1i-l). These results indicate that the regulation of this sprouting activity on blood vessel formations was dependent on pVHL expression. Additionally, we generated VHLS-iPSCs (HPS2574, HPS2583, and HPS2592) expressing lentivirus vectors encoding *GFP* or *VHL* genes and confirmed the VC-differentiation (Extended Data Fig. 6a-c). Although we did not observe any differences in the proliferation ability between the Healthy-iPSCs and VHLS-iPSCs expressing the *GFP* and *VHL*, the *VHL*-overexpressed VHLS-VCs specifically suppressed their growth compared to the GFP-expressed VHLS-VCs (Extended Data Fig. 6e and f). Indeed, the blood vessel lengths in the vascular networks from the *VHL*-overexpressed VHLS-VCs were significantly shorter than those of the GFP-expressed VHLS-VCs (Extended Data Fig. 6g and h). Together, the promotion effect on vasculogenesis in VHLS-VCs is reversibly dependent on pVHL expression levels. Moreover, we examined whether inactivation of the second VHL allele occurring in most VHL-associated tumors accelerated vasculogenesis in this model. Since homozygous *VHL* gene knockout iPSC clones were not obtained by introducing multiple gRNAs targeting the *VHL* gene into Healthy-iPSCs or VHLS-iPSCs and analyzing 57 clones (Data is not shown), we constructed a system where introducing dicer-substrate small interfering RNA (DsiRNA) targeting the *VHL* gene into VCs derived from the patient line (HPS2574) resulted in a significantly lower expression level compared to negative control (NC) DsiRNA-transfected conditions until 10 days post-transfection (Extended Data Fig. 7a). We investigated the growth ability, migration ability, and sprouting activity in the vascular cells from the patient. There were no discernible differences in the oncogenic potentials between NC and VHL-targeted DsiRNA conditions (Extended Data Fig. 7b-f). Given that *VHL*-KO mice carrying heterozygous mutations form cavernous hepatic hemangiomas and focal vascular proliferative lesions^12^, these results suggest that the patient-derived excessive vasculogenesis observed in this model does not necessarily require complete loss of VHL gene.

To further examine whether the rapid induction of vasculogenesis in VHLS-VCs was associated with a canonical pathway, we treated VHLS-VCs and Healthy-VCs with SU5402, which is a pharmacological inhibitor for vascular endothelial growth factor receptor (VEGFR) and fibroblast growth factor receptor (FGFR)^44^, and belzutifan (PT2977), which is a HIF2A specific inhibitor^45,46^. SU5402 treatment significantly inhibited not only the sprouting activity of blood vessels in vascular networks but also cell growth in monolayer cultures derived from both lines in a dose-dependent manner (Extended Data Fig. 8a-d). In contrast, VCs treated with PT2977 specifically ceased to sprout blood vessels in the three-dimensional culture but did not affect proliferation in the two-dimensional culture (Extended Data Fig. 8e-h), indicating that three-dimensional culture may create a pseudo-hypoxic condition inside the vascular networks, leading to induction of the HIF2A expression in response to PT2977 treatment^47^. Notably, both chemicals efficiently increased the percentage of vascular networks without sprouting blood vessels; however, the half of VHLS-derived vascular networks remaining the blood vessels were resistant to the inhibitory effects of PT2977 (Extended Data Fig. 8i and 8j), possibly due to HIF2A-independent mechanisms to promote the vasculogenesis in the VHLS-VCs.

### Development of xenograft vascular mouse models using patient-derived iPSCs *in vivo*

To further examine whether the pVHL-driven abnormal blood vessel formation in VHLS-VCs could lead to excessive vasculogenesis *in vivo*, we established a xenograft vascular mouse model. For monitoring the proliferation rate of live human cells transplanted in a mouse, Venus-Akaluc fusion genes, a highly luminescent luciferase allowing noninvasive visualization inside animals^48^, were transduced into VHLS-iPSCs and Healthy-iPSCs using Piggybac transposon plasmid vectors. After antibiotic selection, the Venus-Akaluc-positive iPSCs with comparable intensity of Akaluc luminescence were further purified using fluorescence-activated cell sorting (FACS) (Extended Data Fig. 9a-c). We confirmed that these established cells maintained the differentiation potential for three-germ layers and vascular cells, and were unaffected by the overexpression (Extended Data Fig. 9d-g). In addition, since the Venus-Akaluc-positive iPSCs showed a strong correlation between tumor weight and Akaluc intensity *in vivo* teratoma formations just before the sacrifice at 6 weeks after the transplantation (Extended Data Fig. 10), the intensity of Akaluc luminescence can be a faithful quantitative indicator for transplanted human cells. We then differentiated these Venus-Akaluc-positive iPSCs into VCs and transplanted these cells at specific cell number under the kidney capsule of immunodeficient NOD.Cg-Prkdc^scid^Il2rg^tm1Wjl^/SzJ (NSG) mice to form blood vessels *in vivo* (Fig. 2a). The transplanted Venus-Akaluc-positive VCs in mice were imaged with Akalumine substrate treatment at 1, 5, 8, and 12 weeks after the transplantation with an *in vivo* imaging system (IVIS), and then sacrificed for histological analysis in the kidney sections (Fig. 2a). The VHLS-VCs rapidly proliferated and enhanced the signals more than 5-fold over the Healthy-VCs at 12 weeks while transplanted Healthy-VCs slightly increased Akaluc intensity in mice from 1 to 12 weeks (Fig. 2b and c). In addition to this high proliferative rate, we observed highly vascularized ectopic tissues including Venus-Akaluc-positive cells around the surface of mouse kidneys implanted with the VHLS-VCs, but not with the Healthy-VCs (Fig. 2d). Notably, Hematoxylin-Eosin (HE) and immunohistochemistry (IHC) staining showed that Venus-Akaluc-positive VHLS-VCs also expressed human CD31 and formed abundant vascular lumens with Venus-Akaluc-negative red blood cells and penetrated the mouse kidney, suggesting that the human-derived blood vessels functionally connect to host blood vessels (Fig. 2e and f). Moreover, Venus-Akaluc-positive VHLS-VCs also constructed hCD31^+^inhibin-alpha^-^ structures characterized by clear foamy cytoplasm containing prominent vacuoles, although the immunophenotypic characteristics of these structures did not fully correspond to the canonical neoplastic stromal cells in the VHLS-associated hemangioblastomas^49–52^ (Fig. 2f and Extended Data Fig. 11). Cyst formation in ciliopathy or RCC is often associated with dysregulation of primary cilia formation and function in the cells^53,54^. In a previous report, inactivation of pVHL directly impaired primary cilium formation in mouse kidney cells^55^. Correlated with these findings, VHLKO^+/-^ significantly suppressed the percentage of cells with ARL13B-positive primary cilia in Healthy-iPSCs (Extended Data Fig. 12a-b). Therefore, our human disease models phenotypically recapitulate excessive vasculogenesis dependent on pVHL expression, leading to the formation of VHL-associated vascular lesions.

**Fig. 2.**
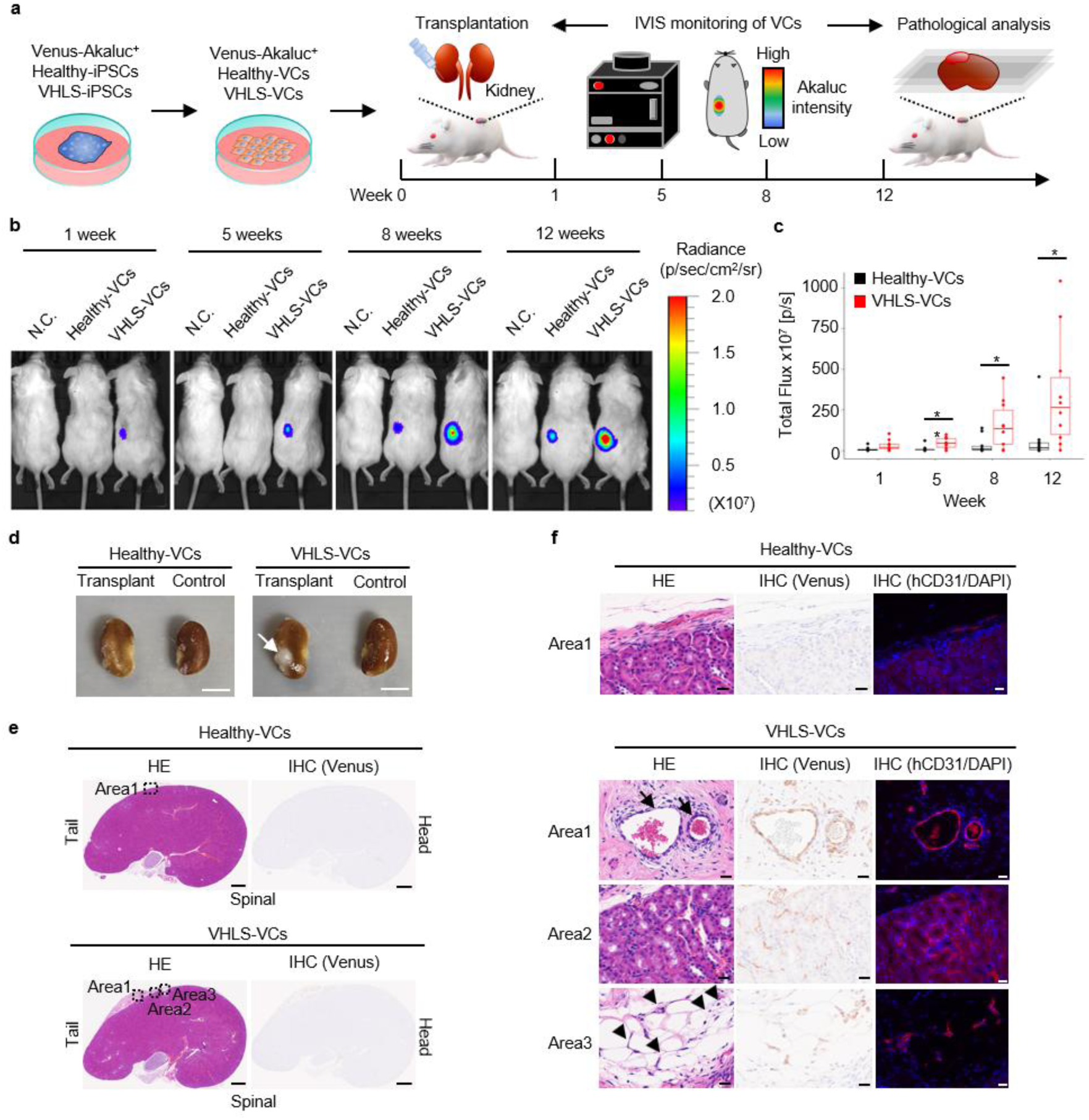
Development of xenograft vascular mouse models using VHLS-iPSCs in vivo a, Schematics of monitoring Venus-Akaluc^+^ VCs transplanted under the kidney capsule of immunodeficient mice. Healthy-VCs or VHLS-VCs expressing Venus-Akaluc fusion genes were differentiated from these IPSCs and expanded. 1.0×10^6^ cells of these VCs were resuspended in 50% geltrex and injected into a kidney. The Venus-Akaluc^+^ VCs were monitored with in vivo imaging system (IVIS) at 1,5, 8, and 12 weeks after the transplantation. At 12 weeks, the kidneys transplanted with Healthy-VCs and VHLS-VCs were isolated and sectioned for pathological analysis, b, Representative photograph and luminescence images of Akaluc-derived signals in the Venus-Akaluc^+^ VCs with IVIS at 1, 5, 8, and 12 weeks. Approximately 10 min before the imaging, the mice were intraperitoneally administered with 100 μL of 30 mM AkaLumine-HCI, and the images were acquired. Akaluc-intensity was indicated by radiance value (p/sec/cm^2^/sr). c, Total flux (p/s) was calculated from the Akaluc-intensity with IVIS (Healthy-VCs n=10, VHLS-VCs n=10). d, Macroscopic images of the control kidneys and transplanted kidneys with Healthy-VCs and VHLS-VCs at 12 weeks after the transplantation. Representative ectopic tissue was indicated by arrows. Scale bars, 5 mm. e-f, Kidney sections transplanted with Healthy-VCs and VHLS-VCs were analyzed by hematoxylin-eosin staining (HE) and immunohistochemistry (IHC) to detect Venus^+^ and hCD31^+^ human cells. Venus^+^ and hCD31^+^ blood vessels were indicated by arrows. Venus^+^ stromal cells were indicated by arrow heads. Scale bars, 1mm (whole kidney), 25 μm (Areal, Area2, and Area3). *P* values were calculated with a two-tailed student’s t-test (\**P* < 0.05). Data are shown as mean ±SEM. Experiments were repeated independently 2 times with similar results.

### VHLS-VCs stabilized RNA-binding proteins (RBPs) with oncogenic molecular alterations for vasculogenesis in VHLS

To elucidate molecular mechanisms underlying excessive vasculogenesis in VHLS patients, we performed multiomic analyses with RNA-seq transcriptome and mass spectrometry proteome profiling of VHLS-VCs and Healthy-VCs (Extended Data Fig. 13a). From the RNA-seq analysis, the differential expression analysis identified 56 up-regulated genes including mitogen-activated protein kinase (MAPK) signaling, which is a hallmark of cancer development^56^, such as *MYC*, *MAP3K14*, *DUSP4*, *HSPA2*, *NGF*, *GFRA3*, and *PRKG2* genes in the VHLS-VCs (|Log_2_ Fold change| > 1, FDR p-value < 0.05) (Fig. 3a). In addition, vein endothelial cell (vein-EC) marker *NR2F2* gene^57,58^, a major regulator of cell cycle and NOTCH signaling pathways^59^, was significantly up-regulated in the VHLS-VCs, in contrast, artery endothelial cell (artery-EC) markers *APLN* and *ESM1* genes^57^ were significantly down-regulated. This expression pattern was matched with the gene expression ratio in the iPSCs-derived vein/artery or human umbilical vein endothelial cells (HUVEC) / human umbilical artery endothelial cells (HUAEC) in previous reports^57,58^ (Extended Data Fig. 13b). Furthermore, to examine whether the arteriovenous markers of VCs were specifically expressed dependent on pVHL expression, we performed RT-qPCR analysis to detect the expression of *NR2F2*, *APLN*, and *ESM1* in VHLS-VCs expressing GFP or VHL. The expression of the arterial markers *APLN* and *ESM1* was significantly increased upon VHL expression; however, no significant differences in the vascular endothelial cell markers *PECAM1* and *EPAS1*, or the vein-EC marker *NR2F2* (Extended Data Fig. 6d). Thus, the expression of arterial markers is different between patient-derived VCs and healthy controls, although the basic differentiation potentials for endothelial cells are indifferent. These results suggest that VHLS-VCs may suppress an artery-EC like phenotype during blood vessel development. Importantly, a gene set enrichment analysis (GSEA) unbiasedly applied for the entire transcriptome data revealed that multiple pathways activated in *VHL*-deficient RCC cell lines and RCC patients^7,15^, such as MYC targets, E2F targets, G2M checkpoint, and mTORC1 signaling, were also enriched in VHLS-VCs. In contrast, interferon α response signaling, a gatekeeper of RCC progression^60^, was also enriched in the Healthy-VCs (FDR q-value < 0.10) (Fig. 3b and Extended Data Fig. 13c). These data indicated that VHLS-VCs reflected similar gene expression patterns to human RCC deficient in *VHL* gene and reprogrammed their transcripts to support their rapid proliferation and migration during tumor development.

**Fig. 3.**
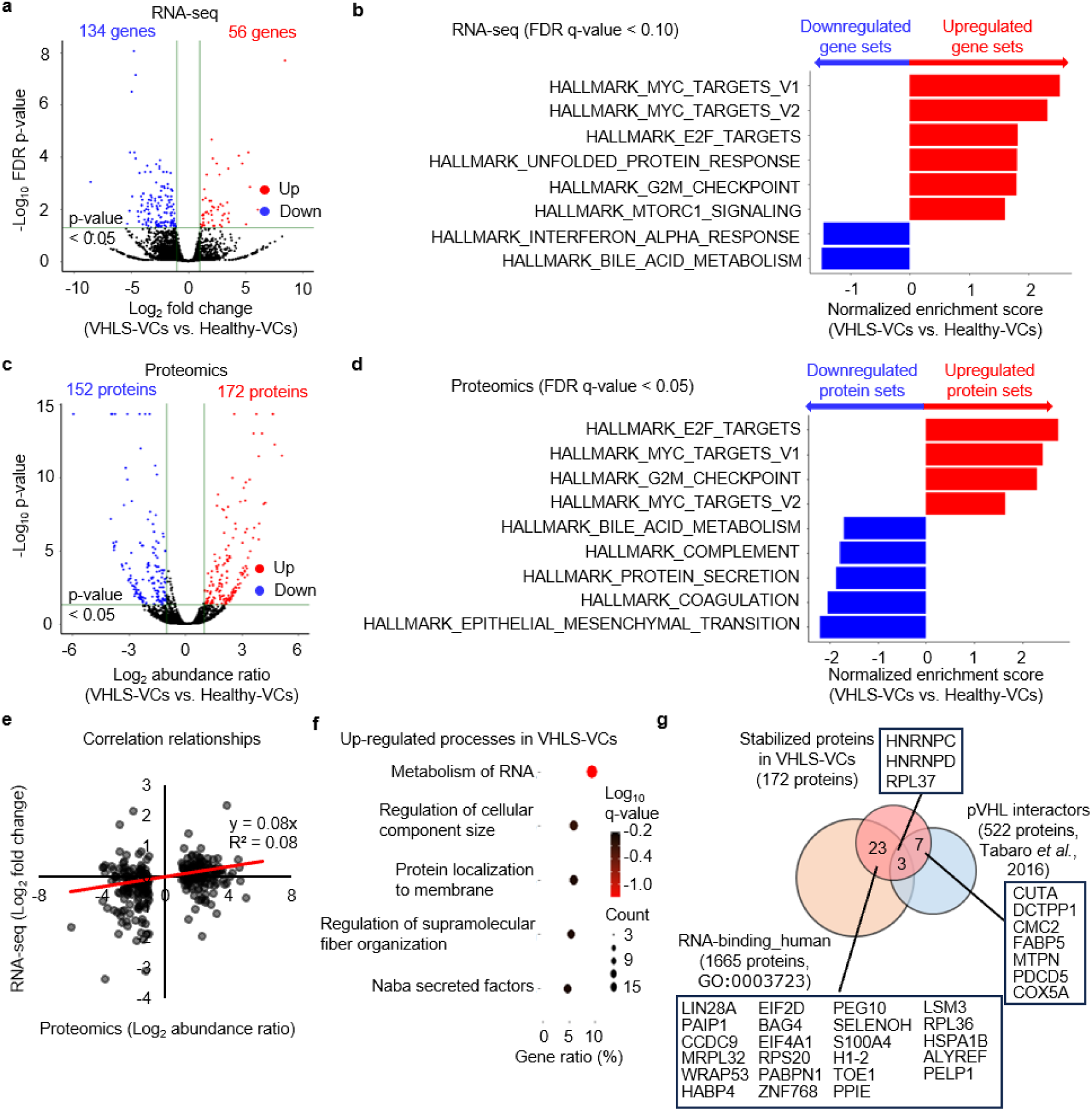
Transcriptional and proteomic profiling of tumorigenic mechanisms in VHLS-VCs a, Volcano plot showing the comparison of mRNA expression between VHLS-VCs (HPS2574, HPS2583, and HPS2592) and Healthy-VCs (1383D6, 6060A1, and 1231 A3) (baseline) based on RNA-seq data. The red color indicates up-regulated genes and the blue color indicates down-regulated genes in VHLS-VCs (|Log2 Fold change) > 1, FDR p-value < 0.05). b, Top enriched GSEA pathways in the entire transcriptome data in VHLS-VCs and Healthy-VCs (FDR q-value < 0.10). Red bars show gene data sets enriched in VHLS-VCs, and blue bars are enriched gene data sets in Healthy-VCs. c, Volcano plot showing protein abundance comparison between VHLS-VCs and Healthy-VCs (baseline) based on mass spectrometry proteomics (|Log2 Fold change) > 1, FDR p-value < 0.05). d, Top enriched GSEA pathways in the entire proteomics data in VHLS-VCs and Healthy-VCs (FDR q-value < 0.05). e, Correlation of mRNA expression and protein abundance identified in a and c. Spearmann’s correlation coefficient is depicted, f, Gene ontology analysis for the upregulated proteins in c. Dots represent term enrichment, Log_10_ q-values indicating the statistical significance of the enrichment are represented along a gradient color from black (less significant) to red (most significant). The sizes of the dots represent the count of differentially expressed proteins belonging to each term, g, Venn diagram showing the overlap among stabilized proteins in VHLS-VCs, RNA-binding proteins in human (G0:0003723), and pVHL interactors previously reported (F. Tabaro et al., 2016.). Representative proteins that fit into each category are displayed.

From the proteomic analysis, 324 proteins showed differential expressions in VHLS-VCs and Healthy-VCs (|Log_2_ Abundance ratio| > 1, p-value < 0.05) (Fig. 3c). The upregulated proteins in the VHLS-VCs included known interactors with pVHL such as RPL37, HNRNPC, HNRNPD, CUTA, DCTPP1, CMC2, FABP5, MTPN, PDCD5, and COX5A^61^ in addition to cell cycle and cell growth-related proteins. Unbiased GSEA revealed that oncogenic pathways correlated with those of transcriptional levels, such as MYC targets, E2F targets, and G2M checkpoint, were also enriched in VHLS-VCs. Conversely, coagulation signaling, associated with hemostasis step to repair vessel wall following injury, was also enriched in the Healthy-VCs (FDR q-value < 0.05) (Fig. 3d). However, no significant changes in “Hypoxia” or “Glycolysis” pathway genes related to HIF2A expression were detected in the VCs under normoxic conditions (20 % O_2_ and 5 % CO_2_) in contrast to the activation of these pathways in *VHL*-deficient RCC cell lines or RCC patients (Extended Data Fig. 13d and e). Of note, we observed poor correlations between transcript and translated protein expressions of the same gene / protein identified in the differential expression analysis (Fig. 3e). Because pVHL as a E3 ubiquitin ligase regulates the degradation of target proteins^62^, the inactivation of pVHL results in the stabilization of target proteins^9^. These stabilized proteins may regulate transcriptional alterations in the VHLS-VCs. Thus, it is understandable that the expression levels of mRNA and proteins may be not correlated well in *VHL*-deficient cells. To characterize biological processes associated with the stabilized proteins identified in the differential expression analysis (Fig. 3c), we performed ontology analysis using “Metascape”, a web-based tool for comprehensive protein list annotation^63^. Remarkably, we found that the stabilized proteins in the VHLS-VCs were mainly categorized into “Metabolism of RNA” process involved in the post-transcriptional control of RNAs such as processing, maturation, and stabilization in the cells (Fig. 3f). Indeed, the numerous stabilized proteins in the VHLS-VCs classified as RNA-binding proteins (RBPs) including three pVHL known interactors, heterogeneous nuclear ribonucleoprotein C (HNRNPC), heterogeneous nuclear ribonucleoprotein D (HNRNPD), and ribosomal protein L37 (RPL37)^61^ (Fig. 3g). Collectively, these results demonstrated that the VHLS-VCs mostly stabilized RBPs for RNA modifications as well as cell cycle and cell growth-related proteins.

### Characterization and purification of vascular cells derived from healthy donors and VHLS-patients

To characterize VCs differentiated from iPSCs deeply at single cell levels, we performed single-cell RNA-seq analysis (scRNA-seq) following the procedure shown in Fig. 4a. We integrated the datasets from Healthy-VCs and VHLS-VCs, and visualized cell transcriptomes in a reduced dimension embedding (Fig. 4b). These VCs were distinguished into 13 clusters based on their molecular signatures, with the majority of these clusters classified as VEC populations featured with *PECAM1*^+^, *KDR*^+^, or *CDH5*^+^ (Fig. 4b-e). Also, non-VEC populations constituting the other clusters comprised *PDGFRβ*^+^ and *NG2*^+^ vascular pericytes, *ACTA2*^+^ smooth muscle cells, *SULF1*^+^ and *COL4A1*^+^ fibroblasts #1, *CD45*^+^ hematopoietic cells, *ETV2*^+^ endothelial progenitor cells, *LUM*^+^ fibroblasts #2, and others (Fig. 4b-e). These results indicate that our vascular system consists of the major vascular cell types to form blood vessels^64^. To further explore cellular features in VHLS-VECs, we extracted and divided the VEC populations into Healthy-VECs and VHLS-VECs. VHLS-VECs displayed different molecular phenotypes compared to Healthy-VECs and formed the specific clusters (cluster 0, 3, 4, 6, 9) composed of more than 90% VHLS-VECs (Fig. 4f and 4g). Although all clusters from the Healthy-VECs and VHLS-VECs broadly expressed VEC markers (*PECAM1* and *KDR*) and *HNRNPC*, MYC-regulated cell cycle factors (*CCND1* and *CDK2*) were enriched in the specific clusters in VHLS-VECs (Fig. 4h), correlating with the bulk RNA-seq analysis results for the Healthy-VCs and VHLS-VCs. In contrast, the expression levels of artery-EC marker (*ESM1*) were enriched in the specific clusters in Healthy-VECs (cluster 1 and 5), while some clusters with hypoxia inducible factor (*EPAS1*) or vein-EC marker (*NR2F2*) high expression were observed in both VECs (Fig. 4h). To identify arteriovenous types in the VECs derived from the healthy donors and VHLS patients, we annotated artery-EC with *ESM1*^+^*CXCR4*^+^*SOX17*^+^ and vein-EC with *NR2F2*^+^*EPHB4*^+^. While VECs before arteriovenous specification mainly comprised the Healthy-VECs and VHLS-VECs, artery type was autonomously suppressed in the VHLS-VECs with vein type nearly identical in Healthy-VECs and VHLS-VECs (Fig. 4i and 4j), correlating with the arteriovenous marker expressions of the VHLS-VCs expressing GFP or VHL (Extended Data Fig. 6d).

**Fig. 4.**
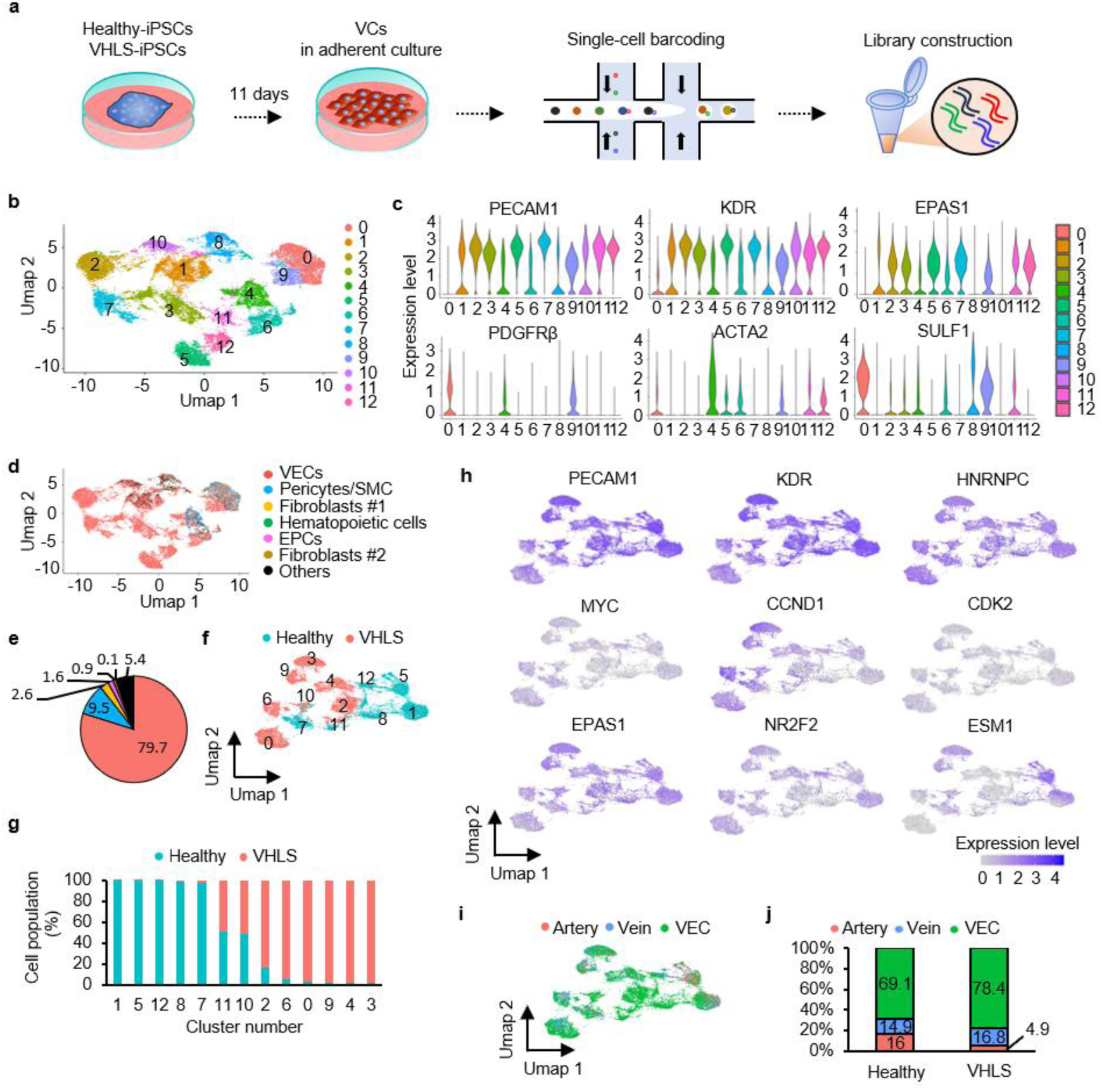
Characterization of vascular cells derived from healthy donors and VHLS-patients a, Schematic of the single-cell RNA sequencing for VCs derived from Healthy-iPSCs (1383D6 and 1231A3) and VHLS-iPSCs (HPS2574 and HPS2583) at day 11. b, A uniform manifold approximation and projection (UMAP) visualization of scRNA-seq data for all cell types in the VCs, including vascular and non-vascular types, after filtering low-quality cells and doublets, c, Violin plots showing the selective expression of key genes, including vascular endothelial cells (PECAM1 and KDR). hypoxia inducible factor (EPAS1). pericyte (PDGFRβ). smooth muscle cell (ACTA2), fibroblast (SULF1), on the 13 clusters, d, UMAP embedding of scRNA-seq data with cells colored by cell-type cluster, including VEC (PECAM1^+^, KDR^+^, orCDH5^+^), pericyte/SMC (NG2^+^, PDGFRβ^+^ or ACTA2^+^), fibroblasts #1 (SULF1^+^ and COL4A1^+^), hematopoietic cells (CD45^+^), EPC (ETV2^+^), fibroblasts #2 (LUM^+^), and others, e, Pie chart indicating the percentage of cells from each cell type, including VEC (red), pericyte/SMC (blue), fibroblast #1 (yellow), Blood (green), EPC (pink), fibroblast #2 (khaki), and others (black), f, UMAP of single-cell transcriptomes in the VEC from the healthy donors (blue) and VHLS patients (red), g, Percentage of cell population from the healthy donors (blue) and VHLS patients (red) in each cluster, h, Feature plots of representative marker genes for VEC (PECAM1 and KDR). RNA-binding protein (HNRNPC), MYC-regulated cell cycle factors (MYC, CCND1, CDK2), hypoxia inducible factor (EPAS1), vein (NR2F2), and artery (ESM1) in (f). i, UMAP of single-cell transcriptomes in VEC from arteriovenous types, including ESM1^+^CXCR4^+^ SOX17^+^ artery(red), NR2F2^+^EPHB4^+^vein (blue), and VEC(green) in (f). j, Bar chart showing the percentage of the arteriovenous types in VEC derived from the healthy donors and VHLS patients.

Regarding the vascular networks sprouting three-dimensionally from a central aggregate, we performed flow cytometry analysis to identify the cell types, and identified that 80% of them comprised CD31^+^CD144^+^ VECs and 15% comprised CD31^-^PDGFRβ^+^ pericytes (Extended Data Fig. 14a and 14b). Unexpectedly, some of CD31^+^CD144^+^ VECs also expressed PDGFRβ implying early endothelial-mesenchymal transition in the vascular networks^65^ (Extended Data Fig. 14a and 14b). Furthermore, after CD31^+^CD144^+^ VECs were sorted from VCs derived from healthy individuals (WTC11, 606A1, 1231A3) and VHLS patients (HPS2574, HPS2583, HPS2592), we compared these vessel lengths in the resulting vascular networks generated from these sorted fractions revealed a marked increase in vessel length within the patient-derived networks (Extended Data Fig. 14c and 14d). Thus, the excessive vascularization observed in patient cells should be primarily attributed to the excessive proliferation and migration of VHLS-VECs, confirmed with robust reproducibility.

### Identification of the HNRNPC protein in RBPs as a natural inhibitor for pVHL-driven vasculogenesis

We hypothesized that a critical protein in the stabilized RBPs was responsible for abnormal vasculogenesis caused by down-regulating pVHL-dependent cellular and molecular alterations in VHLS-VCs. Based on our multiome analysis and previous interactome information (Fig. 3), we selected three RBPs of HNRNPC, HNRNPD, and RPL37 as promising candidates for therapeutic targets in the abnormal vasculogenesis in VHLS-VCs. Then we attempted to establish knockout iPSC lines by introducing multiple gRNAs targeting the HNRNPC, HNRNPD, and RPL37 genes into VHLS-iPSCs; however, only heterozygous knockout lines for HNRNPC were obtained in all 60 clones (Data is not shown). Previous studies revealed that HNRNPC, HNRNPD, and RPL37 were required for mouse embryonic development^66^ or cell survival^67^ and cell cycle progression^68^, indicating these three genes were essential for early development or should be crucial for the survival and proliferation of iPS cells. Therefore, we decided to focus first on HNRNPC and applied the heterozygous knockout lines in HNRNPC as the transcriptional repression model to evaluate inhibitory effects in oncogenic phenotypes of VHLS-VCs. To examine whether HNRNPC plays a role in excessive vasculogenesis using our models, we established three iPSC clones carrying heterozygous knockout caused by a frameshift with 4 bp deletion of *HNRNPC* gene (namely, HNRNPCKO^+/-^#1-iPSCs, HNRNPCKO^+/-^#2-iPSCs and HNRNPCKO^+/-^#3-iPSCs) derived from VHLS-iPSCs using the CRISPR-Cas9 gene editing^69^ (Extended Data Fig. 15a-b). The HNRNPC proteins in HNRNPCKO^+/-^-VCs showed about half the amount of the patient wild type (WT) despite their comparable pVHL levels (Fig. 5a). In addition, HNRNPCKO^+/-^-VCs showed the similar morphology, expression patterns of VCs surface markers (CD31 and CD144) to the original VHLS-VCs, and specifically increased the expressions of vascular endothelial cell markers, PECAM1 and EPAS1, compared to those of iPSCs (Fig. 5b-5c and Extended Data Fig. 18e). Although we did not detect any differences in migration ability between original VHLS-VCs and HNRNPCKO^+/-^-VCs, the proliferation ability was much slower in HNRNPCKO^+/-^-VCs than in original VHLS-VCs (Fig. 5d and 5e). HNRNPCKO^+/-^-VCs had defective in sprouting blood vessels with 20-30% of vascular networks lacking sprouting in the three-dimensional culture compared to the original VHLS-VCs, indicating that HNRNPC plays a critical role in pVHL-driven excessive vasculogenesis (Fig. 5f and 5g). Next, we examined whether HNRNPC depletion affects the vascular lesions in the xenograft vascular mouse models. After the construction of HNRNPCKO^+/-^-iPSCs expressing Venus-Akaluc fusion genes (Extended Data Fig. 16a-d), these iPSCs were differentiated into VCs (Extended Data Fig. 16e) and implanted under the kidney capsule of immunodeficient NSG mice. Interestingly, HNRNPCKO^+/-^-VCs blocked the high engraftment and proliferation of VHLS-VCs *in vivo* detected with the intensity of Akaluc luminescence (Fig. 5h and 5i). Although almost no ectopic tissues on mouse kidneys transplanted with HNRNPCKO^+/-^-VCs were observed, some transplanted kidneys had structures containing Venus-Akaluc-positive small blood vessels and penetrated the mouse kidney (Fig. 5j and 5k). On the other hand, Venus-Akaluc^+^ cells composed of clear foamy cytoplasm with vacuoles were not observed in mouse kidneys transplanted with HNRNPCKO^+/-^-VCs (Fig. 5k). These data demonstrate that HNRNPC regulates blood vessel development induced by pVHL inactivation and its downregulation abrogates these abnormal phenotypes, possibly leading to VHL-associated vascular lesion formation.

**Fig. 5.**
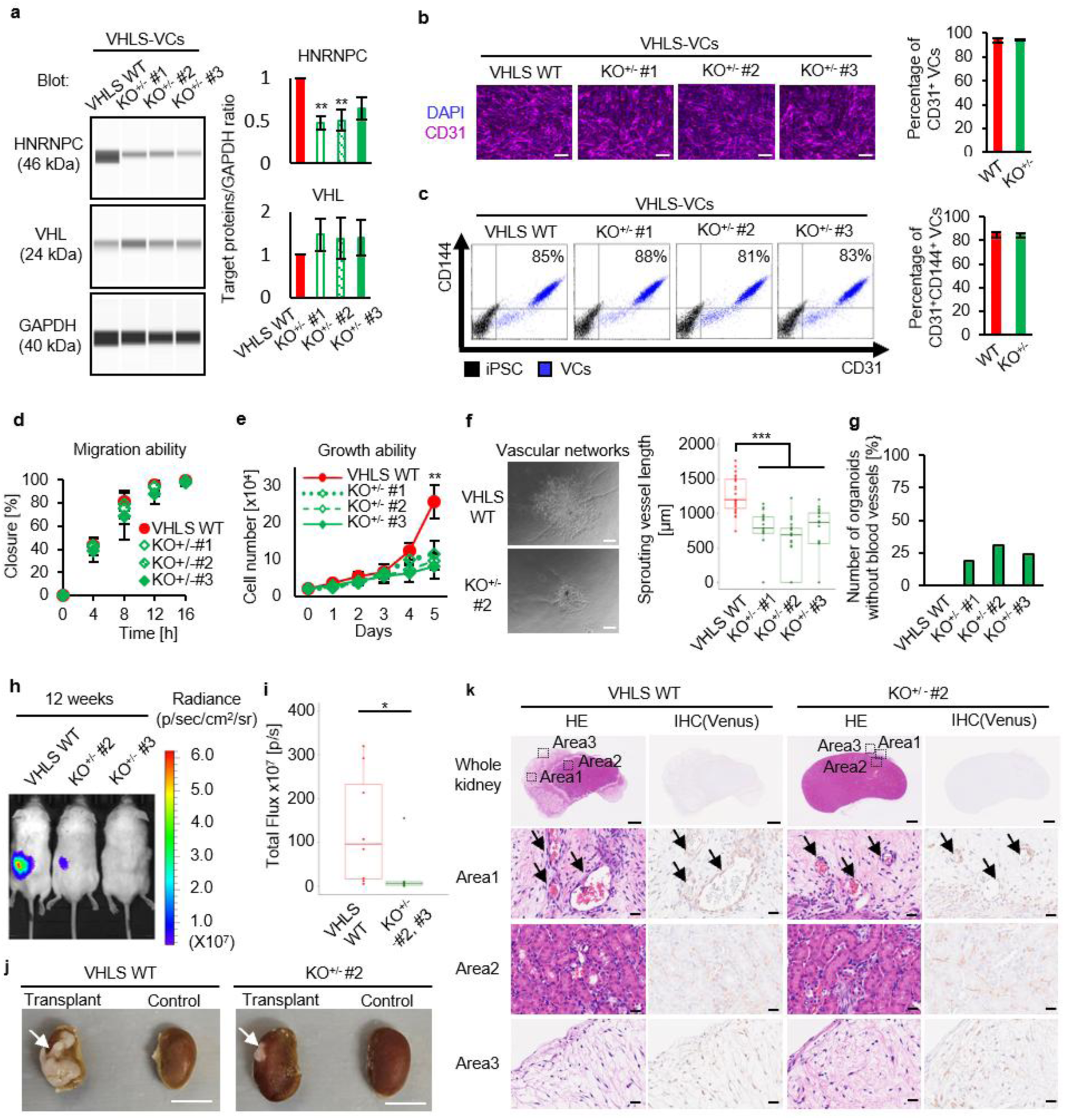
pVHL-driven vasculogenesis was inhibited by HNRNPC downregulation. a, Left, Protein expression profiles of HNRNPC in wild type (VHLS WT) and three HNRNPCKO^+/-^-VCs (HNRNPCKO^+/-^ #1, HNRNPCKO^+/-^ #2, HNRNPCKO^+/-^ #3) derived from VHLS patient line (HPS2574) by an automatic capillary western blot (WES). Right, relative amounts of HNRNPC proteins (HNRNPC/GAPDH) and VHL proteins (VHL/GAPDH) were quantified using WES Data are shown as the mean ± SEM (n = 5). P-values were determined by one-way ANOVA Dunnett’s test (**P < 0.01). b, Left. ICC images of vascular endothelial marker CD31^+^VCs (purple) differentiated from VHLS WT-IPSC and the three HNRNPCKO^+/-^-iPSCs. Nucleus were stained by DAPI (blue). Scale bars, 200 μm. Right. Percentage of the CD31^+^ VCs derived from VHLS WT and the three HNRNPCKO^+/-^lines in the DAPP cells (n=3). c, Left, Flow cytometric analysis for detecting vascular endothelial markers CD31 and CD144 in VCsand iPSCs derived from VHLS WT and the three HNRNPCKO^+/-^ lines. Right, Percentage of the CD31^+^CD144^+^ VCs derived from VHLS WT and the three HNRNPCKO^+/-^ lines in the differentiated cells on dayl 1 (n=3). d, Percent closure measured for VHLS-VCs and HNRNPCKO^+/-^ -VCs on wound healing assay at 0,4, 8, 12. and 16 hours after scratch. Data are shown as mean ± SEM (n=3). e, Cell numbers were counted for growth curves in VHLS-VCs and HNRNPCKO^+/-^-VCs. P-values were determined by one-way ANOVA Tukey’s test (\**P* < 0.01) Data are shown as mean ± SEM (n=3). f, Left, phase-contrast images of blood vessels sprouting from the central core of VC aggregate in a VHLS WT- and HNRNPCKO^+/-^ -derived vascular networks. Scale bars, 200 μm. Right. Sprouting vessel lengths from the central core of VC aggregate in these vascular networks (VHLS WTn=27, HNRNPCKO^+/-^ #1 n=21, HNRNPCKO^+/-^ #2 n=26, HNRNPCKO^+/-^#3 n=21) quantified by Imaged software. P-values were determined by one-way ANOVA Tukey s test (***P < 0.001). g, Percentages of the vascular networks without sprouting vessels in all VHLS WT and HNRNPCKO-lines derived vascular networks (green), h, Kidney capsule of VHLS-VCs and HN RNPCKO^+/-^-VCs expressing a Venus-Akaluc fusion gene monitored by I VIS at 12 weeks after the transplantation. Akaluc-intensity was indicated by radiance value (p/sec/cm^2^/sr) I, Total flux (p/s) was calculated from the Akaluc-intensity with MS LuminaXRMS series IE ver4.7 (VHLS WT-VCs n=9, HNRNPCKO^+/-^-VCs #2 and #3 n=9). P values were calculated with determined by a two-tailed student’s t-test (*P < 0.05). j, Macroscopic images of the control kidneys and transplanted kidneys with VCs of VHLS WT and HNRNPCKO^+/-^ #2 at 12 weeks after the transplantation. Representative ectopic tissue was indicated by arrows. Scale bars. 5 mm. k, Kidney sections implanted with VCs of VHLS WT and HNRNPCKO^+/-^ #2 were isolated and analyzed by hematoxylin-eosin staining (HE) (left) and immunohistochemistry (IHC) (right) to detect Venus^+^ human cells. Venus^+^ blood vessels were indicated by arrows. Scale bars. 1mm (whole kidney), 25 μm (Areal, Area2, and Area3). Experiments were repeated independently 3 times (a-g) and 2 times (h-j) with similar results.

We further investigated the correlation between HNRNPC gene expression and prognosis in VHL deficient-related tumors. Based on clinical data in patients suffering from clear cell RCC (ccRCC), the most common type of RCC, a significantly higher gene expression of HNRNPC was observed in ccRCC compared to healthy kidneys (Extended Data Fig. 17a). Furthermore, the higher expression of HNRNPC contributed to poor prognosis (Extended Data Fig. 17b), correlated with various cancer types including cervical cancer^70^, pancreatic cancer^71^, hepatocellular carcinoma (HCC)^72,73^, and lung adenocarcinoma^74^. This clinical evidence is in contrast to other key factors, such as HNRNPD, DCTPP1, MYC, EPAS1 (HIF2A) (Extended Data Fig. 17c-j). Therefore, HNRNPC can be a prognosis marker and functional therapeutic target for VHLS as well as cancers involving VHL inactivation.

### Loss of pVHL fails to degrade HNRNPC through the ubiquitin-proteasome system to reprogram transcriptional expression patterns for the tumor progression

Although HNRNPC was previously identified as pVHL interactor from an interactome analysis^75^, how pVHL regulates HNRNPC protein expression has not been clarified. Because the most well-known function of pVHL was E3 ubiquitin ligase for proteasome-mediated degradation, we investigated whether HNRNPC is a target substrate of pVHL in VCs. Co-immunoprecipitation (Co-IP) assays performed at the endogenous protein level to validate the physical interaction between pVHL and HNRNPC revealed that pVHL was directly bound to HNRNPC as well as HIF2A in Healthy-VCs (Fig. 6a). We next examined whether the inactivation of pVHL induced the stabilization of the HNRNPC protein. The protein expression of HNRNPC as well as HIF2A and HIF1A was up-regulated in VHLKO^+/-^-VCs compared to healthy-VCs (Fig. 6b and 6c). To further assess whether pVHL induces the ubiquitination of HNRNPC in a proteasome-dependent manner, we treated Healthy-VCs and VHLKO^+/-^-VCs with MG132, a specific chemical inhibitor of proteasomes, and performed an ubiquitination assay. Importantly, the inhibition of proteasomal activity with MG132 resulted in accumulation of polyubiquitinated HNRNPC in both VCs, whereas the poly-ubiquitinated HNRNPC was markedly suppressed in VHLKO^+/-^-VCs (Fig. 6d). These results indicate that loss of pVHL fails to degrade HNRNPC through the ubiquitin-proteasome system, resulting in the stabilization of HNRNPC protein. Using VCs differentiated from VHLS-iPSCs, we also performed analysis of protein-protein interactions, ubiquitination, and protein expression levels VHL-HNRNPC. Similar to the results obtained from VHLKO cell lines, we demonstrated that VHL-mediated ubiquitination of HNRNPC was suppressed and HNRNPC protein expression was stabilized in VHLS-VCs, compared to healthy controls (Extended Data Fig. 18a-d). This stabilization should result in the changes of the mRNA processing demonstrated in our transcriptome analysis.

**Fig. 6.**
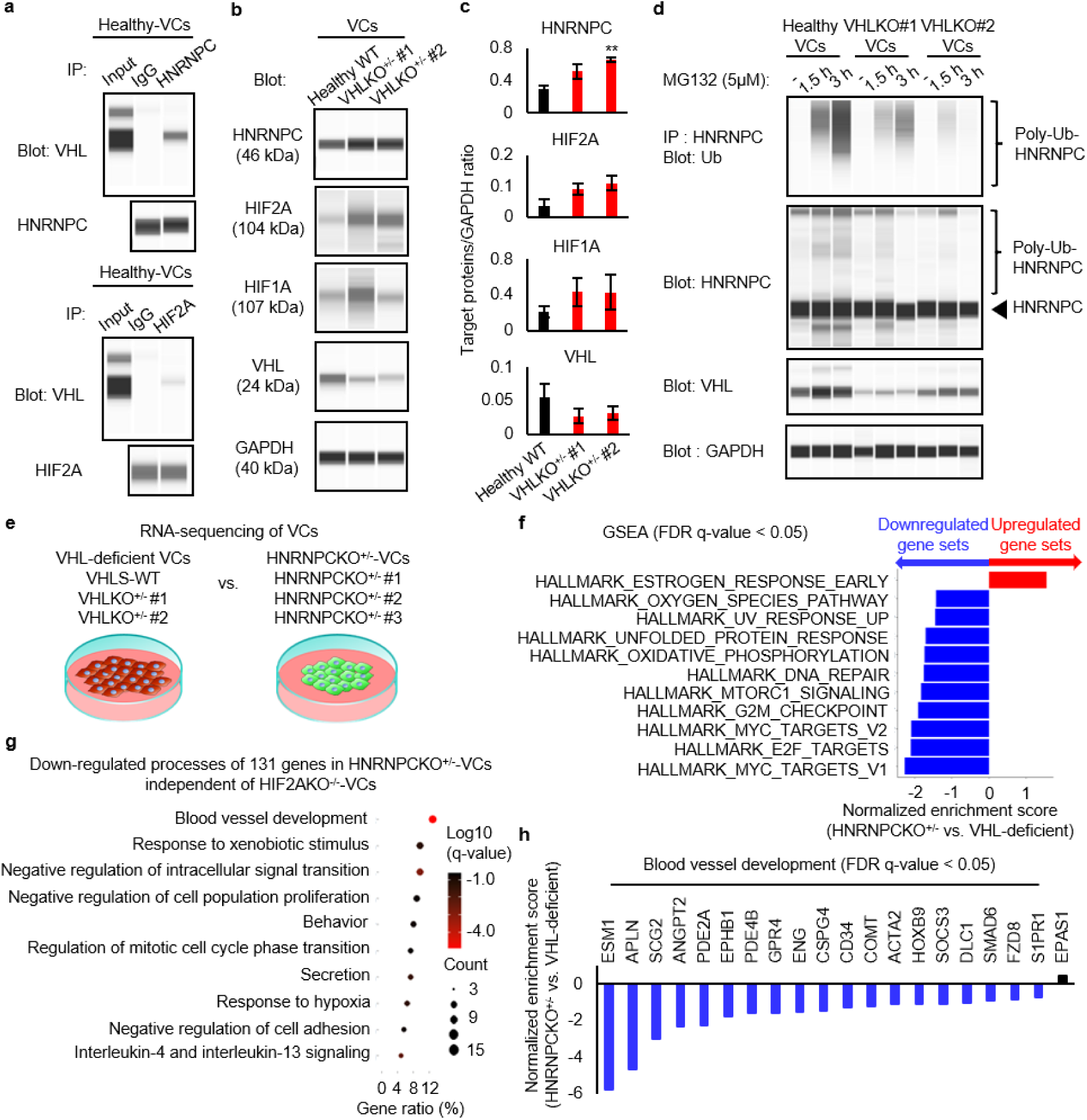
HNRNPC proteins were stabilized by pVHL inactivation through the ubiquitin-proteasome system to support transcriptional advantages for their excessive vasculogenesis independent of HIF2A expression. a, Co-immunoprecipitation (Co-IP) assays of endogenous proteins of HNRNPC and HIF2A, or control IgG were performed. For detecting HIF2A, a Healthy-VCs (1383D6 line) was temporally cultured in hypoxic conditions at 1% O_2_ environment for 6 hours just before the protein collection, b, Western blot analysis of HNRNPC, HIF2A, HIF1A, VHL, and GAPDH proteins in Healthy-VCs (1383D6) and VHLKO^+/-^-VCs (VHLKO^+/-^ #1 and VHLKO^+/-^ #2) stimulated with 1% O_2_ for 6 hours, c, Relative amounts of HNRNPC, HIF2A, HIF1A, and VHL proteins normalized by GAPDH using WES. Data are shown as the mean ± SEM (n = 3). d_s_ Ubiquitination of endogenous HNRNPC proteins in Healthy-VCs (1383D6) and VHLKO^+/-^-VCs (VHLKO^+/-^ #1 and VHLKO^+/-^ #2) treated with 5pM MG-132 for 1.5 or 3 hours. Cell lysates were analyzed by immunoprecipitation with an antibody against HNRNPC followed by immunoblotting with an antibody against ubiquitin, and by immunoblotting with antibodies against HNRNPC, VHL, and GAPDH. e, Schematic of RNA-seq transcriptome on VCs derived from VHL-deficient lines (HPS2574, VHLKO^+/-^ #1, and VHLKO^+/-^ #2) (baseline) and HNRNPCKO^+/-^ lines (HNRNPCKO^+/-^ #1, HNRNPCKOW, and HNRNPCKO^+/-^ #3). f, Top enriched GSEA pathways in the entire transcriptome data in VHL-deficient VCs and HNRNPCKO^+/-^ -VCs (FDR q-value < 0.05). Red bars show gene data sets enriched in VHL-deficient VCs, blue bars are enriched gene data sets in HNRNPCKO^+/-^-VCs. g, Gene ontology analysis for the 131 downregulated genes in HNRNPCKO÷^A^ lines, but not in HIF2AKO^+/-^lines, compared to VHL-deficient lines. Dots represent term enrichment, Log_10_ q-values indicating the statistical significance of the enrichment are represented along a gradient color from black (less significant) to red (most significant). The sizes of the dots represent the count of differentially expressed genes belonging to each term, h, Gene expression profiles associated with blood vessel development (G0:0001568) identified in g (blue) (FDR q-value < 0.05). *EPAS1* (HIF2A) expression was also indicated as a control (black).

To comprehensively understand the molecular mechanisms regulated by HNRNPC in this VHLS model, we examined the transcriptome in HNRNPCKO^+/-^-VCs compared to *VHL*-deficient VCs (i.e., VHLS-VCs, VHLKO^+/-^#1-VCs, and VHLKO^+/-^#2-VCs) using bulk RNA-sequencing analysis (Fig. 6e). GSEA was unbiasedly performed for the entire transcriptome data between HNRNPCKO^+/-^-VCs and *VHL*-deficient VCs. The oncogenic pathways activated in the VHLS-VCs or papillary RCC patients, which comprises approximately 10-15% of all RCC with high HNRNPC expression^76^, such as MYC targets, E2F targets, G2M checkpoint, and mTORC1 signaling were significantly down-regulated by HNRNPC depletion (FDR q-value < 0.05) (Fig. 6f). In contrast, genes related to estrogen response were up-regulated in HNRNPCKO^+/-^-VCs. Indeed, we have examined gene expressions of artery markers (*APLN* and *ESM1*) and the MAPK pathway (*MYC* and *DUSP4*) with RT-qPCR and identified that the expression of these genes was down-regulated in HNRNPCKO^+/-^-VCs compared to WT cells, consistent with the RNA-seq results (Extended Data Fig. 18f).

Next, to investigate the effects of high HNRNPC expression in regulating RNA splicing in VHLS-VCs, we analyzed splicing variants using our RNA-seq data. In VHLS-VCs, 62 transcripts showed significantly suppressed expression compared to healthy controls, while 21 transcripts exhibited increased expression (Extended Data Fig. 19a). These differentially-expressed transcripts contained specific transcripts for 43 genes out of the 190 genes (22%) in total (Extended Data Fig. 19b). Conversely, among the 22,346 genes without changes in expression levels between healthy controls and patients, 37 genes (0.17%) contained specific transcripts (Extended Data Fig. 19b). Thus, we think that VHLS-VCs exhibit altered selective splicing patterns compared to healthy controls related to the differential expression. Similar analysis between patients and HNRNPKO cell lines revealed that in HNRNPKO^+/-^-VCs, 35 transcripts were significantly down-regulated compared to VHLS-VCs, while 51 transcripts were up-regulated (Extended Data Fig. 19c). These differentially expressed transcripts included specific transcripts related to 21 genes out of the 575 genes showing expression changes between VHLS-VCs and HNRNPKO^+/-^-VCs (Extended Data Fig. 19d). Conversely, they included specific transcripts related to 60 genes out of the 23,225 genes showing no expression changes between VHLS-VCs and HNRNPKO^+/-^-VCs (Extended Data Fig. 19d). As an example of typical splicing variants in VHLS, we have focused om *COL12A1* encoding type XII collagen, the variant COL12A1_8 was specifically down-regulated in VHLS-VCs, whereas the longest variant COL12A1_3 was specifically upregulated in HNRNPKO^+/-^-VCs (Extended Data Fig. 19e). These results suggest that excessive HNRNPC dysregulate mRNA splicing in VHLS-VCs.

As a reference line, we also generated three HIF2A-deficient iPSCs (i.e., HIF2AKO^-/-^ #1-iPSCs, HIF2AKO^-/-^#2-iPSCs and HIF2AKO^-/-^#3-iPSCs) from VHLS-iPSCs using the same method as to generate the HNRNPCKO^+/-^ lines (Extended Data Fig. 15c-d). The differential expression analysis in the RNA transcriptomes between HNRNPCKO^+/-^-VCs and HIF2AKO^-/-^-VCs vs. *VHL*-deficient VCs (baselines) revealed 197 downregulated genes in HNRNPCKO^+/-^-VCs including 66 genes overlapped with that of HIF2AKO^-/-^-VCs compared with *VHL*-deficient VCs (|Log_2_ Fold change| > 1, FDR p-value < 0.05) (Extended Data Fig. 15e). To investigate the biological processes specifically down-regulated in HNRNPCKO^+/-^-VCs, gene ontology analysis was performed on the 131 genes, excluding 66 genes duplicated in HIF2AKO^-/-^-VCs (Extended Data Fig. 15e). We found that “Blood vessel development” as well as “Regulation of mitotic cell cycle phase transition” and “Response to hypoxia” related processes in HNRNPCKO^+/-^-VCs were noticeably suppressed despite maintaining HIF2A mRNA expression levels comparably (Fig. 6g and 6h). On the other hand, the down-regulated 299 genes in HIF2AKO^-/-^-VCs independent of HNRNPCKO^+/-^-VCs were mainly categorized into “Cell division” and “Skeletal system development” as well as “Tube morphogenesis” related processes, involving “blood vessel morphogenesis” in which the anatomical structures of blood vessels are generated and organized. (Extended Data Fig. 15f). In addition, the commonly down-regulated processes of the 66 genes in HNRNPCKO^+/-^-VCs and HIF2AKO^-/-^-VCs were mostly classified with “Cytoskeleton in muscle cells” and “Mesenchyme development”, including “cardiac cell development” or “blood vessel development” related genes, such as *ACTC1*, *NKX2-5*, *MYH11*, *COL1A1*, *FOXF1*, *HOXA1*, *TCF21*, and *EMC10* (Extended Data Fig. 15g).

These results indicate that the stabilization of HNRNPC via pVHL inactivation specifically induces transcriptional alterations related to blood vessel development and cancer cell proliferation, independent of HIF2A expression to support pVHL-driven vasculogenesis.

### HIF2A deficiency in VHLS-VCs suppresses the sprouting activity of blood vessels via cellular phenotypes distinct from HNRNPCKO^+/-^-VCs

To investigate the differences in the inhibitory effects on the excessive vasculogenesis between HNRNPCKO^+/-^-VCs and HIF2AKO^-/-^-VCs, we analyzed cellular phenotypes of HIF2AKO^-/-^-VCs. Protein expression of HIF2A in HIF2AKO^-/-^-VCs was completely lost without affecting the expression of HIF1A, HNRNPC, and pVHL (Fig. 7a), and the differentiation efficiency was similar to that of the original VHLS-VCs (Fig. 7b and 7c). No differences were observed in migration and proliferation between HIF2AKO^-/-^-VCs and the original VHLS-VCs (Fig. 7d and 7e); however, the sprouting activity of vascular networks was significantly inhibited in HIF2AKO^-/-^-VCs (Fig. 7f and 7g). These results were consistent with those of the pharmacological inhibition of HIF2A using PT2977 (Extended Data Fig. 8e-h). Next, we induced VCs from HIF2AKO^-/-^ iPSCs expressing Venus-Akaluc fusion genes (Extended Data Fig. 16a-c), and transplanted these VCs (Extended Data Fig. 16e) under kidney capsules of immunodeficient NSG mice. HIF2AKO^-/-^-VCs showed little changes in cell proliferation compared to the original VHLS-VCs *in vivo* detected with the intensity of Akaluc luminescence (Fig. 7h). Additionally, the inhibitory effects of cilium formation via pVHL inactivation in VHLS-iPSCs were not recovered by HNRNPCKO^+/-^ as well as HIF2AKO^-/-^, suggesting that HNRNPC or HIF2A were not involved in primary cilia formation (Extended Data Fig. 12c and 12d). These results indicate that the effects of HNRNPC downregulation on blood vessel development and vascular lesion formation are considerably different from those of HIF2A in this model. In summary, inactivation of pVHL stabilizes HNRNPC proteins via the ubiquitin-proteasome system, leading to activation of the oncogenic signaling, including blood vessel development independent of HIF2A expression. HNRNPC-dependent progression of VHL-associated excessive vasculogenesis contributes to the formation of vascular lesions in the human vascular model of VHLS (Fig. 7i).

**Fig. 7.**
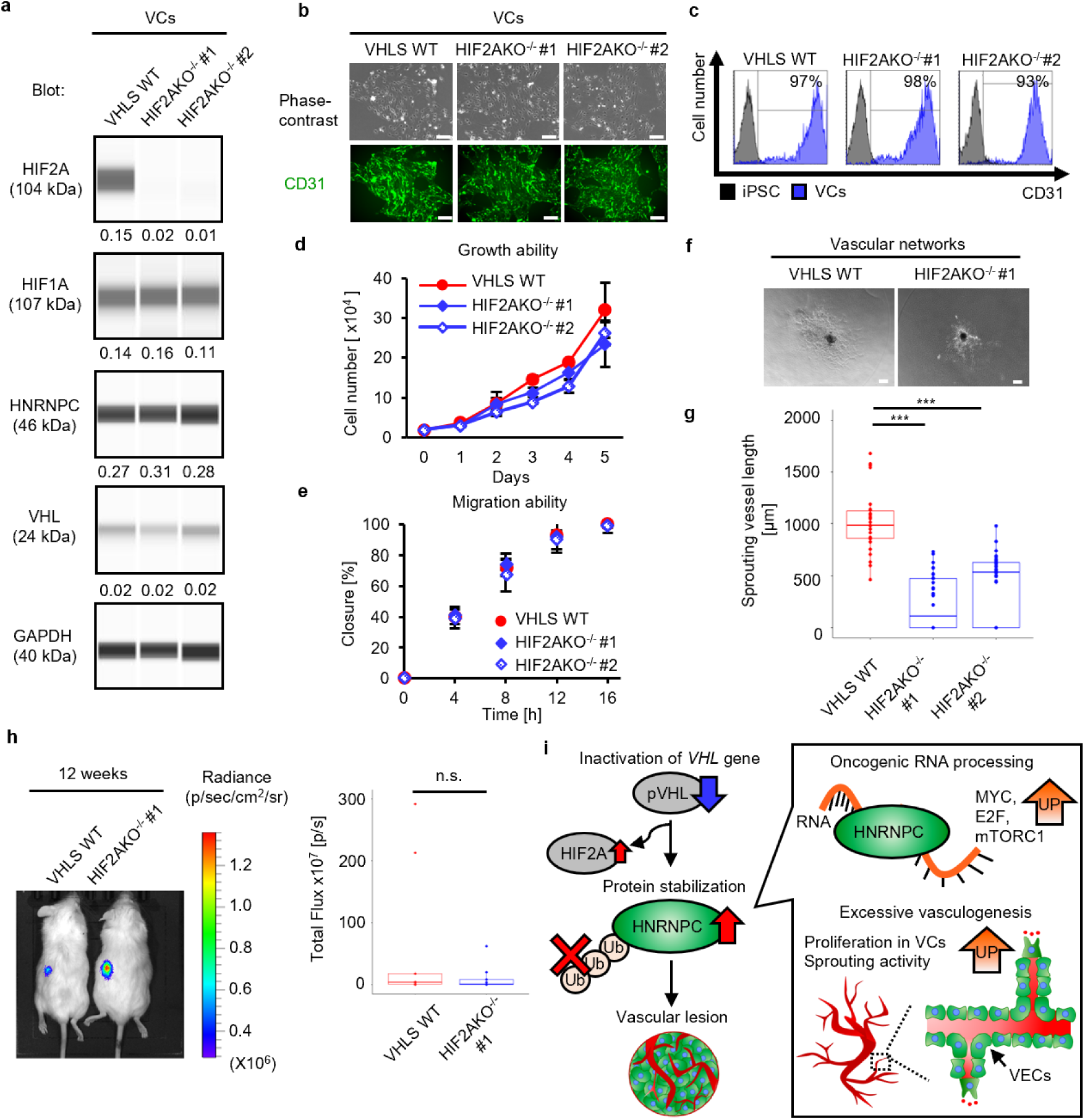
HIF2A deficiency inhibits the sprouting activity of blood vessels via cellular phenotypes distinct from HNRNPC downregulation in VHLS-VCs. a, Western blot analysis of HIF2A, HIF1A, HNRNPC, VHL, and GAPDH proteins in VHLS-VCs (HPS2574) and HIF2AKO^-/-^ - VCs (#1 and #2) stimulated with 1% O_2_ for 6 hours. Numbers below the bands indicate relative amounts of target proteins normalized by GAPDH using WES. b, Representative phase and ICC images of CD31^+^ VCs differentiated from VHLS-iPSCs and HIF2AKO^-/-^ -iPSCs. Scale bars, 200 μm. c. Flow cytometric analysis for detecting a vascular endothelial marker CD31 in the VCs. d, Cell numbers were counted for growth curves in the VCs (VHLS-VCs n=3, HIF2AKO^-/-^ #1-VCs n=3, and HIF2AKO^-/-^ #2-VCs n=3). Data are shown as mean ± SEM. e, Percent closure assessed for VHLS-VCs and HIF2AKO^-/-^-VCs on wound healing assay at 0,4,8, 12, and 16 hours after scratch (VHLS-VCs n=3, HIF2AKO^-/-^ #1-VCs n=3, and HIF2AKO^-/-^ #2-VCs n=3). Data are shown as mean ± SEM. f, Phase images of vascular networks from VHLS-VCs and HIF2AKO’^;^’-VCs. Scale bars, 200 μm. g, Sprouting vessel lengths from the central core of vascular networks (VHLS-VCs n=26, HIF2AKO^-/-^ #1-VCs n=32, and HIF2AKO^-/-^ #2-VCs n=40). P-values were determined by one-way ANOVA Tukey‘s test (***P < 0.001). h, Left, Representative photograph and luminescence images of Akaluc-derived signals in the Venus-Akaluo^+^ VCs transplanted into kidney capsule in mice. The VCs were monitored by MS at 12 weeks after the transplantation. Akaluc-intensity was indicated by radiance value (p/sec/cm^2^/sr). Right. Total flux (p/s) was calculated from the Akaluc-intensity with I VIS Lumina XRMS series in ver4.7 (VHLS-VCs n=9 and HIF2AKO^-/-^ #1-VCs n=9). i, Proposed model of the HNRNPC-dependent progression in VHLS.

## Discussions

In this study, we established a human vascular model of VHLS using patient-iPSCs and *VHL* gene-edited iPSCs. Our model recapitulates excessive vasculogenesis and vascular lesions accompanied by oncogenic molecular alterations following VHL mutation, providing a platform for investigating the mechanisms underlying the vascular manifestations of VHLS and developing potential therapeutic strategies targeting abnormal vascular cells. In this model, VHLS-VCs exhibit higher proliferative and migratory activities than healthy-VCs, providing a cellular basis for their oncogenic potential. Transplanted VHLS-VCs induce excessive blood vessel formation, leading to vascular lesion formation characterized by abundant blood vessels and abnormal cellular growth in immunodeficient mice. To date, VHL-deficient RCC models have identified HIF2A as an essential factor for cellular proliferation, metabolism, and angiogenesis in RCC cells during tumor progression, and contributed to the development of HIF2A-targeted therapies for RCC, including VHLS^10,18,19^. However, RCC cells are distinct from VCs and therefore do not directly address how VHL mutations alter the behavior of human VCs in the context of VHLS. In this regard, our patient-derived vascular model provides an opportunity to investigate the cell-autonomous consequences of VHL inactivation in human VCs and to identify cellular and molecular mechanisms that may not be fully captured by RCC models.

Previous studies suggest that VHLS-associated hemangioblastomas are histologically composed of abundant blood vessels and neoplastic stromal cells arising from mesoderm-derived embryologically arrested hemangioblasts, which are capable of hematopoietic cells and VECs differentiation in multiple organs, and that the blood vessels within VHLS-associated hemangioblastomas might be a result of the mixed population of tumor-derived vasculature and pre-existing blood vessels; however, the origin of hemangioblastoma remains unclear^77,78^. In our model, most of the differentiated VCs and vascular networks are VECs expressing endothelial lineage-specific markers CD31^+^CD144^+^, should be the primary source for blood vessel formation in vitro and in vivo. Indeed, the sorted CD31^+^CD144^+^ VHLS-VECs formed excessive vascular networks compared to healthy-VECs, and the functional blood vessels derived from VHLS-VCs expressed human CD31 in the xenograft mouse model. In addition, VCs and vascular networks contain PDGFRβ^+^or NG2^+^ pericytes and α-SMA^+^-smooth muscle cells, suggesting the presence of other mesodermal cells in the differentiated population. Thus, VHLS-VCs majorly possess cell-autonomous activity to develop the VHL-associated vascular lesions. On the other hand, the vascular lesions in this model also contained stromal-like structures characterized by clear, foamy cytoplasm containing prominent vacuoles, a feature observed in the VHLS-associated hemangioblastomas. However, the immunophenotypic characteristics of hCD31^+^inhibin-alpha^-^ structures did not fully correspond to those of the canonical neoplastic stromal cells in the hemangioblastomas. These structures are therefore more consistent with endothelial or vascular differentiation, and their presence alone does not establish the formation of hemangioblastoma-like stromal cells. Further studies will be required to determine the identity of ectopic blood vessels and abnormal cell clusters in the transplanted vascular lesions using our models. pVHL ubiquitously expresses across various types of organs, tissues, and cells. Nevertheless, the loss of function in pVHL contributes to the formation of organ-specific hemangioblastomas and cancers in VHLS^3^. The definitive answers to this broad question have not been clarified yet. From our study, one possible explanation for this question is that pVHL inactivation causes VEC-specific transcriptional changes related to oncogenic signaling as well as blood vessel formation, including arteriovenous types. Indeed, we did not observe any differences in the proliferation ability and transcriptional changes between the healthy-iPSCs and VHLS-iPSCs. Thus, the oncogenic advantages due to pVHL inactivation might be cell-type specific in the VECs. Given that tumor-specific VECs in human NSCLC acquired oncogenic cellular and molecular types inducing excessive vasculogenesis/angiogenesis^33^, the pVHL-mediated abnormal modifications in VECs may lead to reprogramming of blood vessel identity and organization in VHLS patients.

Although previous reports mainly focused on the central role of pVHL as an E3 ubiquitin ligase, pVHL has multiple functions in diverse cellular processes such as microtubule stabilization and assembly of the extracellular matrix^55,79^. Indeed, we observed that primary cilia formation was significantly inhibited in *VHL*-deficient iPSCs without affecting molecular changes of ciliogenesis-related genes compared to healthy-iPSCs. In addition, HNRNPCKO^+/-^-iPSCs or HIF2AKO^-/-^-iPSCs showed no recovery effects on the percentage of ciliated cells in the VHLS-iPSCs. This data supports the notion that pVHL has a role in ciliary maintenance independent of HIF2A associated with microtubule stability^80^. Moreover, the loss of primary cilia is one of the causes of renal cyst formation. Although little is known regarding the detailed molecular mechanisms, biallelic KO of *VHL* gene impedes primary cilia formation while cooperating with inactivation of phosphatase and tensin homolog (PTEN), which antagonizes PI3K signaling, leading to cystogenesis in the mouse ccRCC model^81^. Correlated with this finding, we detected transcriptional activation of mTORC1 signaling, a downstream of P13K-Akt pathway, in the VHLS-VCs compared to healthy-VCs. These results indicate that loss of function other than as an E3 ubiquitin ligase in pVHL may critically contribute to the formation of cilia, resulting in vascular lesions in our models.

Systemic alteration of RNA processing driven by oncogenic mutations is critical in cancer development; however, the causal relationship is not clear in many cases. We found that HNRNPC, one of the RBPs, was stabilized via the inactivation of pVHL-mediated protein degradation in *VHL*-deficient VCs for blood vessel development and rapid cell growth. The down-regulation of HNRNPC also abrogated pVHL-driven abnormal vasculogenesis independent of HIF2A expression. HNRNPC is crucial in regulating alternative splicing, pre-mRNA maturation, mRNA stabilization, and translocation for controlling its translation^82^. Indeed, the specific transcripts were differentially expressed between healthy-VCs and VHLS-VCs, and the alternative splicing was changed by HNRNPC^+/-^-VCs. HNRNPC up-regulation has been associated with various cancer types, such as cervical cancer^70^ and hepatocellular carcinoma^72^ for their rapid proliferation and migration. Notably, pVHL was found to modify ubiquitination and degradation of HNRNPC independent of HIF2A expression in VCs. Moreover, we observed that HNRNPCKO^+/-^-VCs specifically downregulated gene expression associated with blood vessel development as well as RCC-related signaling in *VHL*-deficient VCs. These findings offer new insights into the mechanism of HNRNPC-mediated RNA metabolism in VHLS pathomechanisms. It will be interesting to characterize translational controls via HNRNPC and its links to specific pathways of RNA metabolism in blood vessel development.

In conclusion, we generated an iPSC-derived vascular platform to elucidate VHLS mechanisms and identified a novel therapeutic target for VHL-associated excessive vasculogenesis. Our studies demonstrated that HNRNPC depletion inhibited the pVHL-driven vasculogenesis by suppressing gene expressions involved in blood vessel development and oncogenic pathways associated with *VHL*-deficient RCC. RBP-regulatory network via pVHL-mediated protein degradation offers a promising approach for cancer therapy.

## Materials and methods

### Human subjects

The generation and use of patient-derived hiPSCs were approved by the Ethics Committee of RIKEN, (Approval numbers: RIKEN-T-2023-004 in RIKEN).

### Animal experiments

All animal experiments were approved by the Animal Experimentation Committee of the RIKEN Tsukuba campus (Approval number: T-2023-007) and were performed according to the committee’s guiding principles and the “Guide for the Care and Use of Laboratory Animals” published by the National Institute of Health.

### Plasmid construction

To generate iPSCs expressing Venus-Akaluc fusion genes, PiggyBac transposase system was applied to integrate the fusion genes into the genome DNA of healthy donors-derived iPSCs and VHLS patients-derived iPSCs. pcDNA3 Venus-Akaluc (#RDB15781, RIKEN BRC DNA BANK) was provided by the RIKEN BRC through the National Bioresource Project of the MEXT/AMED, Japan (RIKEN BRC DNA Bank) and inserted into PiggyBac plasmid vector pPB-PGK-destination (#60436, Addgene) with gateway cloning, and then PiggyBac expression vector pPB-CAG-Venus-Akaluc was constructed (this paper, deposited to Addgene plasmid #209768). For establishing iPSCs expressing *VHL* wild-type gene, pDONR223_VHL_WT (#81874, Addgene) was subcloned into CSII-EF-RfA-IRES-Puro lentivirus vector (#RDB12869, RIKEN BRC DNA Bank) with gateway cloning. CSII-EF-GFP-IRES-Puro (#RDB12868, RIKEN BRC DNA Bank) was used as a control GFP lentivirus vector. The sequence of the expression vectors was confirmed by sanger sequencing at FASMAC. Primers to analyze the sequences are listed in Supplementary Table 1.

### iPSCs culture

One healthy donor-derived iPSC line, WTC11, was provided from Prof. Bruce Conklin in Gladstone Institutes through Coriell Institute as GM25256^83,84^. Other four healthy donor-derived iPSCs, 1383D6, 606A1, 1231A3, and 201B7, and eighteen VHLS patients-derived iPSC lines from three patients, HPS2574, HPS2575, HPS2576, HPS2577, HPS2578, HPS2579, HPS2580, HPS2581, HPS2582, HPS2583, HPS2584, HPS2585, HPS2592, HPS2593, HPS2594, HPS2595, HPS2596, and HPS2597, were provided by the RIKEN BRC through the National BioResource Project of the MEXT/AMED, Japan. All of these iPSCs were cultured under feeder-free conditions as previously described^85,86^. The cells were seeded on 0.25 μg / cm^2^ iMatrix-511 silk (#892011, Matrixome) in StemFit AK02N complete medium (#RCAK02N, Ajinomoto) supplemented with 10 μM Y-27632 (#034-24024, FUJIFILM Wako Pure Chemical) and maintained in the medium without Y-27632 from the next day. The medium was changed every other day.

### Mycoplasma tests

VHLS patients-derived iPSC lines (HPS2574, HPS2583, HPS2592) were confirmed to be mycoplasma-negative by indirect DNA fluorescent staining and a nested-PCR. Medium supernatant of iPSC culture was tested by staining with Hoechst33258 (Sigma) after 5–6 days of co-culture with VERO cells (#RCB0001, RIKEN BRC Cell Bank) as mycoplasma negative indicator cells. DNA was extracted using DNeasy Blood and Tissue Kit (#69506, QIAGEN) and subjected to a nested-PCR. AmpliTaq Gold 360 DNA Polymerase (Thermo Fisher Scientific) was used for each step of the PCR. The same thermocycling conditions were used for both steps: initial denaturation of 10 minutes at 95 °C, 30 cycles of thermocycling; 30 seconds at 95 °C, 2 minutes at 55 °C, 2 minutes at 72 °C; and final extension of 5 minutes at 72 °C, and hold at 4 °C. The primers used in the PCR are listed in Supplementary Table 1.

### STR (short tandem repeat) analysis

The identity of these iPSC lines was verified with short tandem repeat (STR) analysis. Genomic DNA was extracted from VHLS patients-derived iPSCs (HPS2574, HPS2583, and HPS2592) with DNeasy Blood & Tissue Kit, and an STR analysis was performed with PowerPlex16 System (#DC6531, Promega).

### Copy number variation (CNV) / SNP (short nucleotide polymorphism) microarray assay (virtual karyotyping)

Genomic DNA was extracted from VHLS-iPSCs (HPS2574, HPS2583, and HPS2592) using a DNeasy Blood & Tissue Kit for virtual karyotyping. According to the manufacturer’s instructions, CNV/SNP microarray assay was performed with GeneChip Scanner System 3000 (Thermo Fisher Scientific) and Karyostat Assay arrays (#905403, Thermo Fisher Scientific). The results were analyzed using Chromosome Analysis Suite (Thermo Fisher Scientific) and GeneChip Command Console Software (Thermo Fisher Scientific).

### Generation of Healthy-iPSCs and VHLS-iPSCs expressing Venus-Akaluc

Healthy-iPSCs and VHLS-iPSCs were dissociated with Accutase (#12679-54, Nacalai tesque) and 3×10^5^ cells were resuspended in the StemFit AK02N complete medium supplemented with 10 μM Y-27632 and seeded on 0.25 μg/cm^2^ iMatrix-511 silk in 6 well plate (#3810-006N, IWAKI). Next day, the cells were transfected with 50 µL of Opti-MEM I Reduced Serum Medium (#31985070, Gibco) containing 10 µL of Lipofectamine Stem Reagent (#STEM0001, Invitrogen), 1.9 µg of pPB-CAG-Venus-Akaluc (this paper, deposited to Addgene plasmid #209768), and 0.6 µg of pHL-EF1a-hcPBase-iP-A (#RDB13322, RIKEN BRC DNA Bank). After 20 hours, the medium was changed to StemFit AK02N complete medium. On day 3, the transfected cells were treated with 50 µg / mL of hygromycin (#09287-84, Nacalai tesque) until the cells reached semi-confluent. Then, Venus^+^ cells were sorted with SH800 cell sorter (SONY) and maintained in StemFit AK02N complete medium.

### Establishment of gene knockout (KO) iPSC lines with CRISPR-Cas9 system

To construct gene knockout lines, the synthetic guide RNA-purified protein (RNP) complex with Alt-R S.p. HiFi Cas9 Nuclease V3 (#1081058, Integrated DNA Technologies) targeting an exon of human genes was transduced into healthy donors and VHLS patients-derived iPSCs. All crRNAs specific to human genes such as *VHL*, *HIF*2A, and *HNRNPC* were designed based on CRISPR direct software (https://crispr.dbcls.jp/). The sequence of designed crRNAs is listed in Supplementary Table S1. In brief, gRNAs were synthesized from 10 µM crRNA hybridized with 10 µM tracrRNA (#1072532, IDT) at 95 °C for 5 minutes. Subsequently, the RNP complex was formed from 5 µM gRNAs and 1.0 µg of Alt-R S.p. HiFi Cas9 Nuclease V3 incubated at room temperature for 15 minutes. Then, the RNP complex was diluted in 49 µL of Opti-MEM I Reduced Serum Medium and incubated at 4℃. The healthy donors and VHLS patients-derived iPSCs were dissociated into single cells with Accutase. And 2.5×10^5^ cells were resuspended in 100 µL of RNP complex and electroporated by using NEPA21 electroporator (NEPAGENE). After electroporation, the iPSCs were seeded on 0.25 µg/cm^2^ iMatrix-511 silk and cultured in StemFit AK02N medium supplemented with 10 µM Y-27632. The transfected single colonies were picked up and further expanded for genotyping using sanger sequencing at FASMAC. Primers to analyze the sequences are listed in Supplementary Table 1.

### Generation of VHLS-iPSCs expressing lentivirus vectors encoding *GFP* or *VHL* genes

293T human embryonic kidney cell line (RCB2202 in RIKEN Cell bank) was cultured in DMEM (#08458-16, Nacalai tesque) supplemented with 10% FBS (#FB-1365/500, Biosera) and 1% Penicillin-Streptomycin Mixed Solution (#26252-94, Nacalai tesque), and then transfected with packaging plasmid (pCAG-HIVgp) (#RDB04394, RIKEN BRC DNA Bank), envelope plasmid (pCMV-VSV-G-RSV-Rev) (#RDB04393, RIKEN BRC DNA Bank), and CSII-EF-GFP-IRES-Puro lentivirus vector or the CSII-EF-VHL-IRES-Puro lentivirus vector using FuGENE 6 Transfection Reagent (#E2691, Promega). Culture supernatants containing lentiviruses were harvested 48–72 hours after transfection and filtered with 0.45 μm Syringe Filter (AS ONE). After mixing the supernatants with Lenti-X Concentrator (#631231, Takara), the lentivirus mixture was incubated overnight at 4°C and used for infection. VHLS-iPSCs were dissociated with accutase and 1×10^5^ cells were resuspended in the StemFit AK02N complete medium supplemented with 10 μM Y-27632 and seeded on 0.25 μg/cm^2^ iMatrix-511 silk in 6 well plate. Next day, the medium was switched to the StemFit AK02N complete medium supplemented with 10 μM Y-27632 and 8 µg/ml polybrene solution (#12996-81, Nacalai tesque). VHLS-iPSCs were infected with the lentivirus mixture using a spin infection procedure (32℃, 1,200G, 60 min). Three days after infection, the cells were treated with 1.0 μg/mL puromycin (#19752-64, Nacalai tesque) to obtain the drug-resistant cells.

### Embryoid body (EB) formation in vitro

Healthy-iPSCs and VHLS-iPSCs were dissociated with 0.5 mM EDTA (#13567-84, Nacalai tesque) in PBS (#14249-95, Nacalai tesque) and 1.0 x 10^4^ cells were suspended in Stem Fit AK02N medium supplemented with 10 μM Y-27632. EBs were aggregated in a well of an EZ-BindShut 96-well-V plate (#4420-800LP, IWAKI). The next day, EBs were transferred in DMEM high Glucose (#08458-16, Nacalai tesque) supplemented with 10% fetal bovine serum (EB medium) and cultured for 8 days. After that, EBs were seeded on 0.1 w/v% gelatin solution (#190-15805, FUJIFILM Wako Pure Chemical Corporation)-coated plate and further cultured in EB medium for another 8 days.

### Teratoma formation

Healthy-iPSCs and VHLS-iPSCs with/without Venus-Akaluc genes were dissociated with 0.5 mM EDTA in PBS and resuspended in 50% Matrigel (#354230, Corning): StemFit AK02N medium with 10 μM Y-27632 = 1:1. 1.0 x 10^6^ cells were transduced into the testis and legs of NOD.Cg-Prkdcscid Il2rgtm1Wjl/SzJ (NSG) mice (The Jackson Laboratory)^87,88^. After around two months, the mice were sacrificed and the teratoma tissues were isolated. The teratomas were fixed with 4% paraformaldehyde (#09154-85, Nacalai tesque) in PBS, then hematoxylin-eosin (HE) staining was performed by Genostaff, Inc.

### Vascular differentiation from iPSCs

iPSC lines were dissociated with 0.5 mM EDTA in PBS and subsequently incubated at 37 °C with 5% CO_2_ for 5min. After scraping and pipetting, 5.0×10^4^ cells were resuspended in the StemFit AK02N complete medium supplemented with 10 μM Y-27632 and seeded on 0.25 μg/cm^2^ iMatrix-511 silk in 60 mm dish (#3010-060, IWAKI). From day 1 to day 4, the cells were cultured in StemFit AK02N medium without supplement C together with 8 μM CHIR99021 (#034-23103, FUJIFILM Wako Pure Chemical), 50 ng/mL BMP4 (#020-18851, FUJIFILM Wako Pure Chemical), and 1% DMSO (#D8418, Sigma) for mesodermal induction. On day 4, the culture medium was changed to Stempro-34 SFM medium (#10639011, ThermoFisher) supplemented with 50 ng/mL VEGF (#226-01786, FUJIFILM Wako Pure Chemical), 50 ng/mL FGFb (#060-04543, FUJIFILM Wako Pure Chemical), and 10 μM SB431542 (#037-24293, FUJIFILM Wako Pure Chemical) for vascular cell induction. Then, the induction medium was changed every other day. On day 11, the vascular cells (VCs) were passed and expanded for further experiments. The VCs were incubated with 0.5 mM EDTA in PBS at 37 °C with 5% CO_2_ for 5min. After only pipetting, 1.0-2.0×10^5^ cells were resuspended in the induction medium supplemented with 10 μM Y-27632 and seeded on 0.25 μg/cm^2^ iMatrix-411 silk (#892-042, Matrixome) in a 60 mm dish. And then, the induction medium without Y-27632 was changed every other day.

### Construction of three-dimensional vascular networks from vascular cells

Three-dimensional vascular networks were generated from VCs or sorted VECs (CD31⁺CD144⁺), referring to previous reports with modification^31,43^. VCs after passage or the sorted VECs were cultured in the induction medium by day 14, then maintained in Endothelial Growth Medium 2 (EGM2; #C-22111, TAKARA) for 3 days. On day 17, the VCs or sorted VECs were dissociated with 0.5 mM EDTA in PBS. After only pipetting, 2.0 x10^3^ (for VHLS patients vs. healthy donors) or 4.0×10^3^ cells (for VHLS-WT vs. HIF2AKO lines or HNRNPCKO lines) were resuspended in the 100 µL maintenance medium and seeded onto a well of V-bottom 96 well plate (#4420-800LP, IWAKI). To form an aggregation of the VCs or VECs, the cells are centrifuged at 200 G for 3 min and then incubated for 4 days. Afterward, an aggregate was transferred into a 30 µL Type I-A collagen drop (#631-00651, Nitta gelatin) on the bottom of a 35 mm culture dish (#3000-035, IWAKI) and cultured in the maintenance medium for 6 days. The maintenance medium was changed every other day.

### Knockdown of VHL gene via dicer-substrate small interfering RNA (DsiRNA)

DsiRNAs targeting an exon of human VHL gene was transduced into VHLS patient-derived vascular cells (HPS2574). DsiRNA#1 (hs.Ri.VHL.13.1), DsiRNA#2 (hs.Ri.VHL.13.2), and negative control (DS NC1) were designed by IDT. In brief, vascular cells after passage were cultured in the induction medium by day 14, then maintained in EGM2 for 2 days. On day 16, the cells were transfected with 400 µL of Opti-MEM I Reduced Serum Medium (#31985070, Gibco) containing 12 µL of Lipofectamine RNAiMAX Regenet (#13778030, ThermoFisher Scientific) and 4 µL of 10 µM DsiRNAs. After 16 hours, the medium was changed to EGM2 and the cells were used for further assays.

### Evaluation of inhibitory effects of chemical inhibitors on vasculogenesis in VCs or vascular networks

To evaluate inhibitory effects of pharmacological inhibitors, SU5402 (#197-16731, FUJIFILM Wako Pure Chemical) and PT2977 (#S8886, Selleck Biotechnology), on vasculogenesis, three dimensional vascular networks from Healthy-VCs and VHLS-VCs were treated with DMSO, SU5402 and PT2977 at 100 nM, 1 µM, and 10 µM from day 4 to day 10 after transferring a VCs-aggregate into a collagen drop. Then, sprouting vessel lengths were quantified by ImageJ software (https://imagej.net/ij/index.html). On the other hand, Healthy-VCs and VHLS-VCs were treated with these chemicals at 1 nM, 10 nM, 100 nM, 1 µM, and 10 µM from day 1 to day 7 in monolayer cultures. At day7, cell numbers in the VCs were counted on each concentration. During the chemical treatment, vascular networks or VCs were cultured in EGM2 at 37 °C with 5% CO_2_. The medium with these chemicals was changed every other day.

### A xenograft mouse model transplanted with vascular cells

Healthy-VCs, VHLS-VCs, and gene-edited VCs expressing Venus-Akaluc genes were transplanted under the kidney capsule of 8-16-week-old immunodeficient NSG mice (NOD.Cg-Prkdc^scid^Il2rg^tm1Wjl^/SzJ). Before transplantation, VCs maintained in EGM2 were dissociated with 0.5 mM EDTA in PBS, and 1.0×10^6^ cells were resuspended in 50% Geltrex (#A1413302, ThermoFisher): EGM2 with 10 μM Y-27632 = 1:1. The cell suspension was injected into a mouse kidney. Mice were imaged using IVIS imaging system (LuminaXRMS seriesⅢ ver4.7) to monitor the transplanted cells at several weeks after the transplantation. Approximately 10 min before the imaging, the mice were intraperitoneally administered with 100 µL of 30 mM AkaLumine-HCl (#016-26704, FUJIFILM Wako Pure Chemical), and luminescent images were acquired under the following conditions: open for total bioluminescence, exposure time = 5 sec, binning = medium, and f/stop = 1. The total flux was visualized with R software program (version 4.2.2). Kidney tissues in mice were isolated at 12 weeks after the transplantation and fixed with 4% paraformaldehyde in PBS. Then, the tissue sectioning, hematoxylin-eosin (HE) staining, and immunohistochemistry (IHC) staining were performed by Genostaff, Inc.

### Measurement of Akaluc-derived luminescent intensity in iPSCs

Healthy-iPSCs, VHLS-iPSCs, and gene-edited iPSCs expressing Venus-Akaluc genes were seeded (3.0×10^4^ cells/well) on 96 well flat-bottom plate (IWAKI) coated with 0.25 μg/cm^2^ iMatrix-511 silk in StemFit AK02N complete medium supplemented with 10 μM Y-27632. Next day, the medium without Y-27632 was changed. Approximately 10 min before the imaging, the iPSCs were treated with AkaLumine-HCl from 10 µM to 1 mM, then the luminescence intensity was measured by spark multimode microplate reader (TECAN) at exposure time = 10 sec.

### Growth curve analysis

VCs were dissociated with 0.5 mM EDTA in PBS and incubated at 37 ◦C with 5% CO_2_ for 5min. After pipetting, 1.0×10^4^ cells/mL were seeded on 0.25 μg/cm^2^ iMatrix-411 silk in 6 well plate and cultured in the Stempro-34 SFM medium supplemented with 50 ng/mL VEGF, 50 ng/mL FGFb, 10 μM SB431542, and 10 μM Y-27632. From next day, the cells were maintained in EGM2 and counted every day. The medium was changed every other day.

### Migration assay

VCs were dissociated with 0.5 mM EDTA in PBS and subsequently 1.0×10^5^ cells/well were seeded on 0.25 μg/cm^2^ iMatrix-411 silk in µ-Plate 24 Well Black ID 14 mm (#ib80242, Nippon Genetics), and cultured in the Stempro-34 SFM medium supplemented with 50 ng/mL VEGF, 50 ng/mL FGFb, 10 μM SB431542, and 10 μM Y-27632. The next day, the culture medium was switched to EGM2, and then time-lapse images were taken for 16 hours under an all-in-one fluorescent microscope (BZ-X800; KEYENCE).

### Flow cytometry

iPSCs or VCs were dissociated with 0.5 mM EDTA solution in PBS, the single cells (1.0 – 2.0 × 10^5^ cells) were centrifuged at 200 × g for 3 minutes, and resuspended in 100 μL PBS containing 0.1% bovine serum albumin (#015-23871, BSA; FUJIFILM Wako Pure Chemical). On the other hand, the three vascular networks in hydrogels were added with cell recovery solution (#354253, corning) and incubated at 4℃ for 5 min. After PBS washing, the vascular networks were transferred into 1mg/mL collagenase/Dispase Blend solution (#SCR139, Sigma-Aldrich) and incubated at 37℃ for 30 min. The reaction was stopped by the addition of EGM2, then the cells were centrifuged at 300g for 5 min. After PBS washing, the cells were resuspended in 100 μL PBS containing 0.1% BSA. And then, the cells were stained with the following antibodies for 1.5 hour at 4 °C: Alexa Fluor 488 Mouse anti-TRA-1-60 (#330614, BioLegend), DyLight 550 Mouse anti-SSEA-4 (#09-0097, Stemgent), APC anti-human CD31 Antibody (#303116, BioLegend), FITC Mouse Anti-Human CD144 (#560411, BD bioscience), After washing with PBS, the cells were resuspended in 300 µL PBS containing 0.1% BSA and filtrated with 35-mm cell strainer (Thermo Fisher Scientific) for cell analysis or cell sorting with SH800 cell Sorter (SONY).

### Live/Dead assay for vascular networks

Cell viability for iPSCs and vascular networks were evaluated with the Cyto3D Live-Dead Assay Kit (#BM01, The Well BIOSCIENCE) following the manufacturer’s instructions. Culture medium in iPSCs and vascular networks were replaced with the fresh medium and added with 20 µL/mL Cyto3D reagent at 37℃ for 15-30min. Negative controls were fixed 4% paraformaldehyde for 10-30 min at room temperature, and changed the 4% PFA with PBS including 20 µL/mL Cyto3D reagent after PBS washing. Samples were observed under an all-in-one fluorescent microscope (BZ-X800; KEYENCE). For flow cytometric analysis, the vascular networks in hydrogels were added with cell recovery solution (#354253, corning) and incubated at 4℃ for 5 min. After PBS washing, the vascular networks were transferred into 1mg/mL collagenase/Dispase Blend solution (#SCR139, Sigma-Aldrich) and incubated at 37℃ for 30 min. The reaction was stopped by the addition of EGM2, then the cells were centrifuged at 300g for 5 min. After PBS washing, the cells were resuspended in 300 μL PBS containing 0.1% BSA and filtrated with 35-mm cell strainer (Thermo Fisher Scientific) for SH800 cell Sorter (SONY).

### Immunocytochemistry

Immunocytochemistry was performed for self-renewal markers, three germ layer markers, and a primary cilia marker in iPSCs or blood vessel markers in VCs and vascular networks. In dextran assay, vascular networks were treated with RFP-Dextran (#D1863, ThermoFisher) at 25µg/mL for 15 min before immunocytochemistry. For the iPSCs and VCs, the cells were fixed with 4% paraformaldehyde for 10 min at room temperature. After PBS washing, the cells were permeabilized in PBS containing 0.1% Triton X-100 (#35501-15, Nacalai tesque) for another 10 min at room temperature, then washed with PBS. Primary antibodies were incubated in PBS containing 0.1% BSA overnight at 4 ◦C. Secondary antibodies were incubated for 1 h at room temperature in PBS containing 0.1% BSA. For the vascular networks, the samples were fixed with 4% paraformaldehyde for 30 minutes at room temperature. After PBS washing, these vascular networks were permeabilized in PBS containing 0.1% Saponin (#30502-55, Nacalai tesque) for 2 hours at room temperature and were rinsed with PBS. These vascular networks were incubated in PBS containing 0.1% BSA and primary antibodies overnight at 4 ◦C. Next, secondary antibodies were added to these vascular networks and incubated for 2 h at room temperature in PBS containing 0.1% BSA. Fluoro-KEEPER Antifade Reagent DAPI (#12745-74, Nacalai Tesque) was used for nuclear counterstaining. Samples were observed under an all-in-one fluorescent microscope (BZ-X800; KEYENCE). Information on the primary and secondary antibodies are listed in Supplementary Table 2 and 3.

### Western blotting

Protein expressions were analyzed with an automatic capillary western blot (Simple Western, Wes; Bio-Techne) according to the manufacturer’s instructions. The cell lysate was prepared from 1 × 10^5^ - 1 × 10^6^ cells with sodium dodecyl sulfate-polyacrylamide gel electrophoresis (SDS-PAGE) sample buffer solution without 2-ME (2x) (#30567-12, Nacalai Tesque) containing 100 mM Dithiothreitol (DTT) (#044-33873, FUJIFILM Wako Pure Chemical), then incubated for 5 min at 95 °C. When detecting hypoxia-inducible factors (HIFs), the cells were placed in an incubator at 37°C and 1% O_2_ for 6 hours before the protein collection and lysed with SDS-PAGE sample buffer solution containing 100 mM DTT on ice. Primary and secondary antibodies are listed in Supplementary Table 2 and 3. The protein expressions were normalized with GAPDH and quantified by compass for simple Western software program (ProteinSimple).

In the *in vivo* binding assay, VCs were lysed in RIPA buffer (#08714-04, Nacalai tesque) containing protease inhibitor cocktail (#03969-21, Nacalai tesque), and the cell lysates were incubated with Rabbit mAb HNRNP C1/C2 antibody (#ab133607, Abcam), Rabbit mAb HIF2A (#71565, Cell signaling Technology) or Rabbit (DA1E) mAb IgG XP Isotype Control (#3900, Cell signaling Technology) at 4℃ overnight. After washing with RIPA buffer, the cell lysates were incubated with Protein A Magnetic Beads (#73778, Cell signaling Technology) for 2 hours at 4 °C. The Magnetic Beads were then washed four times with RIPA buffer and bound proteins were resolved by SDS-PAGE sample buffer solution without 2-ME (2x) containing 100 mM DTT. After boiling at 95 °C for 5 min and cooling down to room temperature, protein expression was analyzed with the Wes assay.

For in vivo ubiquitination assay, VCs were incubated in the presence or absence of 5 µM MG132 (#139-18451, FUJIFILM Wako Pure Chemical) for 1.5 or 3 hours before the protein collection. Cells were lysed in RIPA buffer containing protease inhibitor cocktail, and the cell lysates were immunoprecipitated with HNRNP C1/C2 antibody or Rabbit (DA1E) mAb IgG XP Isotype Control. After that, the cell lysates were diluted with SDS-PAGE sample buffer solution without 2-ME (2x) containing 100 mM DTT and were incubated for 5 min at 95 °C. Protein expressions were analyzed with an automatic capillary western blot.

### Proteomics analysis

VCs derived from the healthy donors and VHLS patients were lysed with a lysis buffer including 1M Tris-HCl pH 7.6 (#35436-01, Nacalai tesque), 0.1 M DTT, and 2% SDS (#31607-65, Nacalai tesque). After ultrasonic disruption with Covaris S2 (Duty Cycle 20%, Intensity 5, Cycles/Burst 200) (M&S Instruments Inc.), the cell solutions were centrifuged at 16,000 × g for 10 minutes, then the supernatants were collected. The proteins were precipitated with methanol (#133-15355, FUJIFILM Wako Pure Chemical) and chloroform (#034-02603, FUJIFILM Wako Pure Chemical), followed by resolving with a solubilize buffer including 12 mM SLS (#20117-12, Nacalai tesque), 12 mM SDC (#10712-12, Nacalai tesque), and 50 mM TEAB (#T7408-100ML, Sigma). The proteins were added with 1M DTT and incubated at 37℃ for 30 min. Moreover, 500 mM IAA (#19302-54, Nacalai tesque) and 1 µg/µL Lys-C (#125-05061, FUJIFILM Wako Pure Chemical) were mixed with the proteins at 37℃ for 4 hours, and the proteins were digested with 50 mM TEAB buffer including 1 µg/µL trypsin (#V5111, Promega) at 37℃ for 16 hours. The digested peptides were acidified, desalted, and analyzed for liquid chromatography with tandem mass spectrometry (LC–MS/MS) at Proteomics Core Facility in RIKEN Center for Biosystems Dynamics Research. A 135 min gradient elution separated the peptides, and the source data was available through an orbitrap eclipse (Thermo Fisher Scientific) coupled with a nanoflow UHPLC system (Vanquish; Thermo Fisher Scientific). The peptides were analyzed with a CLC genomics workbench v20.0.2 (QIAGEN), GSEA software v4.3.2, and Metascape v3.5 (https://metascape.org/). In volcano plots, the upregulated genes with log_2_ fold change ≧ 1 (P-value < 0.05) are indicated in red, whereas the downregulated genes with log_2_ fold change ≦ −1 (P-value < 0.05) are colored in blue. In bar graph for GSEA results, all gene categories with FDR q-value < 0.05 are shown. The analyzed data were visualized using with R software program (version 4.2.2).

### RT-qPCR

Total RNA was extracted with a FastGene RNA premium kit (#FG-81050, NIPPON Genetics), and cDNA was synthesized with ReverTra Ace qPCR RT kit (#FSQ-101, TOYOBO) according to the manufacturer’s instructions. The cDNA was used for Real-Time qPCR reactions in QuantStudio 3 system (Thermo Fisher Scientific) with THUNDERBIRD Probe qPCR Mix (#QPS-101, TOYOBO) and TaqMan probes (Thermo Fisher Scientific). The probe sequences are listed in Supplementary Table 1. Gene expression was described as the fold change relative to the control sample value (ΔΔCt method) after normalization to the corresponding GAPDH values.

### RNA-seq

Total RNA derived from iPSC and VCs were extracted with the FastGene RNA premium kit (#FG-81250, Nippon Genetics) and strand-specific library preparation was performed according to the manufacturer’s instructions. 1.5 µg of the purified RNA was sequenced in a NovaSeq/HiSeq (Illumina). Sequencing was performed in a 150 bp × 2 paired-end configuration with a data output of 6 Gb per sample (∼ 20 million PE reads). RNA-seq sequencing reads were aligned to hg19 Homo sapiens reference genome. A CLC genomics work bench v20.0.2 (QIAGEN) was used to calculate differential expression genes. Gene set enrichment analysis (GSEA) was performed with GSEA software v4.3.2 and gene ontology was investigated with Metascape v3.5 (https://metascape.org/). In volcano plots, the upregulated genes or transcripts with log_2_ fold change ≧ 1 (P-value < 0.05) are indicated in red, whereas the downregulated genes or transcripts with log_2_ fold change ≦ −1 (P-value < 0.05) are colored in blue. In bar graph for GSEA results, all gene categories with FDR q-value < 0.05 or 0.10 are shown. The analyzed data were used to visualize with R software program (version 4.2.2).

### Single cell RNA-seq

Single-cell RNA sequencing in the VCs from healthy donors (1383D6 and 1231A3) and VHLS patients (HPS2574 and HPS2583) was conducted using the GEM-X Universal 5’ Gene Expression v3 4-plex kit (PN-1000780, PN-1000748, 10x Genomics) following the manufacturer’s instructions. Libraries were sequenced on an Illumina NovaSeq 6000 platform, generating 2×150 bp paired-end reads. Single-cell RNA-seq reads were mapped to the human reference genome (GRCh38/hg38) and matrix files were generated using Cell Ranger (10x Genomics, v10.0.0). Downstream analyses were performed using Seurat (v5.4.0), dplyr (v1.2.0), and patchwork (v1.3.2)^89^. Cells were filtered with >7,500 detected genes or <200 detected genes, and with >20% mitochondrial transcript counts. The top 2000 highly variable genes were used for Principal component analysis (PCA), followed clustering and visualization with uniform manifold approximation and projection (UMAP).

### Bioinformatic analysis of gene expressions or survival data in cell lines and RCC patients

Human gene expression (RSEM norm_count) data, proteomics data, and overall survival data were obtained from data sets in TCGA Kidney Clear Cell Carcinoma (n = 531) and GTEx (n = 28) cohorts from UCSC Toil RNA-seq Recompute on UCSC Xena browser (https://xena.ucsc.edu/). A Kaplan-Meier survival curve was drawn with the high expression group of *HNRNPC* (RSEM normalized count ≧ 13.98), *HNRNPD* (RSEM normalized count ≧ 12.69), *DCTPP1* (RSEM normalized count ≧ 9.949), *MYC* (RSEM normalized count ≧ 11.63), *EPAS1* (RSEM normalized count ≧ 14.87) and the low expression group of *HNRNPC* (RSEM norm_count < 13.98), *HNRNPD* (RSEM norm_count < 12.69), *DCTPP1* gene (RSEM norm_count < 9.949), *MYC* (RSEM norm_count < 11.63), *EPAS1* (RSEM norm_count < 14.87) in TCGA Kidney Clear Cell Carcinoma patients. GSEA data and gene expressions of RNA-seq or proteomics in 786-O RCC lines, HUVEC, HUAEC, iPSCs-derived vein, and iPSCs-derived artery were acquired from a previous report^15,57,58^. The analyzed data were visualized using R software program (version 4.2.2).

### Statistical analysis

Statistical analysis was performed using a two-tailed Student’s t-test, a two-tailed log-rank test, one-way ANOVA Dunnett’s test, or Tukey’s test for multiple comparisons. Statistical P-values were determined with R software program (version 4.2.2) and Microsoft Excel software program for Windows (Microsoft). P values < 0.05 were considered statistically significant. *, **, or *** in the graphs indicate P<0.05, P<0.01, or P<0.001, respectively. The experiments were not randomized and the investigators were not blinded to allocation during experiments and outcome assessment.

## Author Contributions

Hidenori Ito and Yohei Hayashi provided the conception and design of the study. Hidenori Ito, Iori Sato, Tomoya Shimizu, Michiya Noguchi, Yukio Nakamura, Naoki Kubo, Daigo Takagi, Masaru Tamura and Yohei Hayashi acquired data analysis.

Hidenori Ito, Mami Matsuo-Takasaki, Naoki Kubo, Daigo Takagi, and Yohei Hayashi interpreted the results.

Hidenori Ito and Yohei Hayashi wrote the draft of the manuscript.

Hidenori Ito, Mami Matsuo-Takasaki, Megumu K Saito, Shunsaku Takayanagi, Eijiro Nakamura and Yohei Hayashi provided the study materials, laboratory samples, and instrumentation.

Hidenori Ito and Yohei Hayashi acquired financial supports.

Hidenori Ito, Mami Matsuo-Takasaki, Tomoya Shimizu, Naoki Kubo, Daigo Takagi, Masaru Tamura, Megumu K Saito, Shunsaku Takayanagi, Nobuhito Saito, Eijiro Nakamura and Yohei Hayashi analyzed and interpreted the data and revised the manuscript critically for important intellectual content.

Hidenori Ito and Yohei Hayashi managed and coordinated the responsibility for the research activity planning and execution.

## Acknowledgements

We gratefully acknowledge Dr. Toshihiko Shiroishi for his valuable supervision throughout this study, Dr. Kaoru Saijo, Dr. Yutaka Arai, Dr. Tamami Wakabayashi, Ms. Yasuko Hemmi, and Ms. Eiko Yokoyama for their technical assistance, and Ms. Kumiko Omori for her administrative support. This work was supported by the RIKEN internal grants including Special Postdoctoral Researcher Program to H.I, Grants-in-Aid for Scientific Research from the Ministry of Education, Science, and Culture of Japan (23K14457 and 21K20668) to H.I, a grant from AMED (23bm1423010h0001) to Y.H., and grants from AMED (JP23bm1423018 and JP23bm1323001) to M.K.S.

## Data availability

Details of the experiments are available within the article and the Supplementary Information. RNA-seq data or Single cell RNA-seq are publicly available in the Gene Expression Omnibus under accession number JGAS000856 or JGAS000965. Proteomics data can be viewed at the ProteomeXchange under accession number PXD068400. Source data is provided with this paper.

---

**Extended Data Figure 1.**
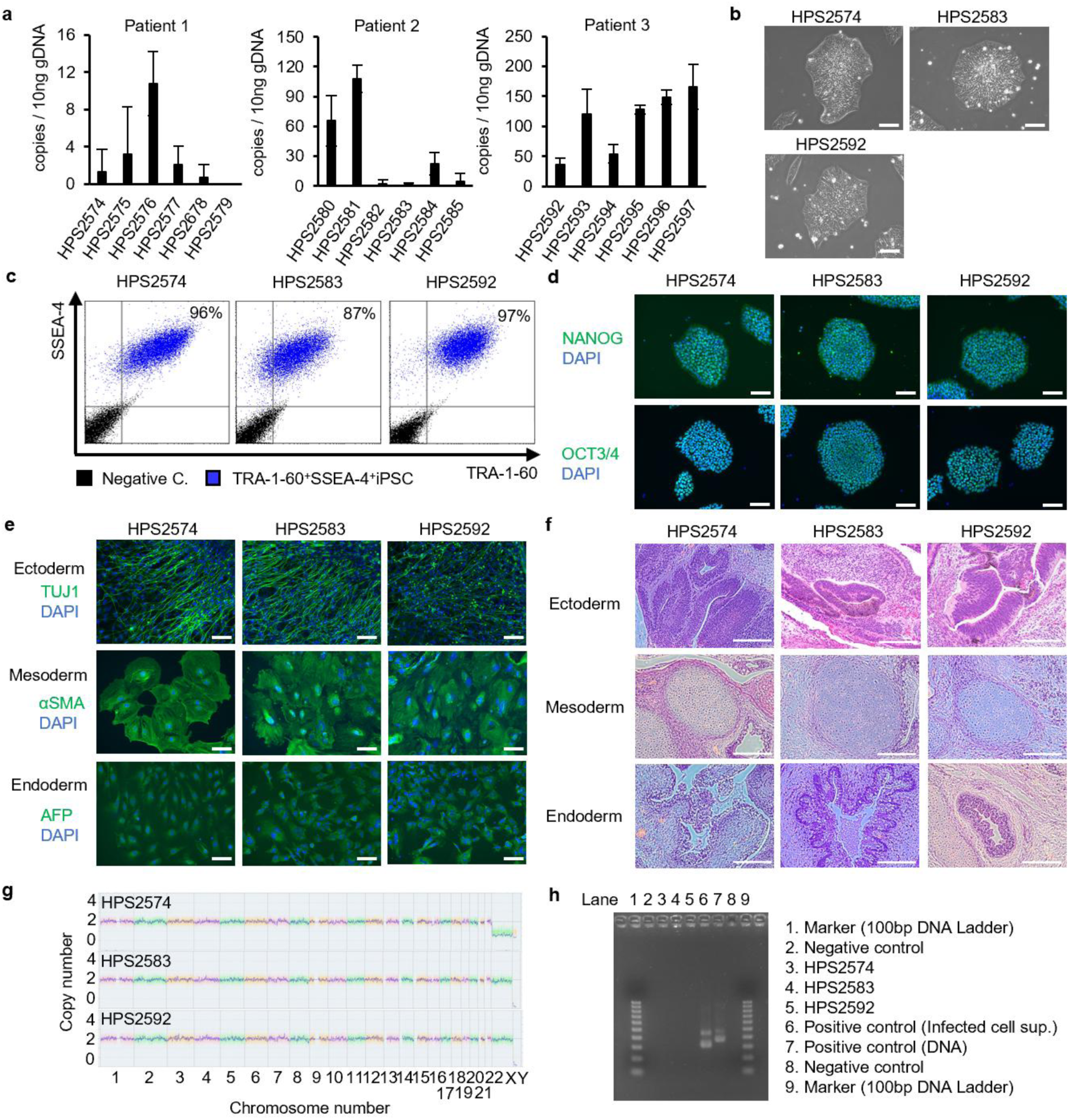
Characterization of VHLS-iPSCs from three VHLS patients. a, Copy numbers of transgene *Ebnal* in 18 clones of VHLS-iPSCs were calculated by genomic quantitative PCR. Data are shown as mean + SEM. b, Representative phase images of three VHLS-iPSCs, HPS2574, HPS2583, and HPS2592 Scale bar=100 μm. c, Flow cytometric analysis for self-renewal markers, SSEA-4. and TRA-1-60, in VHLS-iPSCs. d, Expression of self-renewal markers of iPSCs, NANOG (green) and OCT3/4 (green) and in VHLS-iPSCs. Nucleus was stained with DAPI (blue). Scale bar, 100 μm. e, f, Pluripotency in VHLS-iPSCs was evaluated with in vitro embryoid body (EB) formations and in vivo teratoma formations. TUJ1 (ectoderm marker), alpha-SMA (mesoderm marker), and AFP (endoderm marker) expressions were used for differentiation markers of three germ layers in EB. Scale bar, loop mine, Scale bar, 200 μm in f. g, Virtual karyotyping analysis (CNV/CGH microarray) of three VHLS patient lines without noticeable aberrant copy numbers, h, Mycoplasma tests to check mycoplasma-negative in three VHLS-iPSCs by indirect DNA fluorescent staining and a nested-PCR. Genomic DNA extracts from Mycoplasma fermentans (DNA) and culture medium of VERO cells contaminated with Mycoplasma hyorhin is (Infected cell supernatant) were used as positive controls. One hundred bp DNA ladder was used as DNA size markers.

**Extended Data Figure 2.**
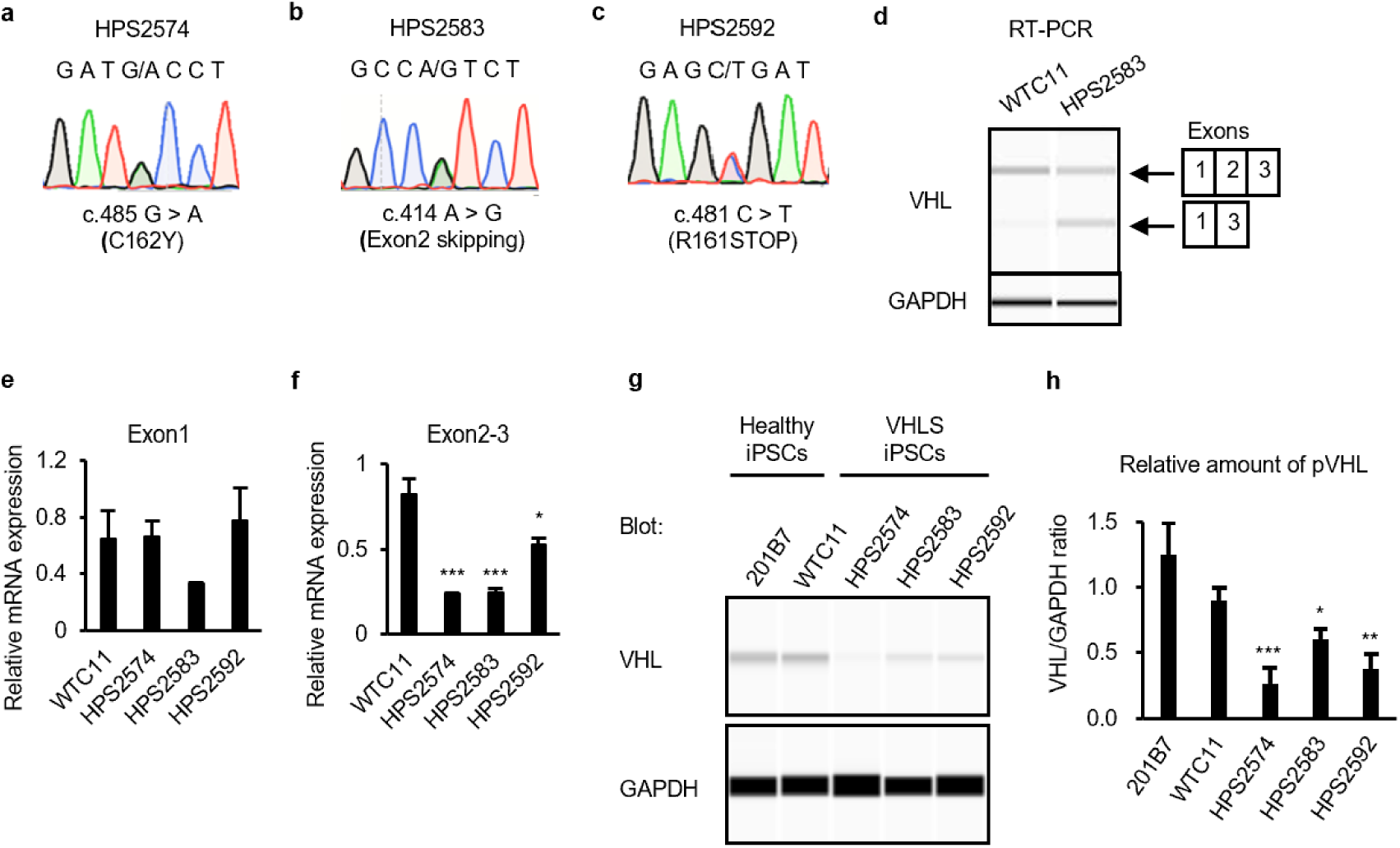
Identification of *VHL* gene mutations in three VHLS-iPSCs. a-c, Sequencing analysis of VHL-mutations in HPS2574 (c.485 G > A), HPS2583 (c.414 A > G), and HPS2592 (c.481 C > T). d, PGR analysis was performed to identify Exon2 skipping with specific primers for *VHL* and *GAPDH* genes in HPS2583. WTC11 was used as a control Healthy-iPSCs. e, f, Gene expression of Exonl (e) and Exon2-3 (f) in *VHL-gene* was evaluated in the IPSCs from Healthy-iPSCs (WTC11)and VHLS patients (HPS2574, HPS2583, HPS2592) by real-time qPCR. g, Protein expression profiles of VHL and GAPDH in Healthy-iPSCs (WTC11 and 201B7) and three VHLS-iPSCs (HPS2574, HPS2583, and HPS2592) by an automatic capillary western blot (WES), h, Relative amounts of VHL proteins normalized by GAPDH in f were quantified using WES. Data are shown as the mean ± SEM (n = 3). P-values were determined by one-way ANOVA Tukey’s test (*P < 0.05, **P < 0.01, ***P < 0.001).

**Extended Data Figure 3.**
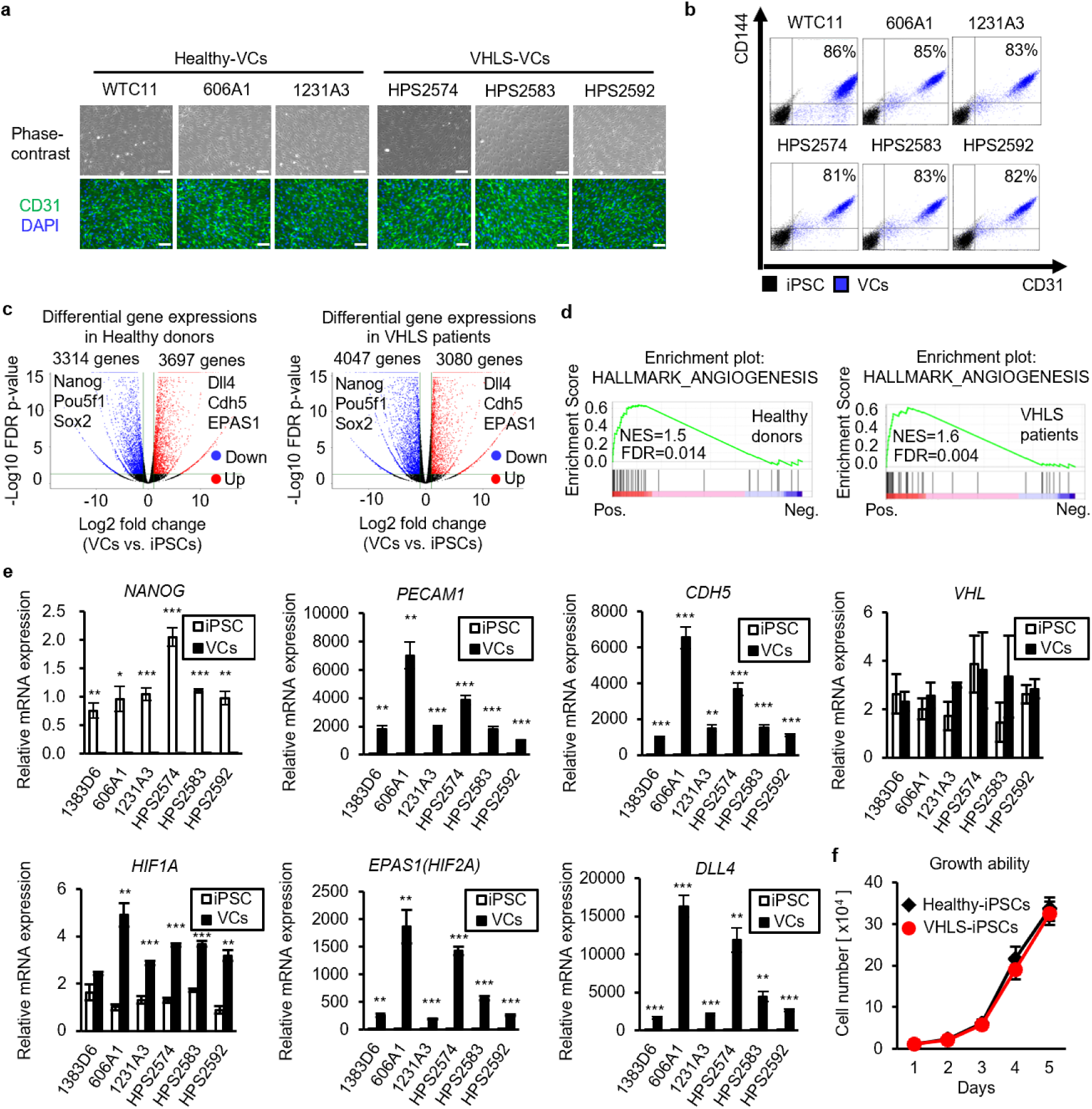
VCs differentiation derived from Healthy-iPSCs and VHLS-iPSCs. a, Representative phase and immunocytochemistry images of CD31^+^ VCs differentiated from Healthy-iPSCs (WTC11, 606A1, and 1231A3) and VHLS-iPSCs (HPS2574, HPS2583, and HPS2592) on day 11. Scale bars, 200 μm. b, Flow cytometric analysis for detecting vascular endothelial markers CD31 and CD144 in the VCs. c, Left, volcano plot showing mRNA expression comparison between Healthy-VCs (1383D6, 606A1, and 1231 A3) and Healthy-iPSCs (1383D6, 606A1, and 1231A3) (baseline) based on RNA-seq. The red color indicates upregulated genes and the blue color indicates down-regulated genes in Healthy-VCs (|Log2 Fold change) > 1, FDR p-value < 0.05). Right, volcano plot showing mRNA expression comparison between VHLS-VCs (HPS2574, HPS2583, and HPS2592) and VHLS-iPSCs (HPS2574, HPS2583, and HPS2592) (baseline) based on RNA-seq. The red color indicates upregulated genes and the blue color indicates downregulated genes in VHLS-VCs (|Log2 Fold change) > 1, FDR p-value < 0.05). d, HALLMARKANGIOGENESIS-related pathway was enriched in Healthy-VCs or VHLS-VCs compared to these IPSCs in GSEA (FDR q-value < 0.05). NES: Normalized enrichment score, e, Expression of a *VHL-gene,* self-renewal marker *(NANOG),* vascular endothelial markers *(PECAM1, CDH5, HIF1A, EPAS1, and DLL4)* were evaluated in the iPSCs and VCs from Healthy donors (1383D6, 606A1, and 1231A3) and VHLS patients (HPS2574, HPS2583, and HPS2592) by real-time qPCR. f, Cell numbers were counted for growth curves in the iPSCs derived from three healthy donors and VHLS patients (n=2). P-values were determined by a two-tailed student’s test (*P < 0.05, **P < 0.01, ***P < 0.001).

**Extended Data Figure 4.**
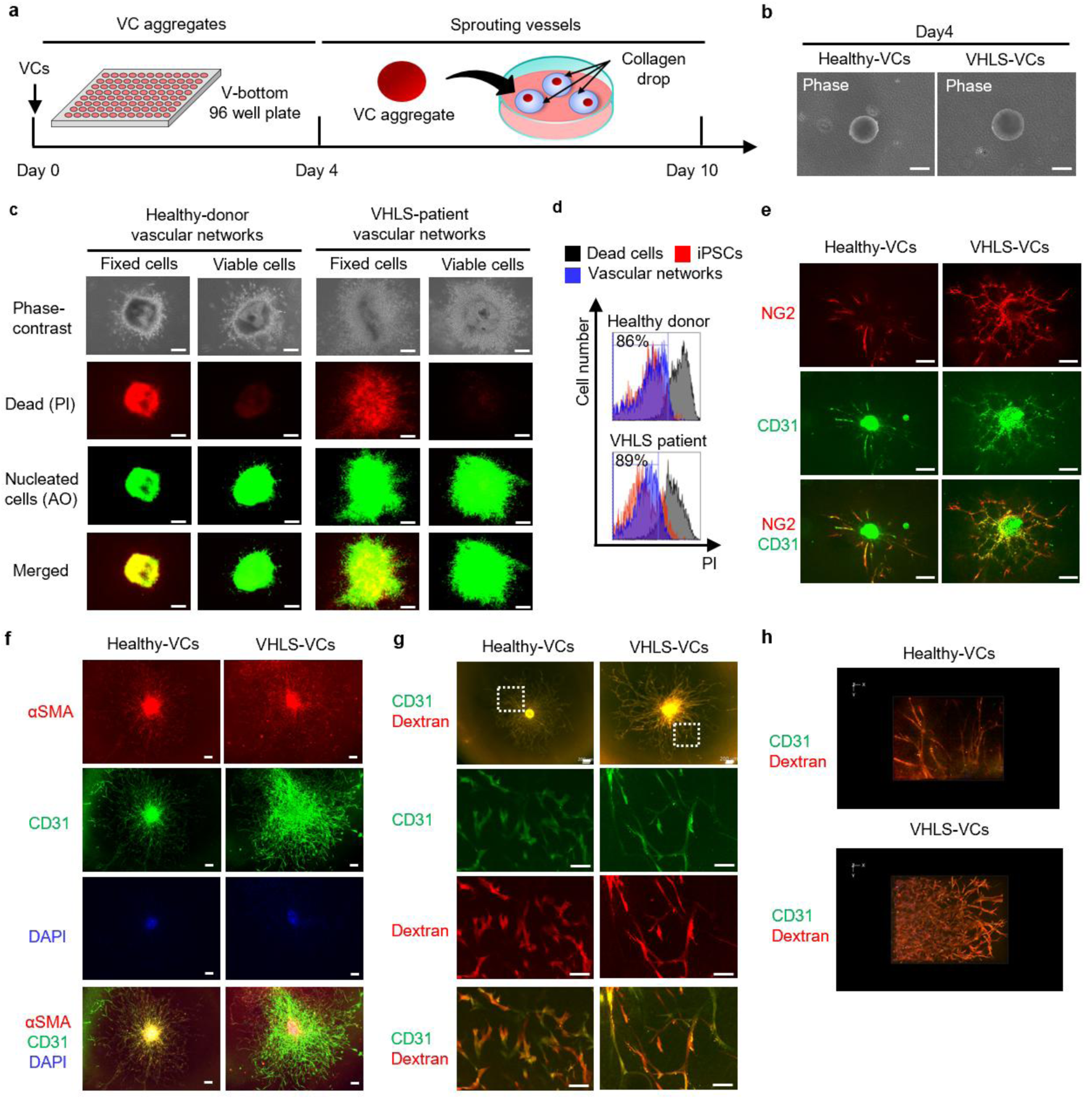
Generation of the vascular networks using VCs. a, Schematic of vascular networks from VCs. Aggregates of VCs were formed for 4 days in floating conditions, then transferred into a collagen drop on the bottom of a culture dish and cultured for 6 days, b, Representative phase images of VCs-derived aggregates from Healthy-VCs and VHLS-VCs at day4. Scale bars, 200 μm. c, Live-dead cell viability images of vascular networks from the Healthy-VCs and VHLS-VCs. Dead cells were stained with propidium Iodide (red), and nucleated cells were acridine orange (AO) positive (green). Fixed cells were used as positive controls for Dead cells in the vascular networks, d, Flow cytometric analysis for detecting dead cells (PI positive) in the vascular networks from the Healthy-VCs and VHLS-VCs (blue). The IPSCs were used to determine live (red)-dead (black) state e, f, ICC images of a vascular endothelial marker CD31 (green), pericyte marker NG2 (red), and smooth muscle marker αSMA (red) in Healthy- and VHLS-derived vascular networks. Nucleus were stained DAPI (blue). Scale bars, 200 μm. g, h, Fluorescence microscopy imaging of the CD31^+^ VCs (green) treated with red fluorescent protein (RFP)-dextran in 2D (g) and 3D (h) morphology. Scale bars, 200 μm (top) and 100 μm (bottom) in g.

**Extended Data Figure 5.**
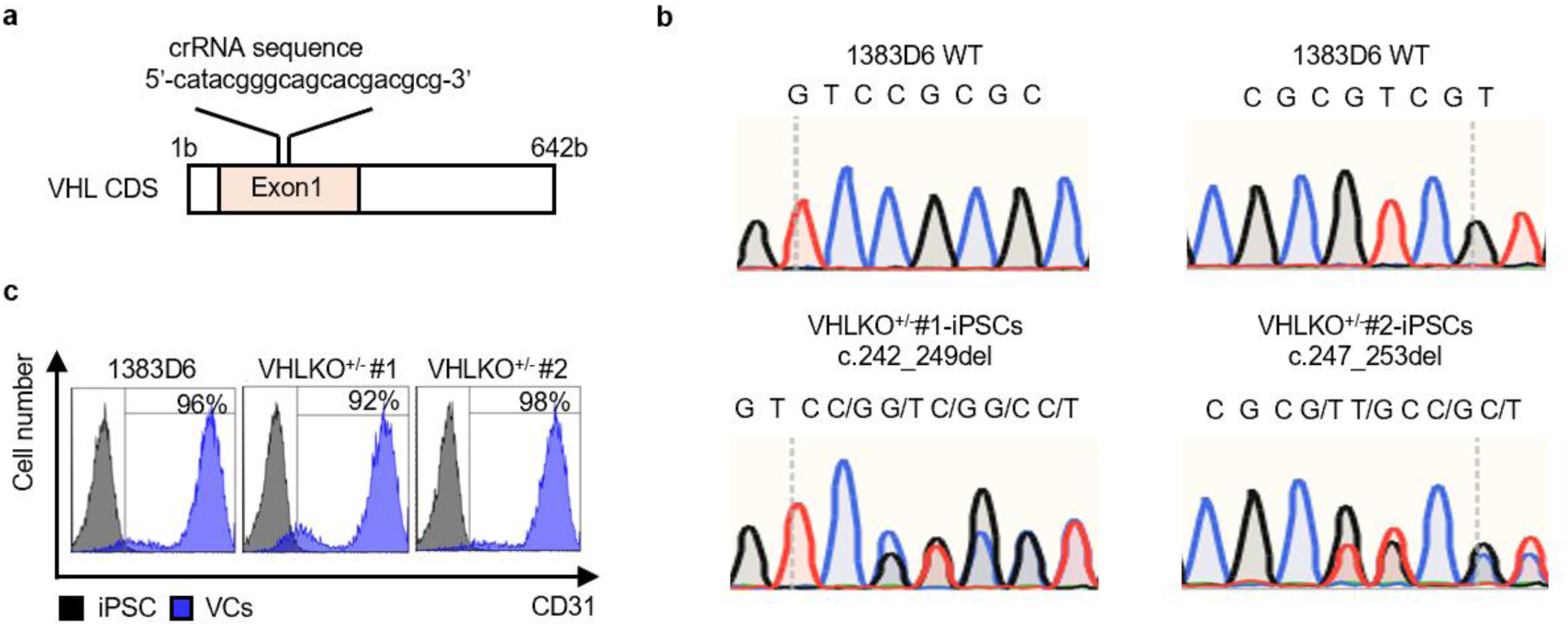
Construction of VHLKO^+/-^ iPSCs from healthy-iPSCs. a, Schematic of the crRNA targeting Exonl in *VHL-gene* in Healthy-iPSCs (1383D6). b, Sequencing analysis of *VHL-*genes in 1383D6, VHLKO^+/-^ #1 (c.242_249del), and VHLKO^+Í^-#2 (c.247_253del). c, Flow cytometric analysis for detecting a vascular endothlial marker CD31 in the 1383D6 and VHLKO^+/-^-VCs.

**Extended Data Figure 6.**
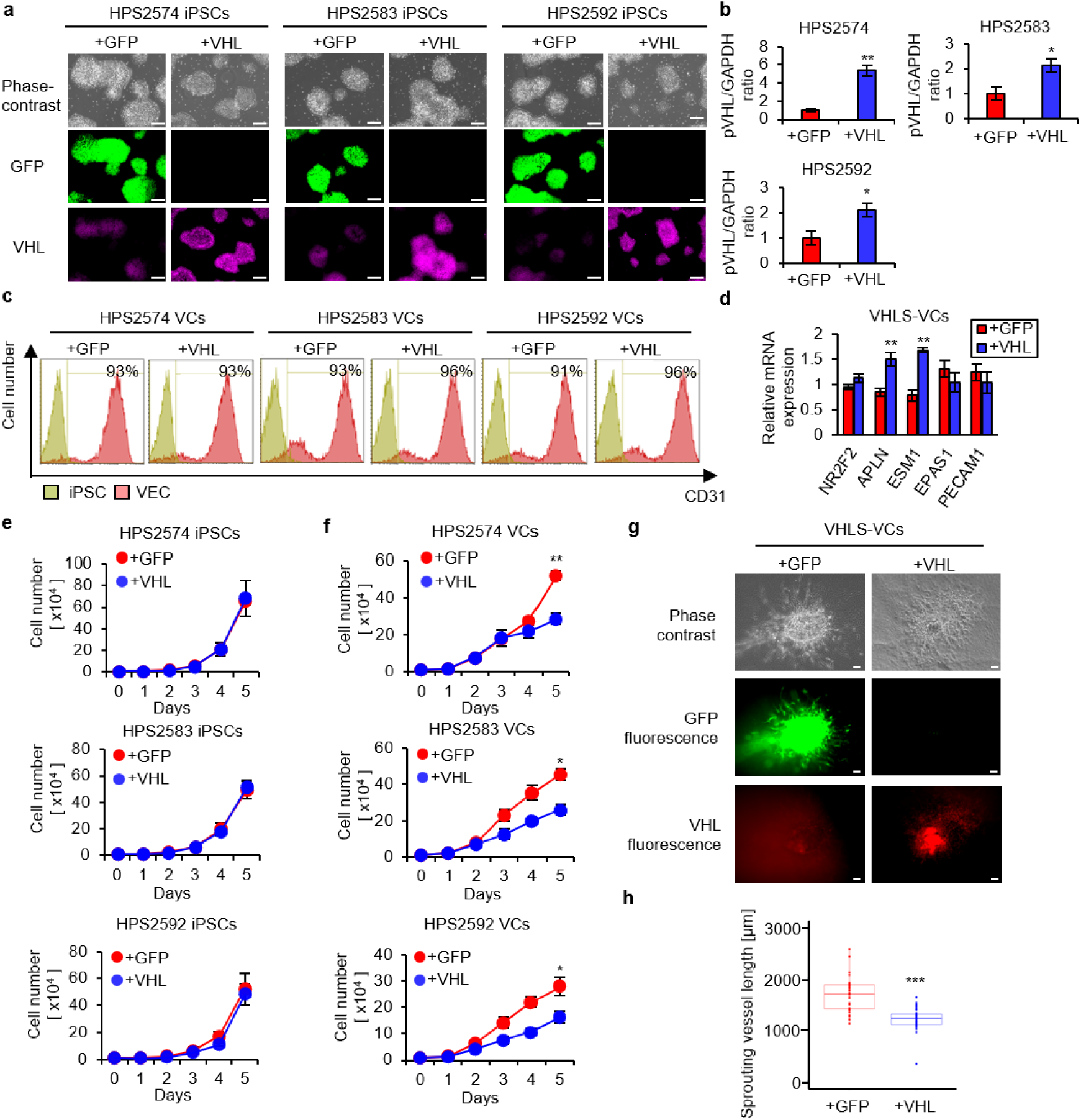
Vasculogenesis is suppressed by VHL-overexpression in VHLS-VCs. a, Representative phases, GFP, and VHL fluorescent images of generated VHLS-iPSCs (HPS2574, HPS2583, HPS592) expressing lentivirus vectors encoding *GFP*or *VHL* genes, b, Relative amounts of VHL proteins (pVHL/GAPDH) in the iPSCs were quantified using WES. Data are shown as the mean = SEM (n = 3). P-values were determined by a two-tailed student’s t-test (*P < 0.05, **P < 0.01). c, Flow cytometric analysis for detecting the CD31 ^+^ VCs. d, Expression of a vein marker (NR2F2), Artery markers (APLN and ESM1), and vascular endothelial markers (EPAS1 and PECAM1) were evaluated in the VHLS-VCs (HPS2574) by real-time qPCR [GFP n=3 and VHL n=3). e, Cell numbers were counted for growth curves in the IPSCs derived from three VHLS patients (GFP n=3 and VHL n=3). f, Cell numbers were counted for growth curves in the VCs derived from three VHLS patients (GFP n=3 and VHL n=3). g, Representative phases and GFP, and VHL fluorescent images of vascular networks expressing *GFP* or *VHL* genes. Scale bars, 200 μm. h, Sprouting vessel lengths from the central core of VC aggregate in the vascular networks quantified by ImageJ software (GFP n=32 and VHL n=36). P-values were determined by a two-tailed student’s t-test (***P < 0.001).

**Extended Data Figure 7.**
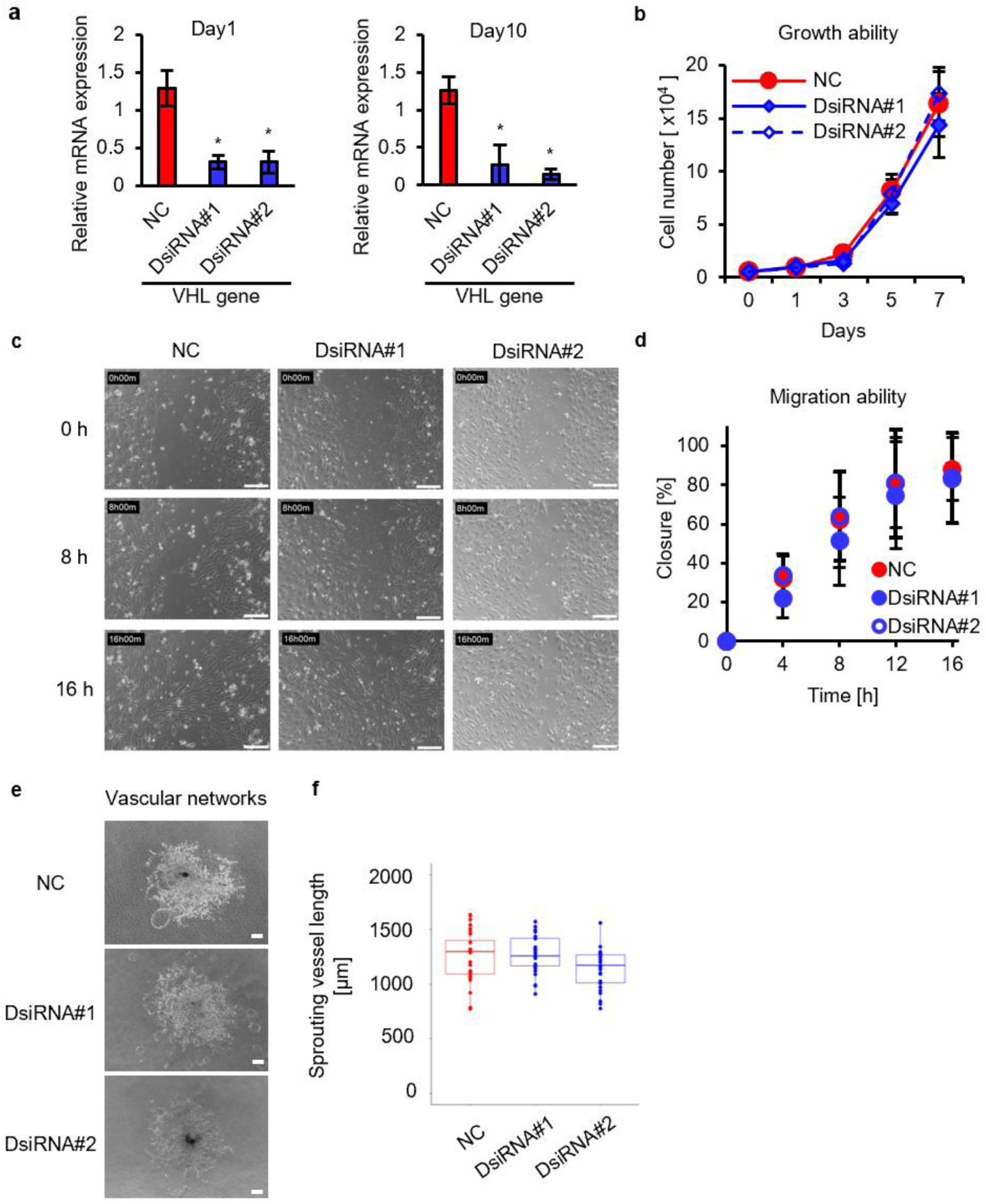
Heterozygous VHL inactivation is sufficient to drive the excessive vasculogenesis in VHLS-VCs. a, The expression of VHL gene was evaluated in VCs from a VHLS patient line, HPS2574, transduced with dicer-substrate short interfering RNAs (Negative control n=3, DsiRNAs#1 n=3, and DsiRNAs#2 n=3) on day1 (Left) and day10 (Right) with qRT-PCR. b, Growth curves were generated from counting cell numbers in the VCs (NC n=3, DsiRNA#1 n=3, and DsiRNA#2 n=3). c, Representative images of the VHLS-VCs with NC, DsiRNA#1, and DsiRNA#2 in wound healing assay at 0, 8, and 16 hours after scratching culture substrates, d, The ratio of closure, determined at consistent locations, assessed for the VHLS-VCs at 0, 4, 8, 12, and 16 hours after scratch (NC n=3, DsiRNA#1 n=3, and DsiRNA#2 n=3). Percentages (%) are shown in the graph, e, Phase-contrast images of blood vessels sprouting from the central core of VC aggregate in the VHLS-VCs with NC, DsiRNA#1, and DsiRNA#2 derived vascular networks. Scale bars, 200 μm. f, Sprouting vessel lengths from the central core of VC aggregate in the vascular networks (NC, DsiRNA#1, DsiRNA#2 n=28). Data are shown as mean ± SEM. P-values were determined by one-way ANOVA Dunnett’s test. * indicates P < 0.05.

**Extended Data Figure 8.**
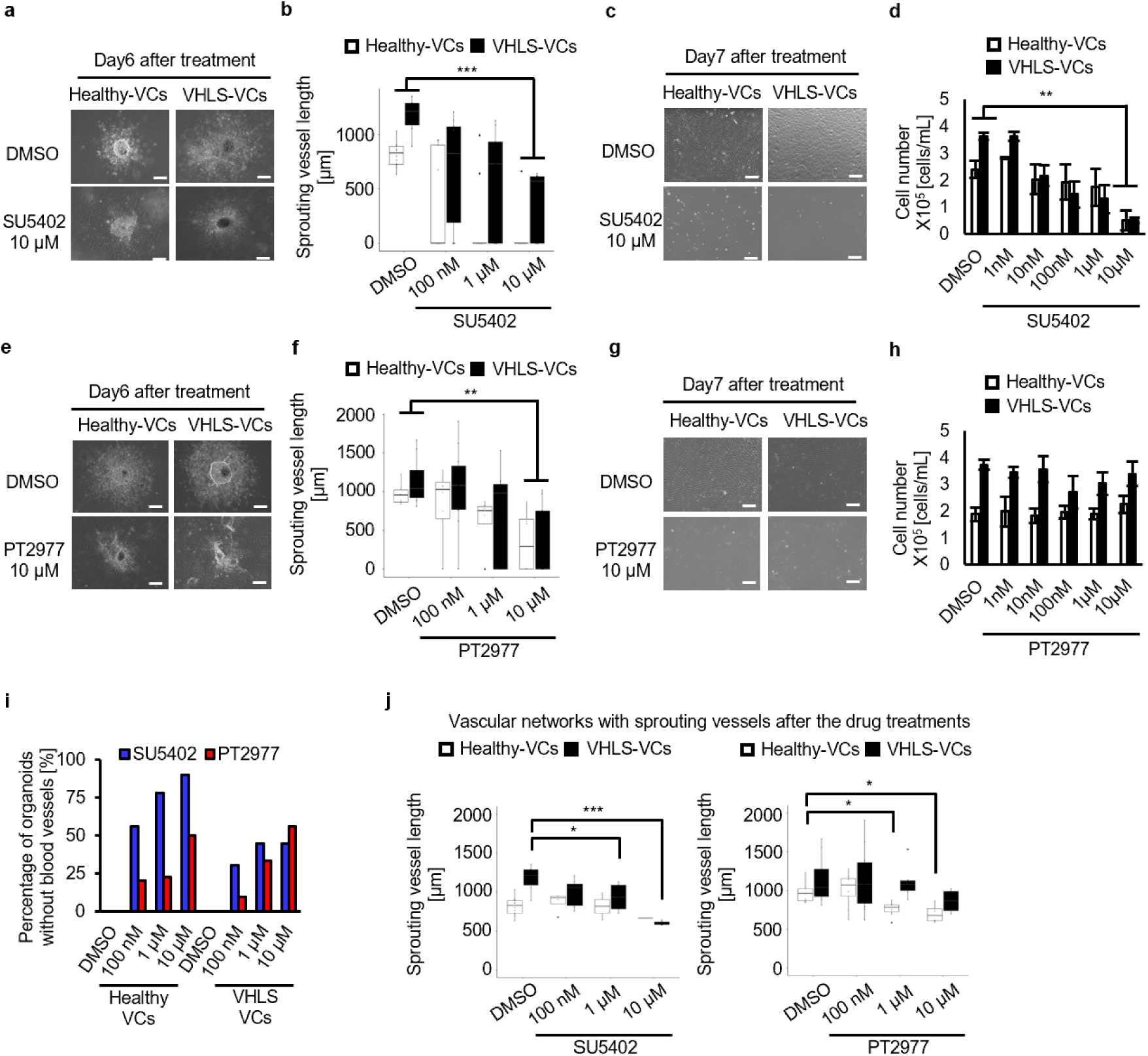
Inhibitory effects of pharmacological inhibitors on Healthy-VCs and VHLS-VCs for vasculogenesis. a, Phase images of sprouting blood vessels in the vascular networks derived from Healthy-VCs and VHLS-VCs with DMSO or SU5402 treatment, an inhibitor for VEGFR and FGFR, from day4 to day 10 at 10 μm. Scale bars, 200 μm. b, Sprouting vessel lengths from the center of these vascular networks (DMSO: Healthy-VCs n=14, VHLS-VCs n=7, SU5402 100 nM: healthy-VCs n=9, VHLS-VCs n=10, SU5402 1 μM: Healthy-VCs n=9, VHLS-VCs n=9, SU5402 10pM: Healthy-VCs n=10, VHLS-VCs n=9). c. Phase images of Healthy-VCs and VHLS-VCs treated with/without SU5402 treatment from day 1 to day7 in monolayer cultures at 10 μM. Scale bars, 200 μm. d, Cell numbers in the VCs were counted after DMSO or SU5402 treatment at 1 nM, 10 nM, 100 nM, 1 μM, and 10 μM for 7days in monolayer cultures (Healthy-VCs n=3, VHLS-VCs n=3). e, Phase images of sprouting blood vessels in the vascular networks derived from Healthy-VCs and VHLS-VCs with/without PT2977 treatment, an inhibitor for HIF2A, from day4 to day 10 at 10 μM. Scale bars, 200 μm. f, Sprouting vessel lengths from the center of vascular networks (DMSO: Healthy-VCs n=9, VHLS-VCs n=8, PT2977 100 nM: Healthy-VCs n=10, VHLS-VCs n=11, PT2977 1 μM: Healthy-VCs n=9, VHLS-VCs n=9, PT2977 10pM: Healthy-VCs n=8, VHLS-VCs n=9) quantified by Imaged software, g, Phase images of Healthy-VCs and VHLS-VCs treated with/without PT2977 treatment from day 1 to day 7 in monolayer cultures at 10 μM. Scale bars, 200 μm. h, Cell numbers in the VCs were counted after DMSO or PT2977 treatment at 1 nM, 10 nM, 100 nM, 1 μM, and 10 μM for 7 days in monolayer cultures (Healthy-VCs n=3 and VHLS-VCs n=3). i, Percentages of the vascular networks without sprouting vessels in all healthy-VCs derived vascular networks (blue) and VHLS-VCs derived vascular networks (red) treated with SU5402 or PT2977. j. Sprouting vessel lengths of vascular networks remaining the vessels after the treatment of SU5402 (left) or PT2977 (right) in (I). Sprouting vessel length from the center of vascular networks (SU5402, DMSO: Healthy-VCs n=12, VHLS-VCs n=7, SU5402 100 nM: healthy-VCs n=4, VHLS-VCs n=7, SU5402 1 μM: Healthy-VCs n=2, VHLS-VCs n=5, SU5402 10pM: Healthy-VCs n=1, VHLS-VCs n=5, PT2977, DMSO: Healthy-VCs n=9, VHLS-VCs n=8, PT2977 100 nM: healthy-VCs n=8, VHLS-VCs n=10, PT2977 1 μM: Healthy-VCs n=7, VHLS-VCs n=6, PT2977 10pM: Healthy-VCs n=4, VHLS-VCs n=4) quantified by Imaged software. Data are shown as mean±SEM. P-values were determined by one-way ANOVA Dunnett’s test (*P < 0.05, *\*\*P* < 0.01, and *\*\*\*P<* 0.001).

**Extended Data Figure 9.**
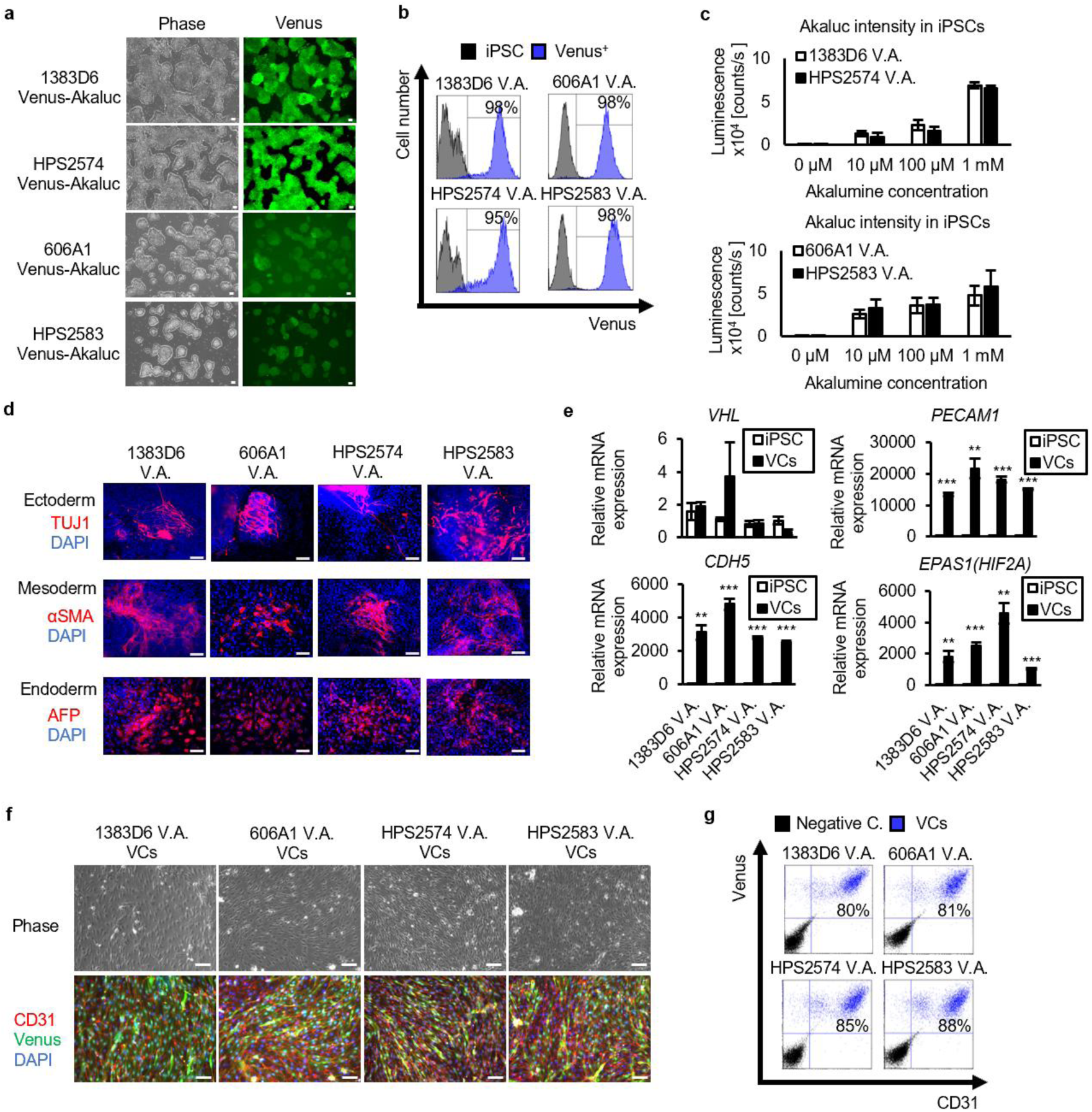
Generation of Healthy-iPSCs and VHLS-iPSCs expressing Venus-Akaluc genes. a, Representative phases and fluorescent images of Venus^+^ iPSCs from Healthy donors (1383D6 and 606A1) and VHLS patients (HPS2574 and HPS2583) after FACS sorting. Scale bars, 200 μm. b, Flow cytometric analysis detecting for Venus proteins in the iPSCs expressing Venus-Akaluc fusion genes, c, Venus-Akaluc^+^ iPSCs were treated with AkaLumine-HCI at 10 μM, 100 μM, and 1 mM (n=3). After 10 min, Akaluc-derived luminescent intensity was measured by spark multimode microplate reader (exposure time = 10 sec.), d, Pluripotency in VHLS-iPSCs was evaluated with in vitro embryoid body (EB) formations. TUJ1 (ectoderm marker), alpha-SMA (mesoderm marker), and AFP (endoderm marker) expressions were used for differentiation markers of three germ layers in EB. Scale bar, 100 μm. e, Expression of a VHL-gene and vascular endothelial markers (PECAM1, CDH5, and EPAS1) were evaluated in the Venus-Akaluc^+^ iPSCs and VCs from Healthy donors (1383D6 and 606A1) and VHLS patients (HPS2574 and HPS2583) by real-time qPCR. f, Representative phase-contrast and immunocytochemistry images of CD31 (red) and Venus (green) proteins in VCs differentiated from the Venus-Akaluc^+^ iPSCs. Nucleus was stained with DAPI (blue). Scale bars, 200 μm. g, Flow cytometric analysis for detecting venus and vascular endothelial marker CD31 in the VCs and iPSCs (Negative control).

**Extended Data Figure 10.**
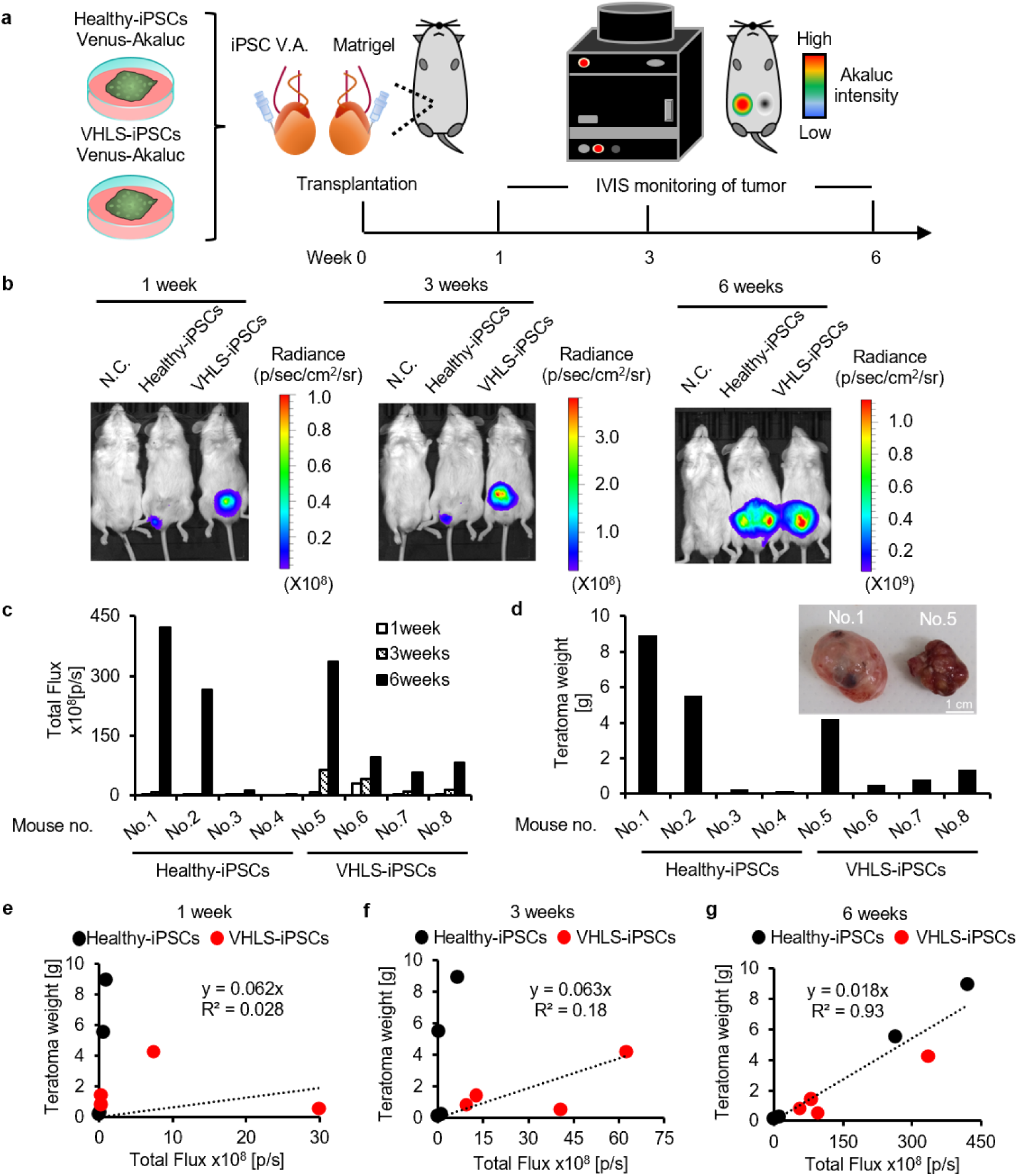
Monitoring mice transplanted with Venus-Akaluc^+^ iPSCs during teratoma formation. a, Schematic of monitoring Venus-Akaluc^+^ iPSCs transplanted into testis in mice with I VIS imaging system. 1.0×10^6^ cells of Healthy-iPSCs (1383D6) or VHLS-iPSCs (HPS2574) expressing Venus-Akaluc fusion genes were resuspended in 50% matrigel and injected into one testis, and matrigel without cells in the other as control. The Venus-Akaluc^+^ iPSCs were imaged at 1,3, and 6 weeks after the transplantation with I VIS. b, Representative photograph and luminescence images of Akaluc-derived signals in the mice at 1, 3, and 6 weeks. Approximately 10 min before the imaging, the mice were intraperitoneally administered with 100 μL of 30 mM AkaLumine-HCI, and the images were acquired. Akaluc-intensity was indicated by radiance value (p/sec/cm^2^/sr). c, Total flux (p/s) was calculated from the Akaluc-intensity with IVIS (Healthy-iPSCs n=4, VHLS-iPSCs n=4). d, Tumor weight was measured after sacrificing mice at 6 weeks, e-g, Correlation of tumor weight and Akaluc-intensity (Total flux) at 1 week (e), 3 weeks (f), and 6 weeks (g) after the transplantation. Spearmann’s correlation coefficient is depicted.

**Extended Data Figure 11.**
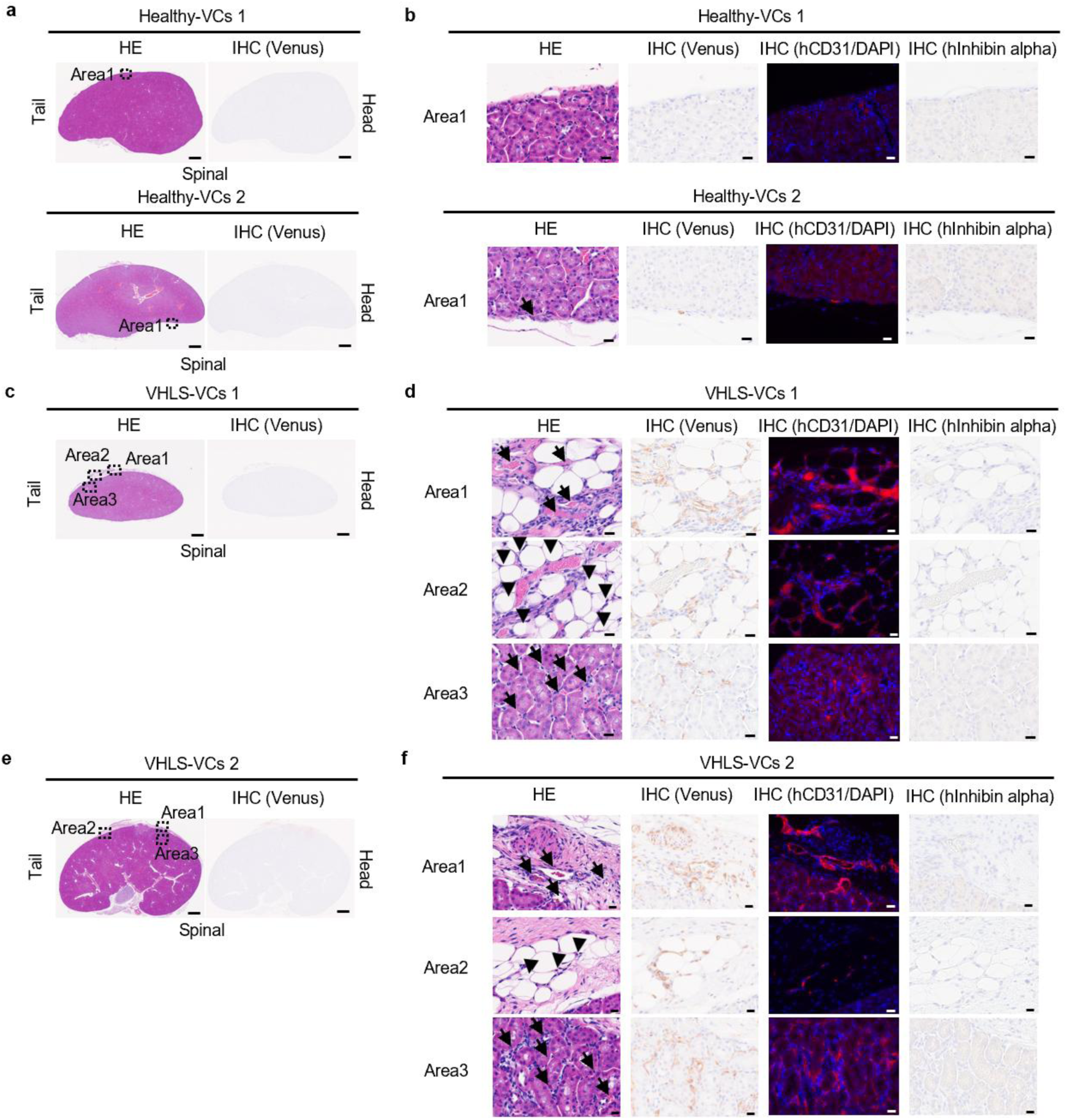
Pathological analysis of kidney sections transplanted with Healthy-VCs and VHLS-VCs. a, b, Transplanted kidneys with Venus-Akaluc^+^ Healthy-VCs (1383D6 and 606A1) were isolated and analyzed by hematoxylin-eosin staining (HE) and immunohistochemistry (IHC) to detect Venus^+^ human cells. Venus^1^ cells were indicated by arrows. Scale bars, 1mm (whole kidney), 25 μm (Areal), c-f, Transplanted kidneys with Venus-Akaluc^+^ VHLS-VCs (HPS2574 and HPS2583) were isolated and stained with HE and IHC (hCD31 and hlnhibin alpha). Venus^1^ blood vessels were indicated by arrows (Areal and Area3). Venus^+^hCD31^+^ stromal cells were indicated by arrowheads (Area2). Scale bars, 1mm (whole kidney), 25 μm (Areal, Area2, and Area3). Experiments were repeated independently 2 times with similar results.

**Extended Data Figure 12.**
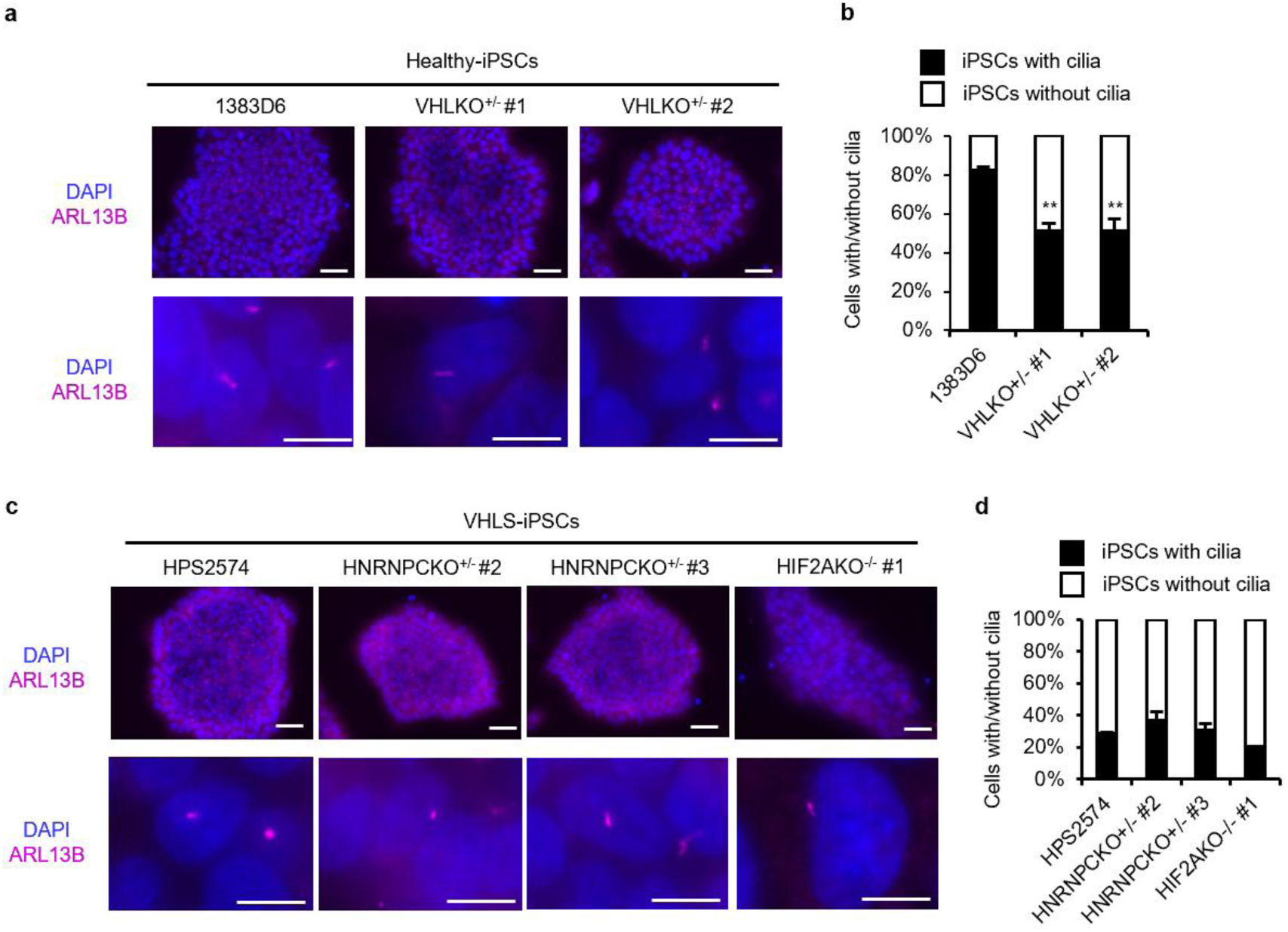
Loss of ciliated cells were in VHLKO^+/-^ iPSCs or VHLS-iPSCs. a,c, Endogeneous expression of a cilia marker ADP-ribosylation factor (Arf)-like protein 13B (ARL13B) (purple) in healthy-IPSCs (1383D6), VHLKO^+/-^ iPSCs (#1 and #2), VHLS-iPSCs (HPS2574), HNRNPCKO^+/-^ iPSCs (#2 and #3), and HIF2AKO^+^-iPSCs (#1). Nucleus was stained with DAPI (blue). Scale bar, 50 μm (Top), 10 μm (Bottom). b,d, Frequency of the IPSCs with/without cilia formation (n=3). P-values were determined by one-way ANOVA Tukey‘s test (**P < 0.01).

**Extended Data Figure 13.**
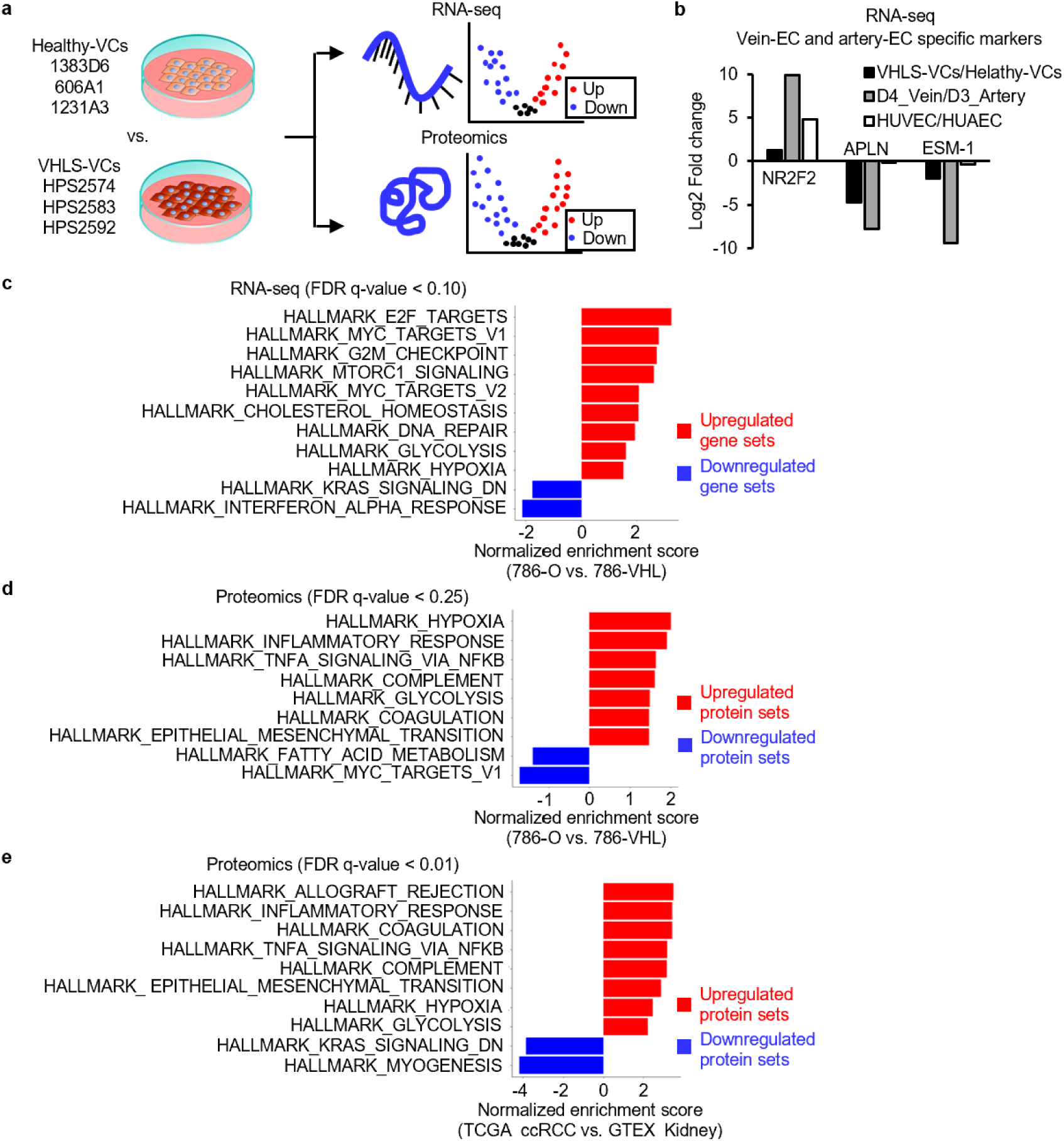
RNA-seq or proteomics analysis in arteriovenous cell lines, 786-0 ccRCC lines, and ccRCC patients, a, Experimental flow for RNA-seq and proteomics in VCs derived from Healthy-iPSCs (1383D6, 606A1, and 1231 A3) and VHLS-iPSCs (HPS2574, HPS2583, and HPS2592). b, Gene expression ratio of vein marker *(NR2F2)* and artery markers *(APLN, ESMI)* between VHLS-VCs and Healthy-VCs (baseline). As a control, the ratios in IPSCs-derived vein on day4/artery on day3 or human umbilical vein endothelial cells (HUVEC)/ human umbilical artery endothelial cells (HUAEC) were calculated using previous data (Lay Teng Ang et al., 2022.; Samir Sissaoui et al., 2020.). c, GSEA of upregulated genes or downregulated genes in VHL-deficient 786-0 ccRCC line compared to VHL re-expression line in 786-0 (786-VHL) (baseline). Red bars show gene data sets enriched in VHL-deficient 786-0, blue bars are enriched gene data sets in 786-VHL (FDR q-value < 0.10). d, GSEA of upregulated proteins or downregulated proteins between VHL-deficient 786-0 line and 786-VHL (baseline). Red bars show protein data sets enriched in VHL-deficient 786-0, blue bars are enriched protein data sets in 786-VHL (FDR q-value < 0.25). e, Enriched pathways in GSEA between TCGA Kidney Clear Cell Carcinoma patients (n = 531) and healthy GTEx (n = 28) cohorts (baseline) using proteomics data. Red bars show protein data sets enriched in ccRCC patients, Blue bars are enriched protein data sets in healthy donors (FDR q-value < 0.01).

**Extended Data Figure 14.**
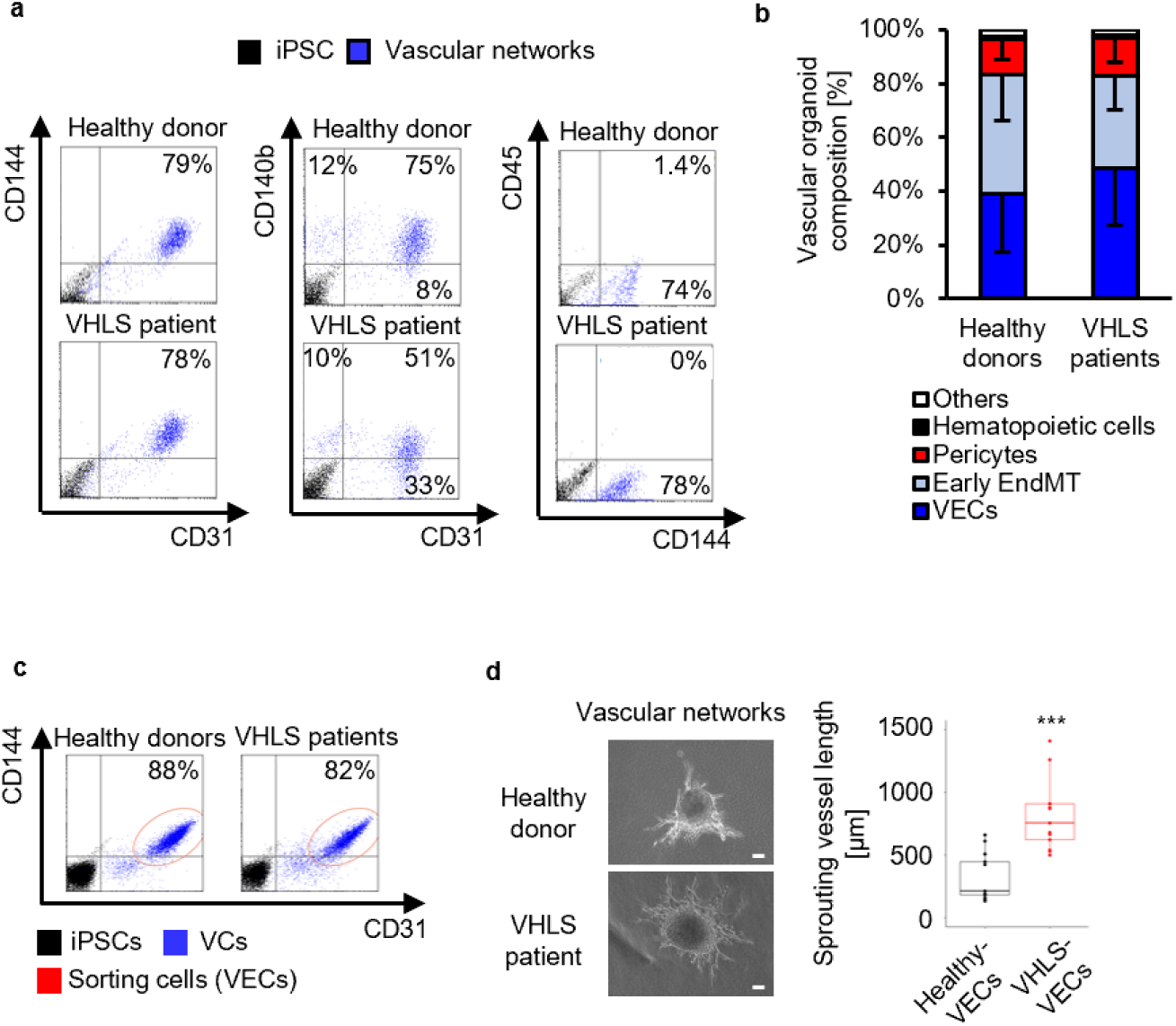
Characterization and purification of vascular cells (VCs) constructing vascular networks a, b, Flow cytometric analysis to characterize key cellular components of the vascular networks derived from Healthy-VCs (1383D6, 606A1, and 1231 A3) and VHLS-VCs (HPS2574, HPS2583, and HPS2592). Vascular endothelial cells (VECs) were defined as CD31^+^CD144^+^, VECs in Early EndMT state as CD31^+^PDGFRβ^+^, pericytes as CD3TPDGFRβ^÷^, and hematopoietic cells by CD45^+^. In the right panels, bar graph shows the relative populations of the VECs, Early EndMT, pericyte, and hematopoietic cells. Data are shown as mean ± SEM. Experiments were independently repeated 3 times, c, Flow cytometric analysis for sorting vascular endothelial cells (CD31^+^CD144^+^) in VCs differentiated from healthy-donors and VHLS-patients. Sorting cells were indicated by a red circle in the plot, d, Left, Phase contrast images of vascular networks from FACS-isolated VECs from the VCs. Scale bars, 100 μm. Right, Sprouting vessel lengths from the central core of vascular networks (Healthy donors; WTC11, 606A1, 1231 A3, n=13, VHLS-patients; HPS2574, HPS2583, HPS2592, n=13). P-values were determined by a two-tailed student‘s test (***P < 0.001).

**Extended Data Figure 15.**
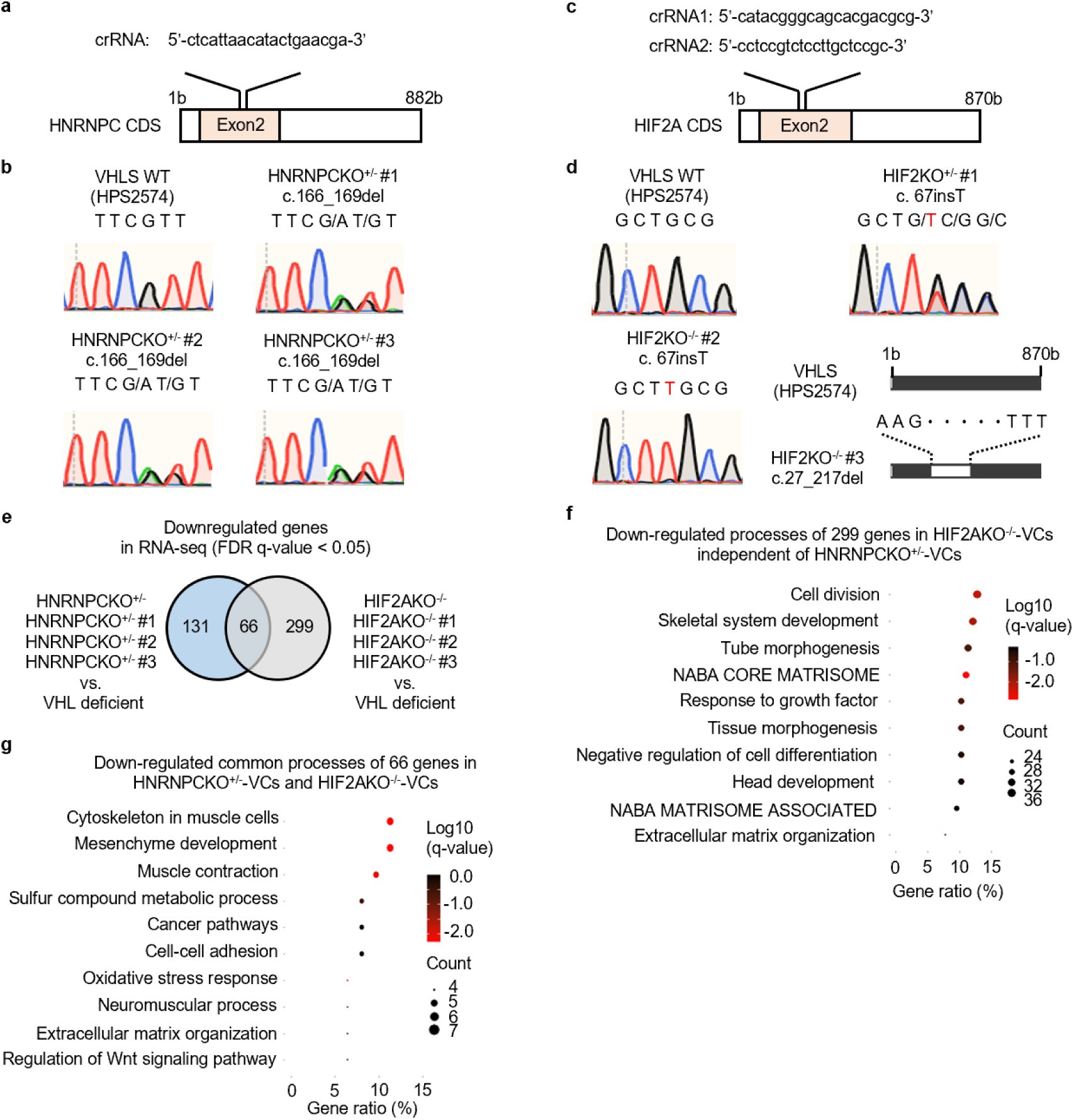
Construction of HNRNPCKO^+/-^ IPSCs and HIF2AKO^-/-^ iPSCs from VHLS-iPSCs. a,c, Schematics of the crRNAs targeting Exon2 in *HNRNPC-gene* or *HIF2A-gene* in VHLS-iPSCs (HPS2574). b,d, Sequencing analysis of *HNRNPC-gene* or *HIF2A-gene* in HPS2574, HNRNPCKO^+/-^#1 (c. 166_169del), HNRNPCKO^+/-^ #2 (c. 166_169del), HNRNPCKO^+/-^ #3 (c.166 169del), HIF2AKO^-/-^ #1 (c. 67insT), HIF2AKO^-/-^ #2 (c. 67insT), HIF2AKO^-/-^ #3 (c.27_217del). e, Venn diagram showing the overlap between downregulated genes in HNRNPCKO^+/-^-VCs and HIF2AKO^-/-^ VCs vs. VHL-deficient-VCs (FDR p-value < 0.05). f,g, Gene ontology analysis for the 299 genes specifically downregulated in HIF2AKO^-/-^ lines independent of HNRNPCKO^+/-^ lines (f) or for the 66 genes commonly downregulated in HNRNPCKO and HIF2AKO lines (g) compared to VHL-deficient lines. Dots represent term enrichment, Log_10_ q-values indicating the statistical significance of the enrichment are represented along a gradient color from black (less significant) to red (most significant). The sizes of the dots represent the count of differentially expressed genes belonging to each term.

**Extended Data Figure 16.**
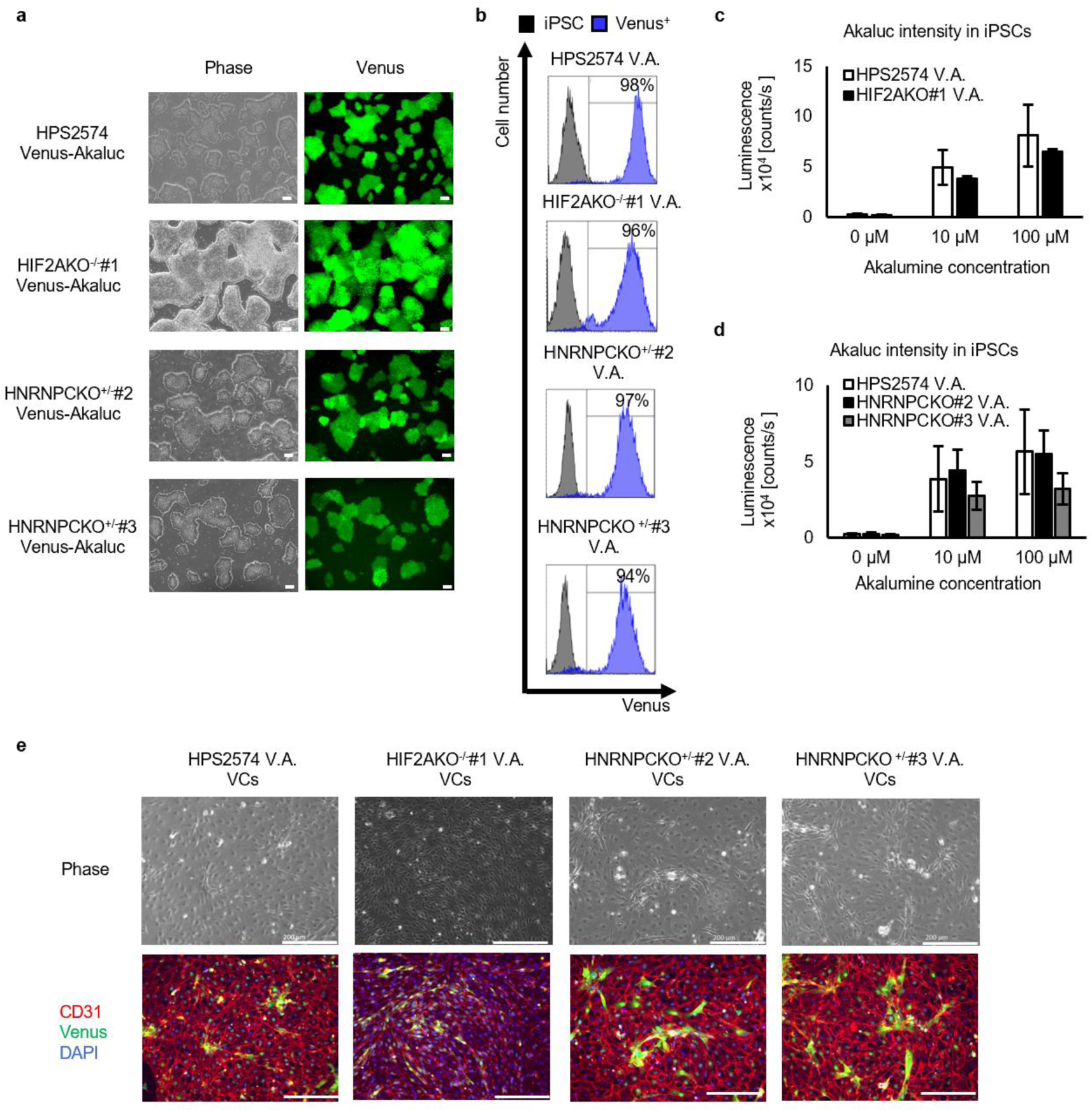
Generation of HNRNPCKO^+A^-iPSCs and HIF2AKO^/^-iPSCs expressing Venus-Akaluc genes. a, Representative phases and fluorescent images of Venus^4^ iPSCs from VHLS WT (HPS2574), HNRNPCKO^+/-^(#2 and #3), and HIF2AKO^+/-^ iPSCs (#1) after FACS sorting. Scale bars, 200 μm. b, Flow cytometric analysis detecting Venus proteins in the iPSCs expressing Venus-Akaluc fusion genes, c, Venus-Akaluc^4^ IPSCs were treated with Akalumine-HCI at 0 μm, 10 μm, and 100 μm (n=3). After 10 min, Akaluc-derived luminescent intensity was measured by spark multimode microplate reader (exposure time = 10 sec.), d, Representative phases and ICC images of CD31 (red) and Venus (green) proteins in VCs differentiated from the Venus-Akaluc^4^ iPSCs. Nucleus was stained with DAPI (blue). Scale bars, 200 μm.

**Extended Data Figure 17.**
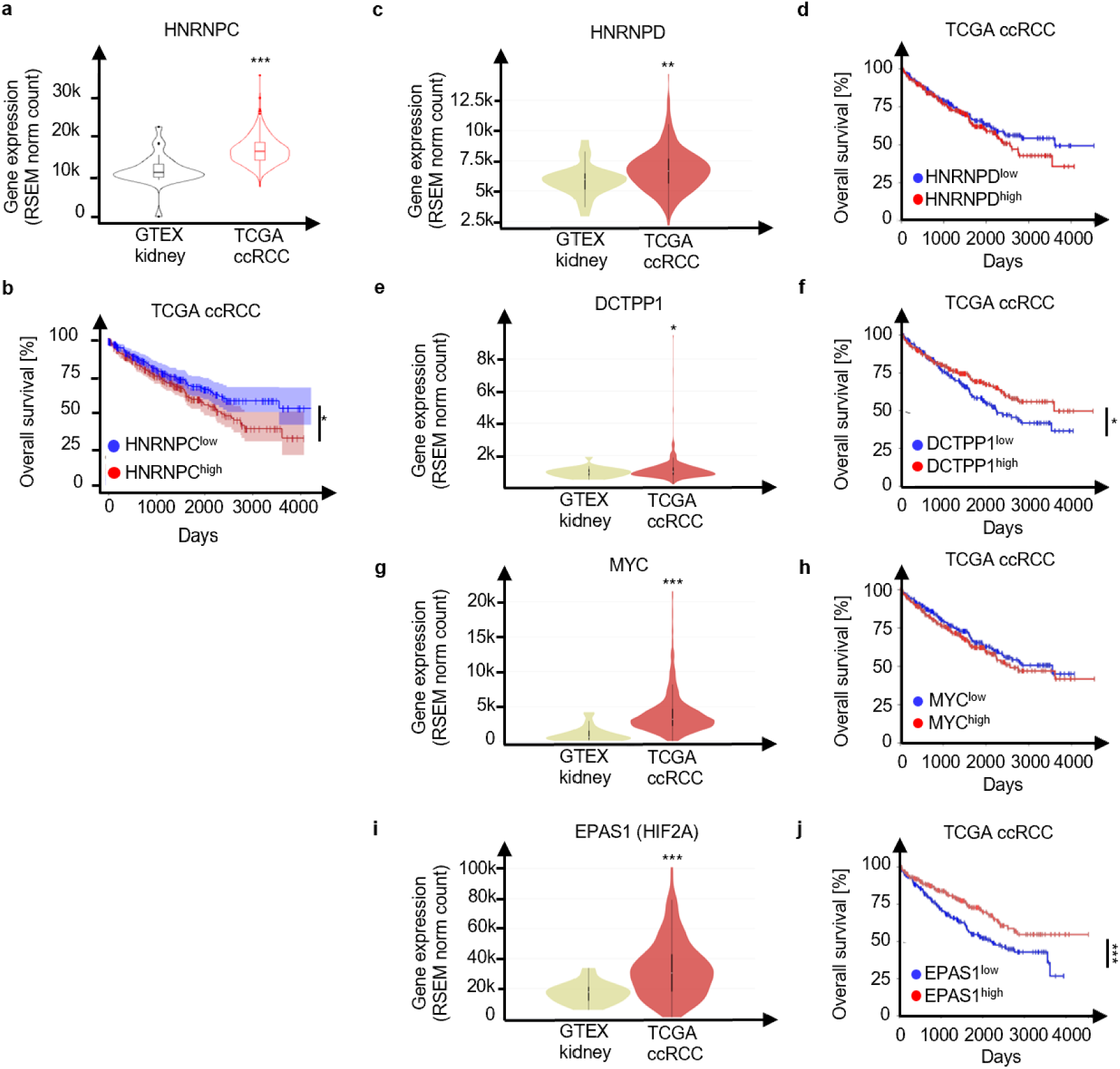
Gene expressions and survival curves on clinical data in clear cell RCC (ccRCC) patients and healthy donors, a, Gene expression (RNA-Seq by Expectation-Maximization (RSEM) normalized count) of *HNRNPC* obtained from data sets in TCGA kidney clear cell carcinoma (n = 531) and healthy GTEx (n = 28) cohorts from UCSC toil RNA-seq recompute on UCSC Xena browser. P values are derived from a two-tailed student’s t test. (***P < 0.001). b, Kaplan-Meier curves showing overall survival of ccRCC patients with high expressions of *HNRNPC* gene (HNRNPC^+^, RSEM normalized count ≥ 13.98) versus patients with low expressions of *HNRNPC* gene (HNRNPC^low^, RSEM normalized count < 13.98). 95% equal precision bands were displayed. P-values are calculated by the two-tailed log-rank test fP < 0.05). c,e,g,i, Gene expression (RNA-Seq by Expectation-Maximization (RSEM) normalized count) of *HNRNPD,. DCTPP1, MYC, EPAS1* obtained from data sets in TCGA kidney clear cell carcinoma (n = 531) and healthy GTEx (n = 28) cohorts from UCSC toil RNA-seq recompute on UCSC Xena browser. P-values are derived from a two-tailed student’s t test. (*\*P <* 0.05,\*\**P* < 0.01, *\*\*\*P <* 0.001). d,f,h,j, Kaplan-Meier curves showing overall survival of ccRCC patients with high expressions of *HNRNPD* gene (HNRNPD^high^, RSEM normalized count ≧ 12.69), *DCTPP1* gene (DCTPP1^high^, RSEM normalized count ≧ 9.949), *MYC* gene (MYC^high^, RSEM normalized count ≧ 11.63), *EPAS1* gene (EPASI^high^, RSEM normalized count ≧ 14.87) versus patients with low expressions of *HNRNPD* gene (HNRNPD^low^, RSEM norm count < 12.69), *DCTPP1* gene (DCTPP1^low^, RSEM norm count < 9.949), A4 YC gene (MYC^low^, RSEM norm_count < 11.63) *EPAS1* gene (EPAS1^low^, RSEM norm_count < 14.87). P values are calculated by the two-tailed log-rank test *\*P<* 0.05, ***P< 0.001).

**Extended Data Figure 18.**
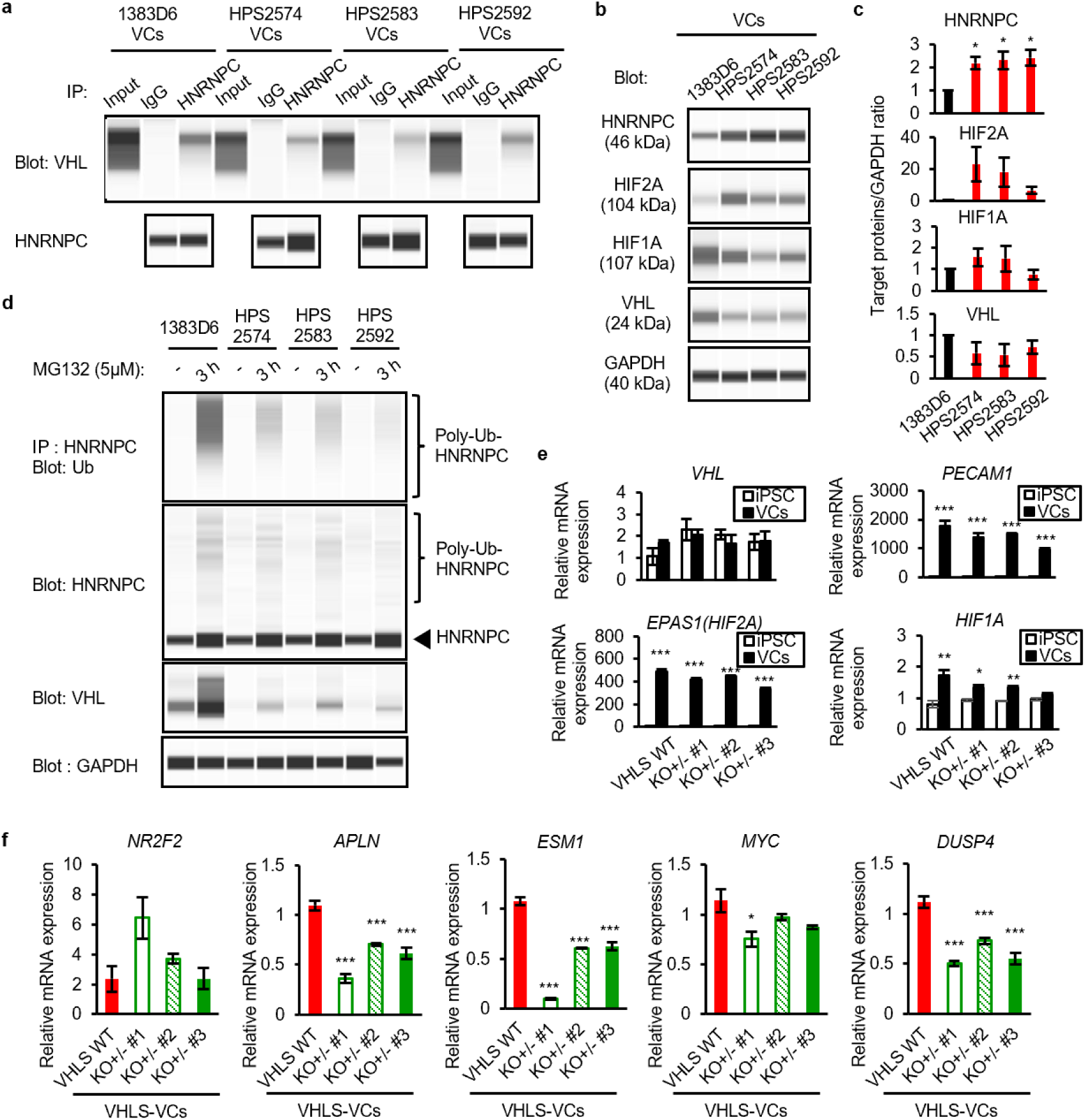
VHL-HNRNPC axis was suppressed with pVHL inactivation via the ubiquitin-proteasome system in VHLS-VCs, contributing to HNRNPC stabilization to maintain transcriptional characteristics for the blood vessel development in VHLS. a, Co-immunoprecipitation (Co-IP) assays of endogenous proteins of HNRNPC or control IgG were performed, b, Western blot analysis of HNRNPC, HIF2A, HIF1 A, VHL, and GAPDH proteins in healthy-VCs (1383D6) and VHLS-VCs (HPS2574, HPS2583, and HPS2592) stimulated with 1% O_2_ for 6 hours, c, Relative amounts of HNRNPC, HIF2A, HIF1A, and VHL proteins normalized by GAPDH using WES. Data are shown as the mean ± SEM (n = 3). d, Ubiquitination of endogenous HNRNPC proteins in Healthy-VCs (1383D6) and VHLS-VCs (HPS2574, HPS2583, and HPS2592) treated with 5 μm MG-132 for 3 hours. Cell lysates were analyzed by immunoprecipitation with an antibody against HNRNPC followed by immunoblotting with an antibody against ubiquitin, and by immunoblotting with antibodies against HNRNPC, VHL, and GAPDH. e, f, Expression of *VHL* gene, vascular endothelial markers (*PECAMt EPAS1(HIF2A)), HIF1A gene* (e), vein marker (*NR2F2),* artery markers (*APLN, ESM1),* and mitogen-activated protein kinase (MAPK)-related regulators (*MYC* and *DUSP4)* (f) were evaluated in the iPSCs or VCs from VHLS WT (HPS2574) and three HNRNPCKO^+/-^ lines derived from VHLS WT (HPS2574) by real-time qPCR. P-values were determined by a two-tailed student‘s test and one-way ANOVA Dunnett’s test (*P < 0.05, **P < 0.01, ***P < 0.001).

**Extended Data Figure 19.**
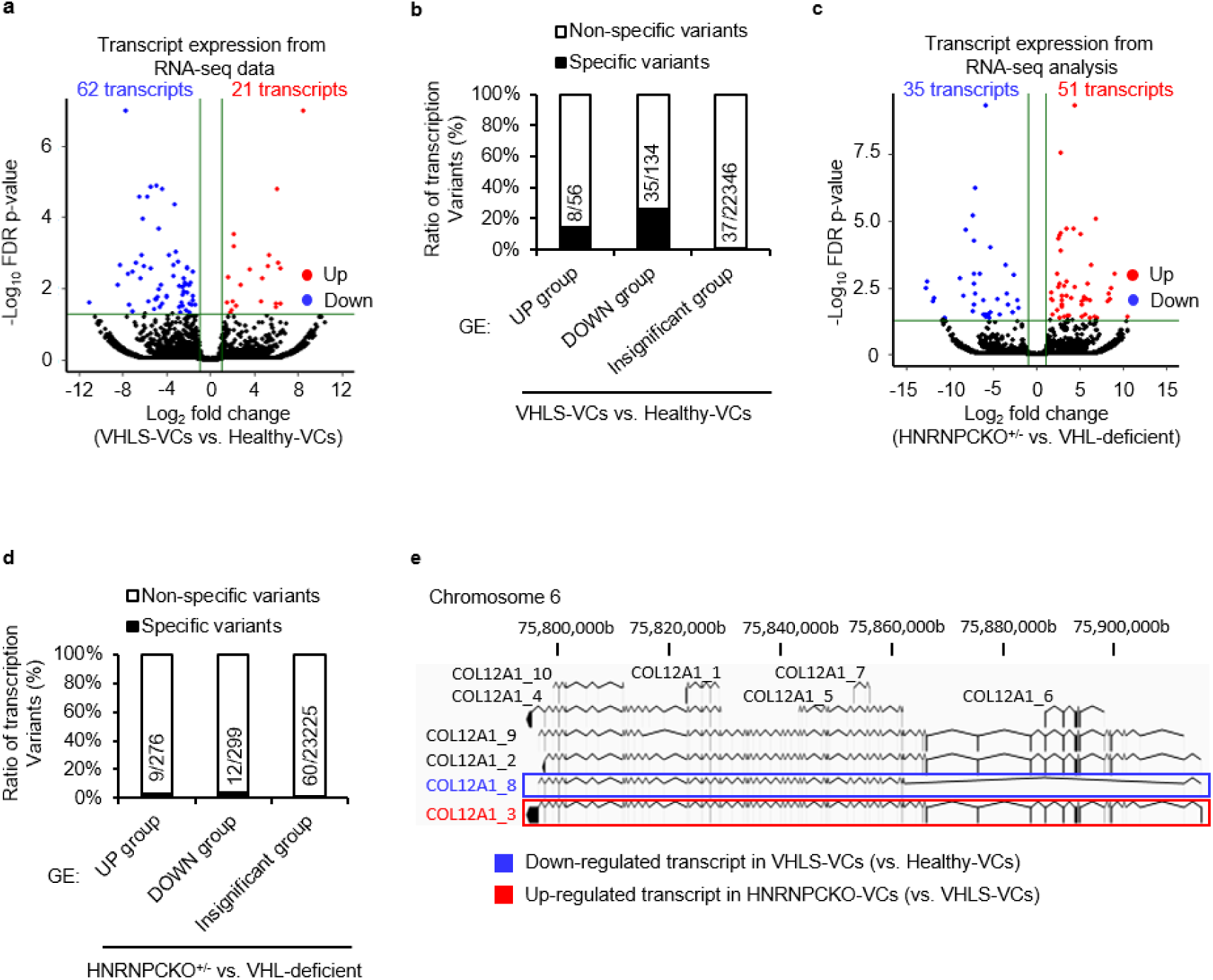
Transcript expression (TE)-based analysis for identifying alternative splicing variants in VHLS-VCs. a, Volcano plot showing the differential expression of the transcript variants between VHLS-VCs (HPS2574, HPS2583, and HPS2592) and healthy-VCs (1383D6, 6060A1, and 1231 A3) (baseline) based on the RNA-seq data. The red circles indicate up-regulated variants, the blue circles indicate down-regulated variants, and the black circles mean insignificant variants in VHLS-VCs (|Log2 Fold change) > 1, FDR p-value < 0.05). b, The ratio of transcript variants specifically upregulated or downregulated in the gene expression groups between VHLS-VCs and healthy-VCs (Figure 3a). The number of transcript variants specifically expressed was indicated in the bar graph (upregulated group; 8 variants/56 genes, downregulated group; 35 variants/134 genes, insignificant group; 37 variants/22346 genes), c, Volcano plot showing the differential expression of the transcript variants on VCs derived from VHL-deficient lines (HPS2574, VHLKO^+/-^ #I, and VHLKO^+/-^ #2) (baseline) and HNRNPCKO^+/-^ lines (HNRNPCKO^+/-^#1, HNRNPCKO^+/-^#2, and HNRNPCKO^+/-^#3). The red color indicates up-regulated variants, the blue color indicates down-regulated variants, and the black plots mean insignificant variants in HNRNPCKO^+/-^-VCs (|Log2 Fold change| > 1, FDR p-value < 0.05). d, Percentage of transcript variants specifically upregulated or downregulated in the gene expression groups between HNRNPCKO^+z^-VCs and VHL-deficient-VCs. The number of transcript variants specifically expressed was indicated in the bar graph (upregulated group; 9 variants/276 genes, downregulated group; 12 variants/299 genes, insignificant group; 60 variants/23225 genes), e, Representative transcript variants in the *COL12A1* gene differentially expressed in VHLS-VCs (compared to Healthy-VCs) or HNRNPCKO^+/-^VCs (compared to VHL-deficient-VCs). COL12A1_8 variant downregulated in the VHLS-VCs was highlighted by blue color, and COL12A13 (encodes the longest isoform) upregulated in the HNRNPCKO^+/-^-VCs was indicated by red color.

## References

1 Glasker, S., Vergauwen, E., Koch, C. A., Kutikov, A. & Vortmeyer, A. O. Von Hippel-Lindau Disease: Current Challenges and Future Prospects. Onco Targets Ther 13, 5669–5690 (2020). 10.2147/OTT.S190753

2 Maher, E. R. & Sandford, R. N. von Hippel-Lindau Disease: an Update. Current Genetic Medicine Reports 7, 227–235 (2019). 10.1007/s40142-019-00180-9

3 Lonser, R. R. et al. von Hippel-Lindau disease. Lancet 361, 2059–2067 (2003). 10.1016/S0140-6736(03)13643-4

4 Bajaj, S., Gandhi, D., Nayar, D. & Serhal, A. Von Hippel-Lindau Disease (VHL): Characteristic Lesions with Classic Imaging Findings. J Kidney Cancer VHL 10, 23–31 (2023). 10.15586/jkcvhl.v10i3.293

5 Maher, E. R., Neumann, H. P. & Richard, S. von Hippel-Lindau disease: a clinical and scientific review. Eur J Hum Genet 19, 617–623 (2011). 10.1038/ejhg.2010.175

6 Senkin, S. et al. Geographic variation of mutagenic exposures in kidney cancer genomes. Nature 629, 910–918 (2024). 10.1038/s41586-024-07368-2

7 Sato, Y. et al. Integrated molecular analysis of clear-cell renal cell carcinoma. Nat Genet 45, 860–867 (2013). 10.1038/ng.2699

8 Kaelin, W. G., Jr. Von Hippel-Lindau disease: insights into oxygen sensing, protein degradation, and cancer. J Clin Invest 132 (2022). 10.1172/JCI162480

9 Maxwell, P. H. et al. The tumour suppressor protein VHL targets hypoxia-inducible factors for oxygen-dependent proteolysis Nature 399, 271–275 (1999). 10.1038/20459.

10 Jonasch, E. et al. Belzutifan for Renal Cell Carcinoma in von Hippel-Lindau Disease. N Engl J Med 385, 2036–2046 (2021). 10.1056/NEJMoa2103425

11 Gnarra, J. R. et al. Defective placental vasculogenesis causes embryonic lethality in VHL-deficient mice. Proceedings of the National Academy of Sciences 94, 9102–9107 (1997). 10.1136/jmg.39.7.463

12 Haase, V. H., Glickman, J. N., Socolovsky, M. & Jaenisch, R. Vascular tumors in livers with targeted inactivation of the von Hippel–Lindau tumor suppressor. Proceedings of the National Academy of Sciences 98, 1583–1588 (2001). 10.1073/pnas.98.4.1583

13 Wang, H. et al. Deletion of the von Hippel-Lindau Gene in Hemangioblasts Causes Hemangioblastoma-like Lesions in Murine Retina. Cancer Res 78, 1266–1274 (2018). 10.1158/0008-5472.CAN-17-1718

14 Pritchett, T. L., Bader, H. L., Henderson, J. & Hsu, T. Conditional inactivation of the mouse von Hippel-Lindau tumor suppressor gene results in wide-spread hyperplastic, inflammatory and fibrotic lesions in the kidney. Oncogene 34, 2631–2639 (2015). 10.1038/onc.2014.197

15 Wang, X. et al. Multi-Omics Profiling to Assess Signaling Changes upon VHL Restoration and Identify Putative VHL Substrates in Clear Cell Renal Cell Carcinoma Cell Lines. Cells 11 (2022). 10.3390/cells11030472

16 Du, W. et al. HIF drives lipid deposition and cancer in ccRCC via repression of fatty acid metabolism. Nat Commun 8, 1769 (2017). 10.1038/s41467-017-01965-8

17 Wang, Z. et al. VHL suppresses autophagy and tumor growth through PHD1-dependent Beclin1 hydroxylation. EMBO J 43, 931–955 (2024). 10.1038/s44318-024-00051-2

18 Cho, H. et al. On-target efficacy of a HIF-2alpha antagonist in preclinical kidney cancer models. Nature 539, 107–111 (2016). 10.1038/nature19795

19 Wallace, E. M. et al. A Small-Molecule Antagonist of HIF2alpha Is Efficacious in Preclinical Models of Renal Cell Carcinoma. Cancer Res 76, 5491–5500 (2016). 10.1158/0008-5472.CAN-16-0473

20 Kondo K., K. J., Nakamura E, Lechpammer M, Kaelin WGJr. Inhibition of HIF is necessary for tumor suppression by the von Hippel-Lindau protein. Cancer Cell 1, 237–246 (2002). 10.1016/s1535-6108(02)00043-0.

21 Haferkamp, I. et al. A candidate NAD+ transporter in an intracellular bacterial symbiont related to Chlamydiae. Nature 432, 622–625 (2004). 10.1038/nature03131

22 Stephanie A. Luff, Nestor A. Fernandez, Christopher M. Sturgeon & Ditadi, A. Generation of functionally distinct hemogenic endothelial cell populations from pluripotent stem cells. Experimental Hematology 138, 104587 (2024). 10.1016/j.exphem.2024.104587.

23 Dudley, A. C. & Griffioen, A. W. Pathological angiogenesis: mechanisms and therapeutic strategies. Angiogenesis 26, 313–347 (2023). 10.1007/s10456-023-09876-7

24 Lugano, R., Ramachandran, M. & Dimberg, A. Tumor angiogenesis: causes, consequences, challenges and opportunities. Cell Mol Life Sci 77, 1745–1770 (2020). 10.1007/s00018-019-03351-7

25 Carmeliet, P. & Jain, R. K. Molecular mechanisms and clinical applications of angiogenesis. Nature 473, 298–307 (2011). 10.1038/nature10144

26 Potente, M. & Makinen, T. Vascular heterogeneity and specialization in development and disease. Nat Rev Mol Cell Biol 18, 477–494 (2017). 10.1038/nrm.2017.36

27 Dao, L. et al. Modeling blood-brain barrier formation and cerebral cavernous malformations in human PSC-derived organoids. Cell Stem Cell 31, 818–833 e811 (2024). 10.1016/j.stem.2024.04.019

28 Romeo, S. G. et al. Human blood vessel organoids reveal a critical role for CTGF in maintaining microvascular integrity. Nat Commun 14, 5552 (2023). 10.1038/s41467-023-41326-2

29 Sun, X. Y. et al. Generation of vascularized brain organoids to study neurovascular interactions. Elife 11 (2022). 10.7554/eLife.76707

30 Frenz-Wiessner, S. et al. Generation of complex bone marrow organoids from human induced pluripotent stem cells. Nat Methods 21, 868–881 (2024). 10.1038/s41592-024-02172-2

31 Wimmer, R. A. et al. Human blood vessel organoids as a model of diabetic vasculopathy. Nature 565, 505–510 (2019). 10.1038/s41586-018-0858-8

32 Cakir, B. et al. Engineering of human brain organoids with a functional vascular-like system. Nat Methods 16, 1169–1175 (2019). 10.1038/s41592-019-0586-5

33 Goveia, J. et al. An Integrated Gene Expression Landscape Profiling Approach to Identify Lung Tumor Endothelial Cell Heterogeneity and Angiogenic Candidates. Cancer Cell 37, 21–36 e13 (2020). 10.1016/j.ccell.2019.12.001

34 Pan, Z. et al. Generation of iPSC-derived human venous endothelial cells for the modeling of vascular malformations and drug discovery. Cell Stem Cell 32, 227–245 e229 (2025). 10.1016/j.stem.2024.10.015

35 Okita, K. et al. An efficient nonviral method to generate integration-free human-induced pluripotent stem cells from cord blood and peripheral blood cells. Stem Cells 31, 458–466 (2013). 10.1002/stem.1293

36 Nakagawa, M. et al. A novel efficient feeder-free culture system for the derivation of human induced pluripotent stem cells. Sci Rep 4, 3594 (2014). 10.1038/srep03594

37 Liu, F. et al. Case report: a synonymous VHL mutation (c.414A > G, p.Pro138Pro) causes pathogenic familial hemangioblastoma through dysregulated splicing. BMC Med Genet 21, 42 (2020). 10.1186/s12881-020-0976-7

38 Zbar, B. et al. Germline mutations in the Von Hippel-Lindau disease (VHL) gene in families from North America, Europe, and Japan. Human Mutation 8, 348–357 (1996). 10.1002/(sici)1098-1004(1996)8:4<348::AID-HUMU8>3.0.CO;2-3.

39 James, D. et al. Expansion and maintenance of human embryonic stem cell–derived endothelial cells by TGFβ inhibition is Id1 dependent. Nature Biotechnology 28, 161– 166 (2010). 10.1038/nbt1605

40 Li, J. et al. A transient DMSO treatment increases the differentiation potential of human pluripotent stem cells through the Rb family. PLoS One 13, e0208110 (2018). 10.1371/journal.pone.0208110

41 Hamad, S. et al. High-efficient serum-free differentiation of endothelial cells from human iPS cells. Stem Cell Res Ther 13, 251 (2022). 10.1186/s13287-022-02924-x

42 Patsch, C. et al. Generation of vascular endothelial and smooth muscle cells from human pluripotent stem cells. Nat Cell Biol 17, 994–1003 (2015). 10.1038/ncb3205

43 Ito, H. et al. Localized Gene Expression Analysis during Sprouting Angiogenesis in Mouse Embryoid Bodies Using a Double Barrel Carbon Probe. Anal Chem 88, 610–613 (2016). 10.1021/acs.analchem.5b04338

44 M Mohammadi et al. Structures of the tyrosine kinase domain of fibroblast growth factor receptor in complex with inhibitors. Science 276, 955–960 (1997). 10.1126/science.276.5314.955

45 Choueiri, T. K. et al. Inhibition of hypoxia-inducible factor-2alpha in renal cell carcinoma with belzutifan: a phase 1 trial and biomarker analysis. Nat Med 27, 802–805 (2021). 10.1038/s41591-021-01324-7

46 Xu, R. et al. 3-[(1S,2S,3R)-2,3-Difluoro-1-hydroxy-7-methylsulfonylindan-4-yl]oxy-5-fluorobenzonitrile (PT2977), a Hypoxia-Inducible Factor 2alpha (HIF-2alpha) Inhibitor for the Treatment of Clear Cell Renal Cell Carcinoma. J Med Chem 62, 6876–6893 (2019). 10.1021/acs.jmedchem.9b00719

47 Bess, S. N., Smart, G. K., Igoe, M. J. & Muldoon, T. J. Scaffold-free development of multicellular tumor spheroids with spatial characterization of structure and metabolic radial profiles. In Vitro Model 3, 91–108 (2024). 10.1007/s44164-024-00074-3

48 Iwano, S. et al. Single-cell bioluminescence imaging ofdeep tissue in freely moving animals. Science 359, 935–939 (2018). 10.1126/science.aaq1067

49 Setyawan, N. H., Hartanto, R. A., Malueka, R. G., Dwianingsih, E. K. & Dharma, D. P. Dual manifestations: spinal and cerebellar hemangioblastomas indicative of von Hippel-Lindau syndrome. Radiology Case Reports 19, 5000–5006 (2024). 10.1016/j.radcr.2024.07.158

50 Yoda, R. A. & Cimino, P. J. Neuropathologic features of central nervous system hemangioblastoma. Journal of Pathology and Translational Medicine 56, 115–125 (2022). 10.4132/jptm.2022.04.13

51 Kim, Y. H. et al. A Case of Von Hippel-Lindau Disease Presented with Multiple Pancreatic Cysts and Medullary Hemangioblastoma. Clinical Pediatric Hematology-Oncology 27, 67–71 (2020). 10.15264/cpho.2020.27.1.67

52 Myong, N. H. & Park, B. J. Malignant glioma arising at the site of an excised cerebellar hemangioblastoma after irradiation in a von Hippel-Lindau disease patient. Yonsei Med J 50, 576–581 (2009). 10.3349/ymj.2009.50.4.576

53 Arai, Y. et al. Patient-derived and gene-edited pluripotent stem cells lacking NPHP1 recapitulate juvenile nephronophthisis in abnormalities of primary cilia and renal cyst formation. Front Cell Dev Biol 12, 1370723 (2024). 10.3389/fcell.2024.1370723

54 Harlander, S. et al. Combined mutation in Vhl, Trp53 and Rb1 causes clear cell renal cell carcinoma in mice. Nat Med 23, 869–877 (2017). 10.1038/nm.4343

55 Lolkema, M. P. et al. Allele-specific regulation of primary cilia function by the von Hippel-Lindau tumor suppressor. Eur J Hum Genet 16, 73–78 (2008). 10.1038/sj.ejhg.5201930

56 Bahar, M. E., Kim, H. J. & Kim, D. R. Targeting the RAS/RAF/MAPK pathway for cancer therapy: from mechanism to clinical studies. Signal Transduct Target Ther 8, 455 (2023). 10.1038/s41392-023-01705-z

57 Ang, L. T. et al. Generating human artery and vein cells from pluripotent stem cells highlights the arterial tropism of Nipah and Hendra viruses. Cell 185, 2523–2541 e2530 (2022). 10.1016/j.cell.2022.05.024

58 Sissaoui, S. et al. Genomic Characterization of Endothelial Enhancers Reveals a Multifunctional Role for NR2F2 in Regulation of Arteriovenous Gene Expression. Circ Res 126, 875–888 (2020). 10.1161/CIRCRESAHA.119.316075

59 Chen, X., Qin, J., Cheng, C. M., Tsai, M. J. & Tsai, S. Y. COUP-TFII is a major regulator of cell cycle and Notch signaling pathways. Mol Endocrinol 26, 1268–1277 (2012). 10.1210/me.2011-1305

60 Perelli, L. et al. Interferon signaling promotes tolerance to chromosomal instability during metastatic evolution in renal cancer. Nat Cancer 4, 984–1000 (2023). 10.1038/s43018-023-00584-1

61 Tabaro, F. et al. VHLdb: A database of von Hippel-Lindau protein interactors and mutations. Sci Rep 6, 31128 (2016). 10.1038/srep31128

62 Gossage, L., Eisen, T. & Maher, E. R. VHL, the story of a tumour suppressor gene. Nat Rev Cancer 15, 55–64 (2015). 10.1038/nrc3844

63 Zhou, Y. et al. Metascape provides a biologist-oriented resource for the analysis of systems-level datasets. Nat Commun 10, 1523 (2019). 10.1038/s41467-019-09234-6

64 Nikolova, M. T. et al. Fate and state transitions during human blood vessel organoid development. Cell 188, 3329–3348 e3331 (2025). 10.1016/j.cell.2025.03.037

65 Piera-Velazquez, S. & Jimenez, S. A. Endothelial to Mesenchymal Transition: Role in Physiology and in the Pathogenesis of Human Diseases. Physiol Rev 99, 1281–1324 (2019). 10.1152/physrev.00021.2018

66 Douglas J. Williamson, S. B.-M., James Degregori, AND H. Earl Ruley. hnRNP C Is Required for Postimplantation Mouse Development but Is Dispensable for Cell Viability. MOLECULAR AND CELLULAR BIOLOGY 20, 4094–4105 (2000). 10.1128/MCB.20.11.4094-4105.

67 Pont, A. R., Sadri, N., Hsiao, S. J., Smith, S. & Schneider, R. J. mRNA decay factor AUF1 maintains normal aging, telomere maintenance, and suppression of senescence by activation of telomerase transcription. Mol Cell 47, 5–15 (2012). 10.1016/j.molcel.2012.04.019

68 Serrano, S. L. a. M. Depletion of ribosomal protein L37 occurs in response to DNA damage and activates p53 through the L11/MDM2 pathway. cell cycle, 4005–4012 (2010).

69 Xu, H., Kita, Y., Bang, U., Gee, P. & Hotta, A. Optimized electroporation of CRISPR-Cas9/gRNA ribonucleoprotein complex for selection-free homologous recombination in human pluripotent stem cells. STAR Protoc 2, 100965 (2021). 10.1016/j.xpro.2021.100965

70 Liu, Y. Y. et al. HNRNPC mediates lymphatic metastasis of cervical cancer through m6A-dependent alternative splicing of FOXM1. Cell Death Dis 15, 732 (2024). 10.1038/s41419-024-07108-4

71 Yang, N. et al. hnRNPC Promotes Malignancy in Pancreatic Cancer through Stabilization of IQGAP3. Biomed Res Int 2022, 6319685 (2022). 10.1155/2022/6319685

72 Hu, J., Cai, D., Zhao, Z., Zhong, G. C. & Gong, J. Suppression of Heterogeneous Nuclear Ribonucleoprotein C Inhibit Hepatocellular Carcinoma Proliferation, Migration, and Invasion via Ras/MAPK Signaling Pathway. Front Oncol 11, 659676 (2021). 10.3389/fonc.2021.659676

73 Liu, D. et al. HNRNPC downregulation inhibits IL-6/STAT3-mediated HCC metastasis by decreasing HIF1A expression. Cancer Sci 113, 3347–3361 (2022). 10.1111/cas.15494

74 Guo, W. et al. Elevated Heterogeneous Nuclear Ribonucleoprotein C Expression Correlates With Poor Prognosis in Patients With Surgically Resected Lung Adenocarcinoma. Front Oncol 10, 598437 (2020). 10.3389/fonc.2020.598437

75 Lai, Y., Song, M., Hakala, K., Weintraub, S. T. & Shiio, Y. Proteomic dissection of the von Hippel-Lindau (VHL) interactome. J Proteome Res 10, 5175–5182 (2011). 10.1021/pr200642c

76 Wu, J. et al. Essential m(6)A Methylation Regulator HNRNPC Serves as a Targetable Biomarker for Papillary Renal Cell Carcinoma. J Oncol 2022, 9411692 (2022). 10.1155/2022/9411692

77 Park, D. M. et al. von Hippel-Lindau disease-associated hemangioblastomas are derived from embryologic multipotent cells. PLoS Med 4, e60 (2007). 10.1371/journal.pmed.0040060

78 Zhuang, Z. et al. Tumor derived vasculogenesis in von Hippel-Lindau disease-associated tumors. Sci Rep 4, 4102 (2014). 10.1038/srep04102

79 Ohh, M. et al. The von Hippel-Lindau Tumor Suppressor Protein Is Required for Proper Assembly of an Extracellular Fibronectin Matrix. Molecular Cell 1, 959–968 (1998). 10.1016/s1097-2765(00)80096-9.

80 Lutz, M. S. & Burk, R. D. Primary cilium formation requires von hippel-lindau gene function in renal-derived cells. Cancer Res 66, 6903–6907 (2006). 10.1158/0008-5472.CAN-06-0501

81 Frew, I. J. et al. pVHL and PTEN tumour suppressor proteins cooperatively suppress kidney cyst formation. EMBO J 27, 1747–1757 (2008). 10.1038/emboj.2008.96

82 Mo, L. et al. An analysis of the role of HnRNP C dysregulation in cancers. Biomark Res 10, 19 (2022). 10.1186/s40364-022-00366-4

83 Kreitzer, F. R. et al. A robust method to derive functional neural crest cells from human pluripotent stem cells. American Journal of Stem Cells 2, 119–131 (2013).

84 Hayashi, Y. et al. Structure-based discovery of NANOG variant with enhanced properties to promote self-renewal and reprogramming of pluripotent stem cells. Proc Natl Acad Sci U S A 112, 4666–4671 (2015). 10.1073/pnas.1502855112

85 Li, R. et al. Generation of human induced pluripotent stem cell lines derived from two glucose transporter 1 deficiency syndrome patients. Stem Cell Res 81, 103584 (2024). 10.1016/j.scr.2024.103584

86 Mori, M. et al. Generation of human induced pluripotent stem cell lines derived from four Rett syndrome patients with MECP2 mutations. Stem Cell Res 77, 103432 (2024). 10.1016/j.scr.2024.103432

87 Miyawaki, S., Okada, Y., Okano, H. & Miura, K. Teratoma Formation Assay for Assessing Pluripotency and Tumorigenicity of Pluripotent Stem Cells. Bio Protoc 7, e2518 (2017). 10.21769/BioProtoc.2518

88 Nelakanti, R. V., Kooreman, N. G. & Wu, J. C. Teratoma formation: a tool for monitoring pluripotency in stem cell research. Curr Protoc Stem Cell Biol 32, 4A 8 1–4A 8 17 (2015). 10.1002/9780470151808.sc04a08s32

89 Hao, Y. et al. Dictionary learning for integrative, multimodal and scalable single-cell analysis. Nat Biotechnol 42, 293–304 (2024). 10.1038/s41587-023-01767-y

