## Supplementary Materials for "HNRNPC-mediated RNA processing promotes VHL-associated excessive vasculogenesis in von Hippel-Lindau syndrome revealed from patient-derived blood vessel models"

Table of contents:
Supplementary Material, Table 1-3

**Supplementary Table S1**

The list of primer sets, crRNA sequences, and Taqman probes used in this study.

| **Primers** | **Primer sequence (5'→3')** |
|---|---|
| Piggy back（Venus-Akaluc） | F: caccatggtgagcaagggcgaggag<br>R: ttacacggcgatcttgccgtccttc |
| Exon1 in *VHL* | F: agcgcgttccatcctctaccga<br>R: gtagaggggcttcagaccgtgct |
| Exon2 in *VHL* | F: cagccaccggtgtggctctttaac<br>R: tctcctggaactccagggcacaa |
| Exon3 in *VHL* | F: tgagaccctagtctgccactgagg<br>R: agcaaaaatgccaccaccttctcct |
| Full length in *VHL* cDNA | F: tctggatcgcggagggaat<br>R: aaatcttcaatctcccatccgttg |
| KO check primers in *EPAS1* | F: ctcttttcgggtctgacagc<br>R: gtcgcagggatgagtgaagt |
| KO check primers in *HNRNPC* | F: ataagggctttgccttcgtt<br>R:ccctcctcagcagagtcatc |
| *EBNA1* | F: atcagggccaagacatagagatg<br>R: gccaatgcaacttggacgtt |
| *GAPDH* | F: gagccacatcgctcagacac<br>R: ttgccatgggtggaatcata |
| *Nested-PCR, 1st step PCR (MCGpF11/MCGpR1)* | F: acaccatgggag(c/t)tggtaat<br>R: cttc(a/t)tcgactt(c/t)cagacccaaggcat |
| *Nested-PCR, 2nd step PCR (R16–2/MCGpR21)* | F: gtg(c/g)gg(a/c)tggatcacctcct<br>R: gcatccacca(a/t)a(a/t)ac(c/t)ctt |

| **crRNA** | **crRNA sequence (5'→3')** |
|---|---|
| *VHL* | catacgggcagcacgacgcg |
| *EPAS1* | crRNA1: agaagtcccgggatgctgcg<br>crRNA2: cctccgtctccttgctccgc |

| | |
|---|---|
| *HNRNPC* | ctcattaacatactgaacga |

| Taqman probes | **Taqman Assay ID** |
|---|---|
| *GAPDH* | Hs02786624_g1 |
| *VHL(EXON1)* | Hs03046964_s1 |
| *VHL(EXON2-3)* | Hs00184451_m1 |
| *HIF1A* | Hs00153153_m1 |
| *EPAS1(HIF2A)* | Hs01026149_m1 |
| *NANOG* | Hs02387400_g1 |
| *PECAM1* | Hs01065279_m1 |
| *CDH5* | Hs00901465_m1 |
| *DLL4* | Hs00184092_m1 |
| *NR2F2* | Hs00819630_m1 |
| *APLN* | Hs00175572_m1 |
| *ESM1* | Hs00199831_m1 |
| *MYC* | Hs00153408_m1 |
| *DUSP4* | Hs01027785_m1 |

**Supplementary Table S2**

The list of primary antibodies used in this study.

| Name | Host species | Dilutions or concentrations | Source | Identifier |
|---|---|---|---|---|
| GAPDH | Mouse | 1:1000 | R&D systems | Cat#MAB5718 |
| VHL | Rabbit | 1:50 or 1:200 | Cell signaling | Cat#68547 |
| HIF1A | Mouse | 1:50 | R&D systems | Cat#MAB1536 |
| HIF2A | Rabbit | 1:50 | Cell signaling | Cat#71565S |
| CD31 | Mouse | 1:100 | BioLegend | Cat#303102 |
| CD31 | Sheep | 1:100 or 0.4μg/mL | R&D systems | Cat#AF806 |
| APC anti-human CD31 | Mouse | 1:100 | BioLegend | Cat#303116 |
| FITC anti-human CD144 | Mouse | 1:100 | BD bioscience | Cat#560874 |
| NG2 | Rabbit | 1:100 | Sigma | Cat#AB5320 |
| NANOG | Rabbit | 1:500 | Reprocell | Cat#RCAB004P-F |
| OCT3/4 | Mouse | 1:200 | Santa Cruz | Cat#sc-5279 |
| HNRNPC | Rabbit | 1:50 | Abcam | Cat#ab133607 |
| ARL13B | Rabbit | 1:1000 | Proteintech | Cat#17711-1-AP |
| Alexa Fluor 488 anti-human TRA-1-60 | Mouse | 1:100 | BioLegend | Cat#330613 |
| APC anti-human SSEA4 | Mouse | 1:100 | BioLegend | Cat#330418 |
| TUJ1 | Mouse | 1:250 | R&D systems | Cat#MAB1195 |
| Alpha-Fetoprotein(AFP) | Mouse | 1:200 | R&D systems | Cat#MAB1368 |

| | | | | |
|---|---|---|---|---|
| αSMA | Mouse | 1:250 | R&D systems | Cat#MAB1420 |
| Inhibin alpha | Mouse | 2μg/mL | abcam | Cat#ab14087 |
| PE anti-human CD140b | Mouse | 1:100 | BD Bioscience | Cat#558821 |
| PE anti-human CD45 | Mouse | 1:100 | BD Bioscience | Cat#555483 |
| GFP | Rabbit | 1μg/mL | Invitrogen | Cat#A-11122 |

**Supplementary Table S3**

The list of secondary antibodies used in this study.

| **Name** | **Host species** | **Dilutions or concentrations** | **Source** | **Identifier** |
|---|---|---|---|---|
| Anti-Mouse IgG（H+L) Highly Cross-Absorbed Secondary Antibody, Alexa Fluor Plus 488 | Donkey | 1:500 | Thermo Fisher Scientific | Cat#A-21202 |
| Anti-Mouse IgG（H+L) Highly Cross-Absorbed Secondary Antibody, Alexa Fluor Plus 555 | Donkey | 1:500 | Thermo Fisher Scientific | Cat#A-31570 |
| Anti-sheep IgG （H+L) Cross-Absorbed Secondary Antibody, Alexa Fluor 647 | Donkey | 1:500 | Invitrogen | Cat#A-21448 |
| DyLight™ 488 Donkey anti-rabbit IgG (minimal x-reactivity) Antibody | Donkey | 1:500 | Biolegend | Cat#406404 |

| | | | | |
|---|---|---|---|---|
| DyLight™ 649 Donkey anti-rabbit IgG (minimal x-reactivity) Antibody | Donkey | 1:500 | Biolegend | Cat#406406 |
| Cyanine3 Donkey anti-rabbit IgG (minimal x-reactivity) Antibody | Donkey | 1:500 | Biolegend | Cat#406402 |
| Anti-Mouse Secondary HRP Antibody | - | - | ProteinSimple | Cat#042-205 |
| Anti-Rabbit Secondary HRP Antibody | - | - | ProteinSimple | Cat#042-206 |
